# Mutant-selective Degradation by BRAF-targeting PROTACs

**DOI:** 10.1101/2020.08.10.245159

**Authors:** Shanique Alabi, Saul Jaime-Figueroa, Zhan Yao, Yijun Gao, John Hines, Kusal T.G. Samarasinghe, Lea Vogt, Neal Rosen, Craig M. Crews

## Abstract

Over 300 BRAF missense mutations have been identified in patients, yet currently approved drugs target V600 mutants alone. Moreover, acquired resistance inevitably emerges, primarily due to RAF lesions that prevent inhibition of BRAF V600 with current treatments. Therefore, there is a need for new therapies that target other mechanisms of activated BRAF. In this study, we use the Proteolysis Targeting Chimera (PROTAC) technology, which promotes ubiquitination and degradation of neo-substrates, to address the limitations of BRAF inhibitor-based therapies. Using vemurafenib-based PROTACs, we successfully achieve sub-nanomolar degradation of all classes of BRAF mutants, but spare degradation of WT RAF family members. Our lead PROTAC outperforms vemurafenib in inhibiting cancer cell growth and shows *in vivo* efficacy in a Class 2 BRAF xenograft model. Mechanistic studies reveal that BRAF^WT^ is spared due to weak ternary complex formation in cells owing to its quiescent inactivated conformation, and activation of BRAF^WT^ sensitizes it to degradation. This study highlights the degree of selectivity achievable using degradation-based therapies by targeting mutant BRAF-driven cancers while sparing BRAF^WT^ and thus expanding the therapeutic window using a new anti-tumor drug modality.

## Main

The Ras-RAF-MEK-ERK pathway is important for many aspects of cellular homeostasis^1^. The pathway is initiated upon extracellular growth factor binding to receptor tyrosine kinases (RTKs), thereby activating the kinase cascade^2^. Upon activation of upstream effectors, GTP-bound RAS recruits RAF (ARAF, BRAF or CRAF) to the cell membrane, promoting its dimerization and activation^2^. Thus, the scaffolding and enzymatic role of BRAF are both essential for its function^3–5^. Activated RAF phosphorylates and activates MEK, which in turn phosphorylates and activates ERK leading to cell proliferation, differentiation, and survival^2^. BRAF is mutated in 8% of observed tumors including melanoma (60%)^6^, colorectal cancer (10%)^7^, non-small cell lung cancer (NSCLC) (10%)^8^ and hairy cell leukemia (100%)^9^. These mutations (often missense mutations found in the kinase domain) distinctly affect the biochemical characteristics of the kinase^10–12^. Class 1 BRAF mutants such as V600E and V600K are hyper-activating and can signal as monomers in the absence of activated RAS^13^. Class 2 BRAF mutants such as K601E and G469A signal as constitutive, RAS-independent dimers^14^. Lastly, Class 3 BRAF mutants such as G466V and D594N harbor low to no kinase activity and function by binding tightly to RAS thus recruiting CRAF into hyperactivated heterodimers ^15–17^. FDA-approved inhibitors such as vemurafenib have been successful in increasing progression-free survival of patients harboring hyperactive BRAF V600E mutations^18^. However, as with many kinase inhibitors, resistance occurs that renders patients insensitive to continued treatment ^19^. Significant efforts have focused on creating drugs that target Class 2 BRAF mutations by inhibiting dimer formation, but adequate drugs have not yet been approved^10^. Therefore, there is a need for new and innovative therapies to address BRAF-driven cancers.

Proteolysis Targeting Chimeras (PROTACs) are heterobifunctional small molecules composed of a warhead that binds a protein of interest (POI), a flexible linker, and a ligand that binds an E3 ligase^20, 21^. These molecules recruit an E3 ligase (*e.g*. VHL) to a POI to form a ternary complex. Upon complex formation, ubiquitin molecules are transferred to accessible lysines on the POI, marking it for proteasomal degradation. Importantly, by eliminating the entire protein scaffold, PROTACs are able to target both the enzymatic and non-enzymatic roles of disease-causing proteins. In recent years, our lab and others have made considerable progress in using PROTAC technology to induce degradation of proteins involved in disease, such as AR, ER, BRD4, RIPK2, BCR-Abl, EGFR, MET, p38 MAPK, BTK, and ERRα ^22–32^ . While traditional inhibitors require sustained target engagement for therapeutic effect, PROTACs simply require transient interaction, offering the ability to degrade proteins with limited target engagement^22^. Furthermore, the modular design of PROTACs allows for additional selectivity to be tuned into the small molecule, making it ideal for addressing difficult targets such as BRAF.

## Results

### SJF-0628 induces efficient and potent degradation of mutant BRAF but spares BRAF^WT^

Although the utility of vemurafenib is limited to the treatment of tumors driven by BRAF^V600^ mutations, biochemical and binding studies show that these inhibitors also interact with BRAF^WT^, Class 2, and Class 3 BRAF mutants^11, 15, 18, 33^. We therefore hypothesized that all BRAF isoforms would be susceptible to degradation by a vemurafenib-based PROTAC. Crystal structures of vemurafenib bound to BRAF^V600E^ reveal a solvent-exposed chloride at the *para*-position on the phenyl ring, which we posited would be ideal for linker addition (PDB: 3OG7) ^34^ (Extended Data Fig.1a-b). Pursuant to this, we iteratively optimized a lead vemurafenib-based PROTAC, SJF-0628, by coupling vemurafenib to a ligand for the von Hippel Lindau (VHL) E3 ligase using a rigid piperazine linker (Fig. 1a). In addition, we synthesized a degradation-incompetent control, **SJF-0661**, by inverting the stereocenter of the critical hydroxyl-proline group in the VHL ligand ^22, 35^. In NIH3T3 cells expressing doxycycline-inducible^14, 15^, V5-tagged BRAF^WT^, Class 1, 2, or 3 BRAF mutations, SJF-0628 caused a dose-dependent decrease in the expression of all tested BRAF mutants, but spared BRAF^WT^, ARAF, and CRAF (Fig. 1b, Extended Data Fig.1c). Mutant selectivity was also observed in T-Rex 293 cells expressing HA-tagged BRAF isoforms (Extended Data Fig.1d).

**Figure 1.**
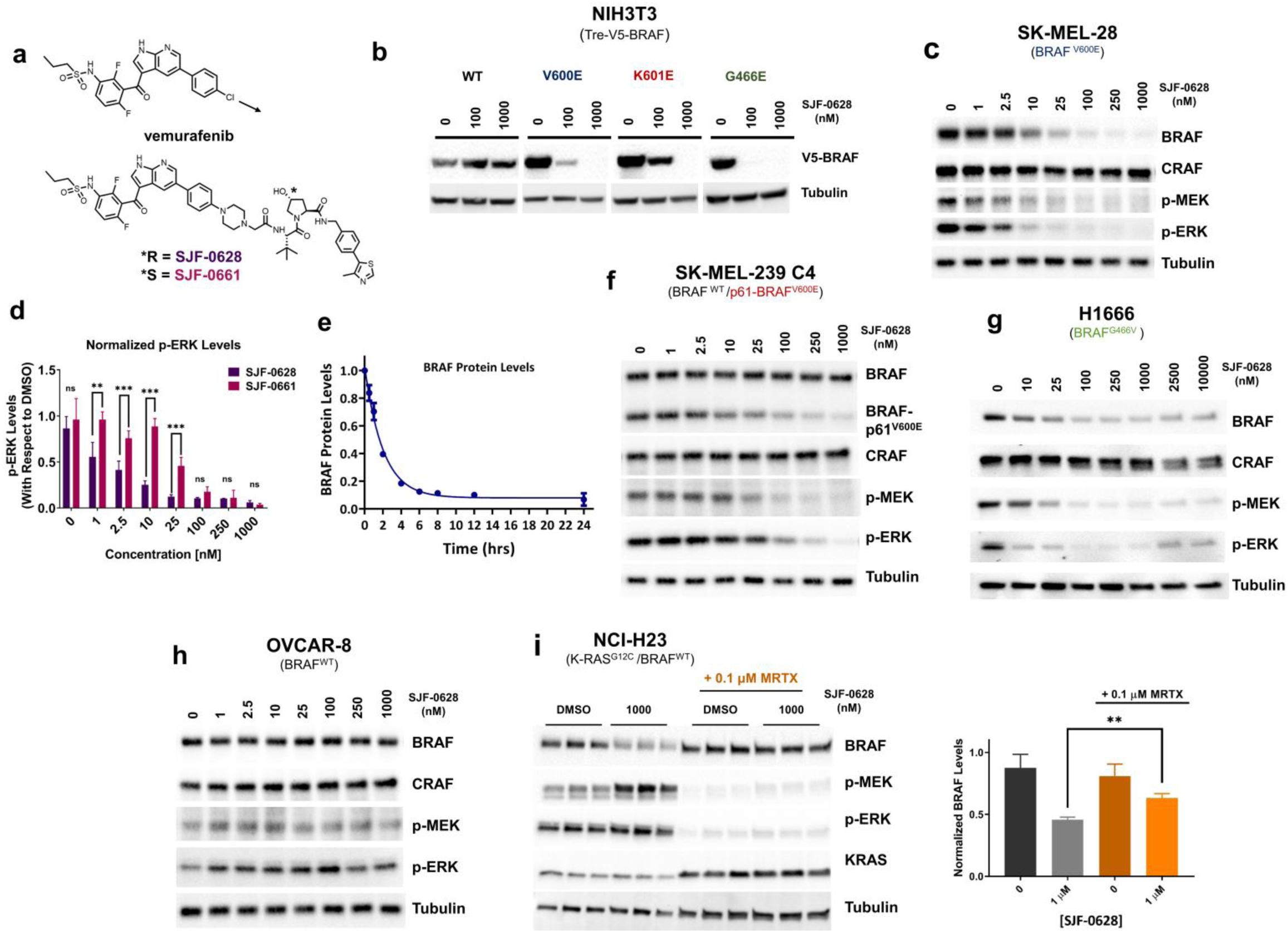
Vemurafenib-based PROTAC SJF-0628 potently, selectively, and efficiently induces degradation of mutant BRAF. **a**, Chemical structure of vemurafenib and BRAF targeting PROTAC, SJF-0628, and its epimer, SJF-0661. SJF-0628 is composed of vemurafenib, a short piperazine-based linker, and a VHL recruiting ligand. SJF-0661 has an identical warhead and linker as SJF-0628 but contains an inverted hydroxyl group in the VHL ligand and is therefore unable to engage VHL to induce ubiquitination. **b**, Inducible NIH3T3 cells expressing indicated V5-BRAF constructs (doxycycline 100-200 ng/mL, 24 hours) treated with increasing amounts of SJF-0628. **c**, SK-MEL-28 cells (homozygous BRAF^V600E^) treated with indicated amounts of SJF-0628 induced BRAF degradation and suppression of MEK and ERK phosphorylation. **d**, Quantitation of ERK inhibition in SK-MEL-28 cells treated with SJF-0628 or SJF-0661 (mean ± s.d., n=3) *** *P* value < 0.001. **e**, Quantitation of SJF-0628 treatment time course (100 nM) at indicated times in SK-MEL-28 cells shows maximal degradation within 4 hours (mean ± s.d., n=2). **f**, SJF-0628 induces selective degradation of p61-BRAF^V600E^ mutant and inhibits MEK and ERK phosphorylation but spares BRAF^WT^ and CRAF in SK-MEL-239-C4 cells. **g**, H1666 (heterozygous BRAF^G466V^) treated with SJF-0628 shows BRAF degradation, but incomplete suppression of ERK signaling. **h**, BRAF^WT^ is spared by SJF-0628 in OVCAR-8 cells but induces slight activation of ERK phosphorylation. **i**, Covalent inhibition of KRAS^G12C^ by MRTX849 in H23 cells hinders PROTAC induced BRAF^WT^ degradation ((mean ± s.d., n=3, ** P value < 0.01). *P* value calculated by unpaired t-test.

To confirm these findings, we evaluated the ability of SJF-0628 to degrade endogenously expressed BRAF mutants in cancer cells. SJF-0628 treatment of SK-MEL-28 cells (homozygous BRAF^V600E^) resulted in a DC_50_ (half-maximal degradation) value of 6.8 nM and D_MAX_ (percent of maximal degradation) of > 95% (Fig. 1c); similar results were seen in A375 cells (homozygous BRAF^V600E^) (Extended Data Fig. 2a). In SK-MEL-239 cells (heterozygous BRAF^V600E^), minimal BRAF degradation was observed, likely due to residual BRAF^WT^ although, there is a similarly sustained decrease in MAPK signaling (Extended Data Fig. 2b).

Treatment with 10 nM SJF-0628 caused maximal inhibition of MEK and ERK phosphorylation in SK-MEL-28 cells (Fig. 1c). As expected, while the epimer control, SJF-0661, did not decrease BRAF protein levels (Extended Data Fig. 2c), inhibition of MEK and ERK phosphorylation by this vemurafenib-based molecule was nonetheless observed. However, maximal suppression of p-ERK required 100 nM of SJF-0661, affirming a 10-fold increase in potency for the PROTAC from targeting both the enzymatic and non-enzymatic roles of BRAF (Fig. 1d). SJF-0628 induced near complete BRAF^V600E^ degradation within 4 hours (Fig. 1e, Extended Data Fig. 2d), and BRAF^V600E^ degradation and p-ERK inhibition was sustained for up to 72 hours (Extended Data Fig. 2e). A wash-out experiment of SJF-0628 after a 24 hour treatment showed 30% recovery of BRAF levels and MAPK phosphorylation after 24 hours, confirming the long-acting and possibly catalytic effect of PROTACs (Extended Data Fig. 2f), BRAF^V600E^ degradation was prevented when cells were pre-treated with epoxomicin (proteasome inhibitor)^36^, MLN-4924 (neddylation inhibitor)^37^ or 100-fold excess vemurafenib, confirming SJF-0628 mediated protein loss is consistent with a PROTAC mechanism of action (Extended Data Fig. 2g). Treatment of VHL ligand alone did not affect MAPK phosphorylation, showing that the effect of the PROTAC is primarily due to BRAF degradation (Extended Data Fig. 2h)

One clinically observed acquired resistance mechanism to vemurafenib is the aberrantly spliced BRAF mRNA transcript encoding an N-terminally truncated isoform that signals as a constitutive dimer (BRAF-p61^V600E^)^38^. In SK-MEL-239 C4 cells (BRAF^WT^/ BRAF-p61^V600E^)^38^, SJF-0628 induced the degradation of the p61 dimer with a DC_50_ of 72 nM and D_MAX_ >80%, while notably sparing BRAF^WT^ and CRAF (Fig. 1f). Similar results were seen in HCC-364 vr1cells (BRAF^WT^/ BRAF-p61^V600E^) ^39^ (DC_50_ of 147 nM, D_MAX_ >90%) and 293 T-Rex cells overexpressing HA-BRAF-p61^V600E^ (Extended Data Fig. 3a,b). In t SK-MEL-246 cancer cells^40^ (Class 2, BRAF^G469A^), SJF-0628 induced dose-dependent degradation of BRAF (DC_50_ = 15 nM, D_MAX_ >95%) and concomitant inhibition of ERK phosphorylation while CRAF is slightly stabilized. (Extended Data Fig. 3c).

Class 3 BRAF mutants are kinase dead or hypoactive and are frequently observed in NSCLC^15, 41^. Unlike inhibitors, PROTACs offer a way to target the non-enzymatic/scaffolding role of these BRAF mutants by promoting their degradation. Treatment of NSCLC cell lines H1666 and CAL-12-T cells (Class 3, BRAF^G466V^; heterozygous and homozygous, respectively) with SJF-0628 caused a dose-dependent loss in BRAF protein levels in both cell lines (CAL-12T: DC_50_ = 23 nM, D_MAX_ >90 %) (H1666 cells: DC_50_ = 29 nM, D_MAX_ >80 %) (Fig. 1g, Extended Data Fig. 3d) as well as substantial p-ERK inhibition but showed slight stabilization at SFJ-0628 concentrations higher than 1 µM.

### BRAF^WT^ activation via upstream effectors sensitizes it to SJF-0628 induced degradation at high concentrations

We next asked whether BRAF^WT^ is spared from SJF-0628-induced degradation in cancer cells as observed in the NIH3T3 overexpression system. In the ovarian carcinoma cell line OVCAR-8, SJF-0628 similarly induced no BRAF^WT^ degradation and a slight induction of p-ERK. (Fig. 1h). Given that activation shifts BRAF^WT^ from a closed (extended contact with N terminus) to an open conformation^42–44^, we sought to determine whether this conformational change can affect BRAF^WT^ susceptibility to SJF-0628 in cells with either amplified receptor tyrosine kinases (RTKs) or mutant RAS. In contrast to cells lacking constitutive upstream signaling, we observed ∼30% degradation of BRAF^WT^ in A-431 cells (HER1 amplification) and ∼ 50% degradation of BRAF^WT^ in SK-BR-3 cells (HER2 amplification) at PROTAC concentrations greater than 1µM (Extended Data Fig. 4a, 8c-left panel). Despite the limited BRAF^WT^ degradation, we still observe paradoxical activation of MAPK signaling (increased p-ERK levels), likely due to PROTAC engagement of residual BRAF^WT^ and/or CRAF. Furthermore, addition of EGF to stimulate the MAPK pathway in OVCAR8 cells sensitized BRAF^WT^ to PROTAC-induced degradation (Extended Data Fig. 4b). In cells with mutant RAS (HCT-116, NCI-H23, and SK-MEL-30), the PROTAC also reduced BRAF^WT^ protein levels by 50-60% and caused ERK activation (Extended Data Fig. 4c,d,e). In H23 cells, reduction in BRAF expression was not accompanied by a change in its mRNA, suggesting that its degradation is induced by SJF-0628 (Extended Data Fig. 4f).

Our results suggest that the activated conformation of BRAF may be sensitized to PROTAC-induced degradation. Accordingly, we tested whether inhibition of upstream signaling in these cells, which would reduce BRAF activation, would also reduce the effects of the PROTAC. In SK-BR-3 cells, the HER2/EGFR kinase inhibitor lapatinib (2-hour pre-treatment) reduced SJF-0628-dependent BRAF^WT^ degradation from 48% to 10% (Extended Data Fig. 4f). Similarly, in NCI-H23 cells, pre-treatment with the K-RAS^G12C^ inhibitor, MRTX849, for 2 hours also desensitized BRAF^WT^ to the PROTAC: from 50% BRAF^WT^ degradation in cells treated with the PROTAC alone to 20% degradation in the MRTX849 pre-treated cells (Fig. 1i). Thus, sensitivity of BRAF^WT^ to SJF-0628-mediated degradation is associated with activation of its upstream effectors.

### Exploration of SJF-0628 mutant selectivity shows BRAF^WT^ is unable to form a stable ternary complex *in cellulo*

The selectivity of SJF-0628 for mutant BRAF over BRAF^WT^ suggests that it may have little on-target toxicity and therefore a wide therapeutic index in patients. However, the mechanism of this selectivity is not clear since vemurafenib binds BRAF^WT^ as well as BRAF mutants. During lead optimization, several vemurafenib-based PROTACs were synthesized with varied linker lengths and composition. Similar to SJF-0628, these PROTACs selectively induced degradation of mutant BRAF (Extended Data Fig. 5a,b). Furthermore, during the preparation of this manuscript, Han *et al*. published cereblon-PROTACs, which incorporated vemurafenib or BI882370, that induces BRAF^V600E^ degradation and also spared BRAF^WT^ ^45^. As this phenomenon appears to hold true for multiple BRAF-targeting PROTACs, we explored the mechanism that underlies the observed selectivity.

We evaluated the distinct mechanistic steps that PROTACs undertake to induce degradation: target engagement, ternary complex formation (Target protein: PROTAC: E3 ligase) and target ubiquitination. In a radioactive *in vitro* assay of purified RAF kinase activity, SJF-0628 potently inhibited both BRAF^WT^ (IC_50_ = 5.8 nM) and BRAF^V600E^ (IC_50_ = 1.87 nM). (Fig. 2a; Table 1) Generally, Class 2 mutants bound SJF-0628 with weaker affinity, but nevertheless are successfully degraded in cells; Class 3 mutants were not tested due to their inherent weak kinase activity. SJF-0628 also induces paradoxical activation, showing that BRAF^WT^ engagement is also achieved in cells. Thus, binary binding is not the basis of isoform degradation selectivity by SJF-0628.

**Figure 2.**
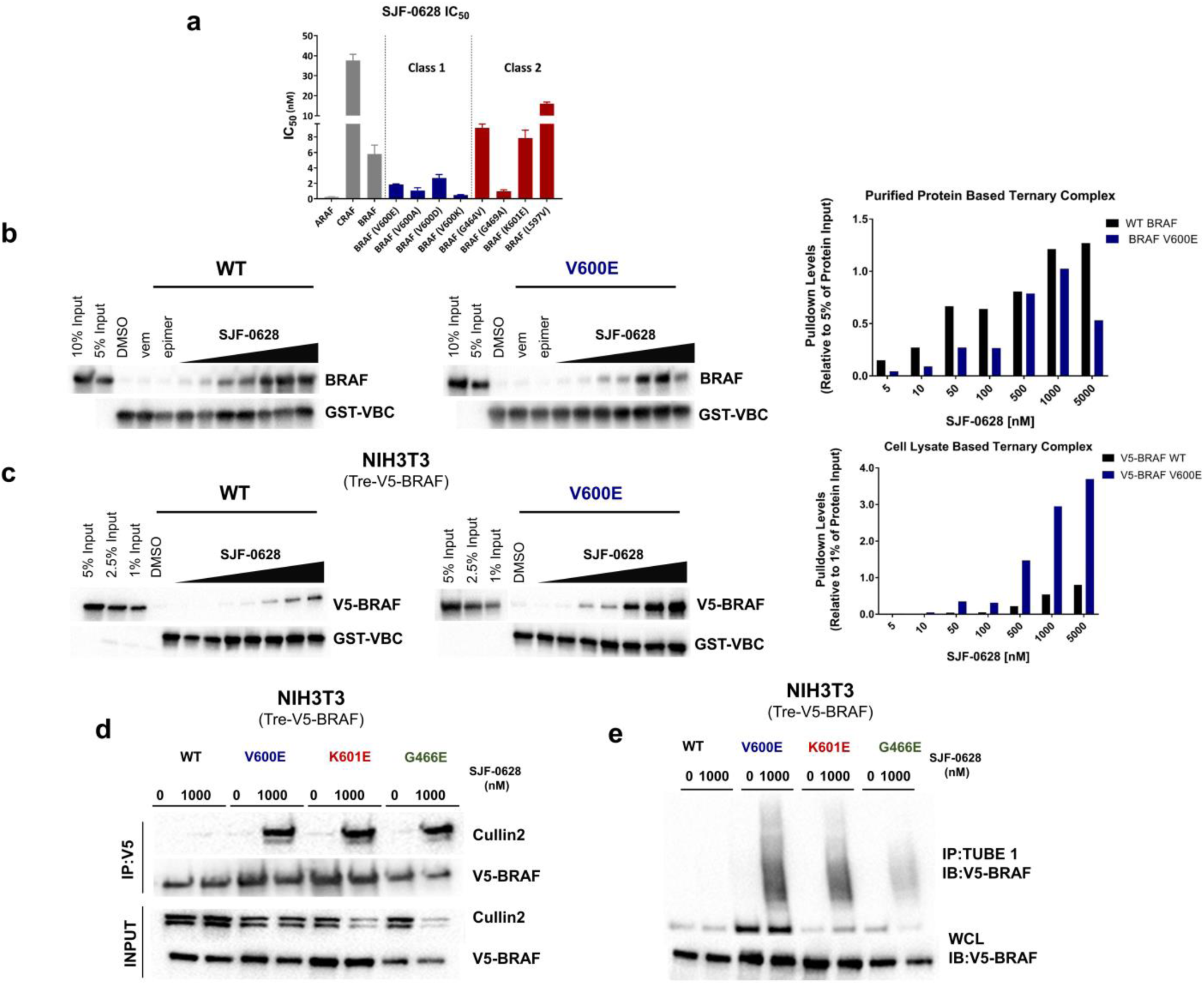
BRAF^WT^ is unable to form a PROTAC-induced ternary complex in cells and thus not degraded. **a**, IC_50_ values of radiolabeled kinase assay for WT RAF and Class 1 and 2 BRAF mutants (mean ± s.d., n=2). Plotted values shown in Table 1. **b**, Purified protein ternary complex assay. GST-VBC (VHL, Elongin B, Elongin C) is immobilized on glutathione beads and incubated with DMSO, vemurafenib (500 nM), SJF-0661 (500 nM) or increasing concentrations of SJF-0628 and purified full length-BRAF to observe VBC:PROTAC:BRAF ternary complex. Quantified with respect to 5% input. **c**, Cell lysate based ternary complex assay (as described in **b**) but using NIH3T3 cell lysates (doxycycline 800 ng/mL) containing V5-BRAF^WT^ or V5-BRAF^V600E^ as input. Quantified with respect to 1% input. **d**, NIH3T3 cells expressing indicated V5-BRAF treated with DMSO or 1µM SJF-0628 for 1-hour followed by immunoprecipitation of V5-BRAF. **e,** Tandem Ubiquitin Binding Entities 1 (TUBE1) pull down of tetra-ubiquitinated proteins in NIH3T3 cells expressing indicated V5-BRAF after 1-hour treatment with vehicle and SJF-0628. Immunoblotted for V5-BRAF.

**Table 1.**
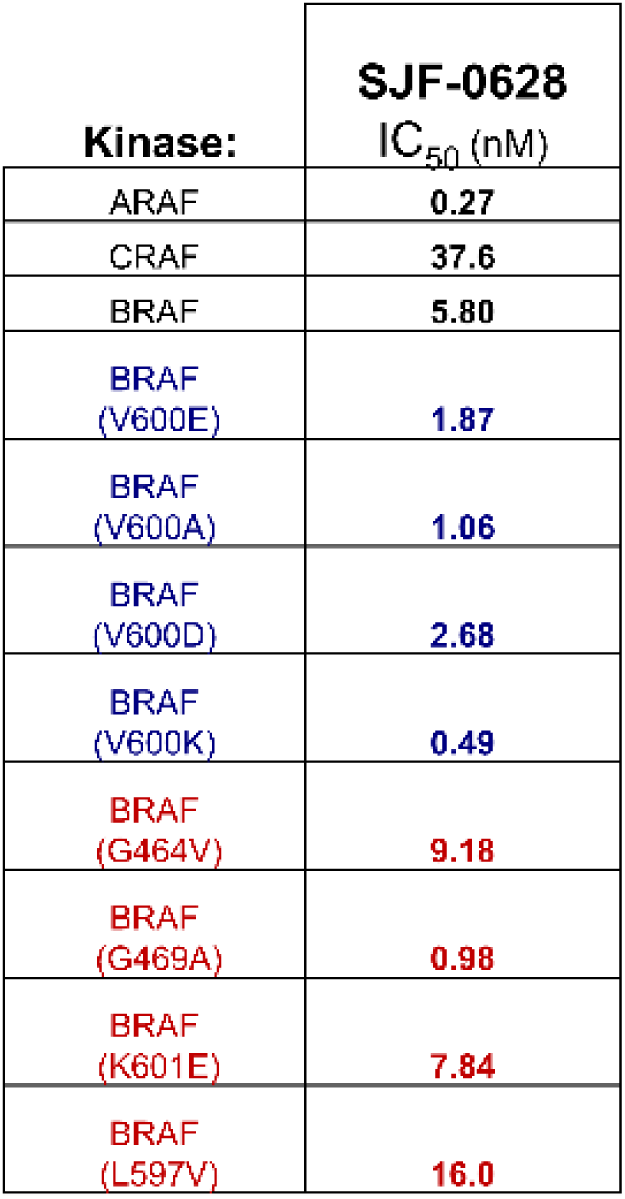
Table of IC_50_ values from radio labeled kinase assay (n=2)

To determine whether differences in ternary complex formation explain the differential degradation observed, we performed a pulldown experiment by immobilizing recombinant GST-tagged VHL/Elongin B/Elongin C (VBC) on glutathione-sepharose and incubating the beads with purified full-length BRAF and increasing amounts of PROTAC, with the goal of detecting a VHL: PROTAC: BRAF trimer. Indeed, there was a dose-dependent increase of BRAF^WT^ and BRAF^V600E^ complexed with the VBC at comparable levels (Fig. 2b), thus the innate capacity to form a trimer appears equivalent in both cases and therefore does not contribute to BRAF isoform selectivity of degradation. However, mutant and BRAF^WT^ adopt different conformations and form unique complexes within cells^4, 46^. Thus, we hypothesized that BRAF cellular conformations or associated proteins may affect trimer formation. To investigate this, we performed pull-down assays using NIH3T3 cell lysates expressing V5-BRAF^WT^ or V5-BRAF^V600E^. Interestingly, we found that while BRAF^WT^ forms a ternary complex with VHL in this lysate-based assay, this occurs to a lesser extent than it does for BRAF^V600E^ (> 3 fold greater than BRAF^WT^ at 5000 nM), as well as for BRAF^K601E^ and BRAF^G466E^ (Fig. 2c and Extended Data Fig. 6a). This result suggested that the *in vitro* trimer formation system using purified recombinant BRAF may not fully recapitulate PROTAC-induced complex formation in cells.

Therefore, we sought to examine ternary complex formation in intact cells. We treated NIH3T3 cells with SJF-0628 for 1 hour (to minimize degradation) and pulled down V5-BRAF and any associated VHL E3 ligase components. While SJF-0628 induced an interaction of all three mutant BRAF classes with Cullin 2, the E3 adaptor of VHL, it did not promote BRAF^WT^ trimer formation (Fig. 2d). At higher concentrations, we observe some ternary complex formation with BRAF^WT^, but 20-fold less than that seen with mutant BRAF (Extended Data Fig. 6b,c). Furthermore, while all three mutant classes are ubiquitinated in cells, BRAF^WT^ is not (Fig. 2e, Extended Data Fig. 6d,e). Overall, these studies show that BRAF^WT^ weakly associates with the E3 ligase complex in the cellular milieu, and this leads to minimal ubiquitination and degradation as compared to mutant BRAF; hence, the mutant selectivity of the PROTAC.

### Relief of negative feedback sensitizes BRAF^WT^ to SJF-0628 induced degradation

Our results suggest that BRAF^WT^ conformation and complex heavily influences its degradability. Therefore, we directly interrogated how these properties affect the ability to induce BRAF^WT^ degradation. Studies have shown that MEK inhibition potentiates the activated state of RAF by attenuating feedback inhibition from downstream effectors to stabilize/increase RAF dimerization and association with RAS^47, 48^. Therefore, we hypothesized that MEK inhibition would allow for enhanced BRAF^WT^ degradation.

To test this, we pre-treated NIH3T3 cells with the allosteric MEK inhibitor, trametinib, followed by increasing doses of SJF-0628. Interestingly, trametinib addition caused dose-dependent PROTAC-induced BRAF^WT^ degradation, as well as increased MEK phosphorylation (Fig. 3a, Extended Data Fig. 7a); trametinib effectiveness was confirmed by the minimal ERK phosphorylation observed. Trametinib pre-treatment did not affect BRAF^V600E^ degradation, nor did it promote epimer-induced degradation of BRAF^V600E^ or BRAF^WT^ in NIH3T3 cells (Extended Data Fig. 7b,c). SJF-0628 also induced BRAF^WT^ degradation in NIH3T3 cells that were pre-treated with cobimetinib - a second, structurally distinct allosteric MEK inhibitor (Fig. 3a). In OVCAR-8 cells, pre-treatment with cobimetinib or trametinib also enabled PROTAC-induced BRAF^WT^ degradation while increasing MEK and CRAF phosphorylation (Fig. 3b). Cobimetinib pre-treatment stimulated MEK phosphorylation within 30 minutes, and enabled PROTAC-induced degradation of BRAF^WT^ within 4 hours, with complete degradation observed after 12 hours (Extended Data Fig. 7d). No changes in BRAF^WT^ mRNA levels were observed (Extended Data Fig. 7e) confirming that BRAF^WT^ downregulation by SJF-0628 in the presence of cobimetinib is, indeed, post-translational. Moreover, we observed markedly increased trimer formation in cell lysates pre-treated with cobimetinib (Fig. 3c). In addition, MEK inhibitor-pretreated cells generated a 4-fold increase in PROTAC-induced Cullin-2 association with BRAF^WT^ (Fig. 3d) and increased SJF-0628-dependent BRAF^WT^ ubiquitination (Fig 3e) showing that the BRAF^WT^ degradation observed occurred via a PROTAC mechanism of action. These data, taken together, support our hypothesis that the activated conformation drives the ability of the PROTAC to degrade BRAF^WT^.

**Figure 3.**
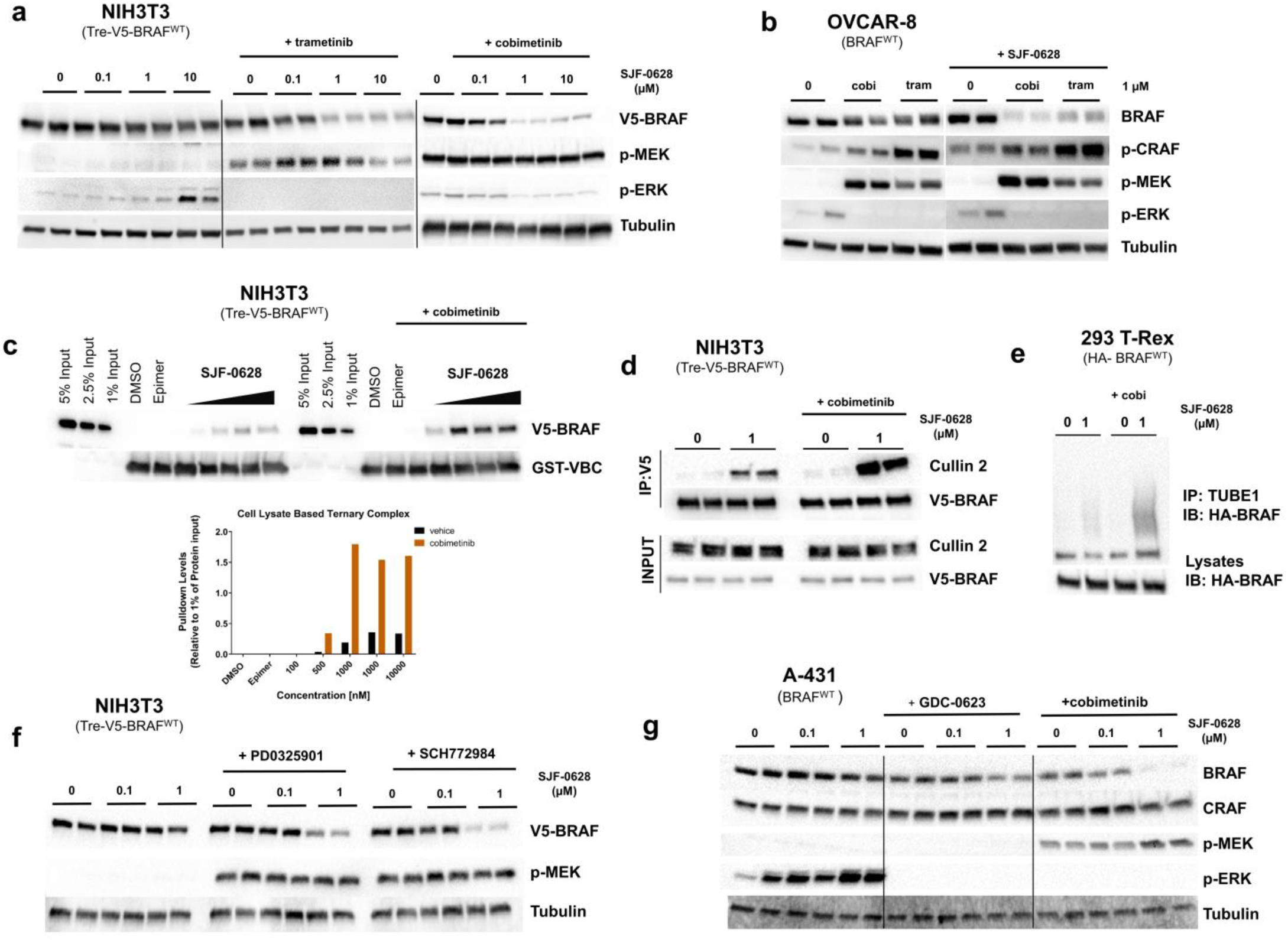
MEK inhibitors that activate BRAF also sensitize BRAF to PROTAC-induced ubiquitination and degradation. **a**, NIH3T3 cells with trametinib (1µM, 5 hours) or cobimetinib (500 nM, 3 hours) pre-treatment subsequently treated with increasing amounts of SJF-0628 (20 hours) promote degradation of BRAF^WT^ and show a marked increase in p-MEK. **b**, OVCAR8 cells pre-treated with cobimetinib and trametinib (1µM, 2 hours) promote MEK and CRAF phosphorylation as well as BRAF degradation in the presence of SJF-0628. **c**, Cell lysate-based ternary complex assay shown in 2**c** but using NIH3T3 lysates expressing BRAF^WT^ and pre-treated with DMSO or 1µM cobimetinib for 3 hours. Cobimetinib pre-treatment promotes ternary complex formation. **d**, V5-BRAF immunoprecipitation in NIH3T3 cells pre-treated with 1µM of cobimetinib (2 hours) followed by treatment of SJF-0628 for 2.5 hours. **e**, TUBE1 pulldown in 293 T-Rex cells stably expressing HA-BRAF^WT^ treated with cobimetinib (cobi) (2 hours, 1µM) and subsequently treated with SJF-0628 (2 hours). **f,** NIH3T3 cells pre-treated with 1µM PD0325901 (MEK inhibitor) or SCH772984 (ERK inhibitor) for 3 hours followed by treatment with indicated amount of SJF-0628 for 20 hours. **g**, A431 cells pre-treated with GDC-0623 and cobimetinib (500nM for 3 hours) then treated with SJF-0628 for 20 hours.

To rule out other aspects of MEK inhibition that may promote BRAF^WT^ degradation by SJF-0628, we undertook a series of pharmacological studies. In addition to inhibiting negative feedback and increasing BRAF activity, cobimetinib and trametinib have also been shown to decrease BRAF^WT^ association with MEK^49^. As such, MEK might hinder BRAF^WT^ ternary complex formation with SJF-0628 and VHL, preventing degradation of BRAF^WT^. To investigate this, we pre-treated cells with an early generation MEK inhibitor, PD0325901, known to stabilize the BRAF:MEK complex^50^ (Extended Data Fig. 8a). However, cells pretreated with PD0325901 also enabled PROTAC-mediated BRAF^WT^ degradation (∼80% degradation at 1 μM) (Fig. 3f). We hypothesized that if relief of MAPK negative feedback promotes BRAF^WT^ degradation, ERK inhibition would do the same. As predicted, pretreatment with the selective ERK inhibitor SCH772984^51^ at 1 µM enabled ∼90% degradation of BRAF^WT^ by the PROTAC (Fig. 3f). SCH772984 also does not disrupt BRAF-MEK association, further demonstrating that the presence of MEK does not affect BRAF^WT^ degradation (Extended Data Fig. 8a).

GDC-0623 is an allosteric MEK inhibitor which binds MEK in a manner that sequesters BRAF and hinders dimerization with itself or CRAF (Extended Data Fig. 8a) and membrane localization^49, 52^. Therefore, treatment with GDC-0623 dampens relief of feedback induced signaling on RAF kinase activity. We hypothesized that if the BRAF^WT^ conformation induced by the previously-tested MEK/ERK inhibitors is primarily responsible for enabling the kinase’s degradation by the PROTAC, then GDC-0623 would enable significantly less degradation. As expected, GDC-0623 pre-treatment permitted only minimal PROTAC-dependent degradation of BRAF^WT^ – far less than was seen in parallel treatment with cobimetinib (Fig. 3g, Extended Data Fig. 8b,c). These results further show that SJF-0628 selectively induces degradation of BRAF^WT^ in its active conformation. Indeed, studies such as Rock *et al.* show that all three mutant BRAF classes (including kinase dead mutations) favor an open, active conformation^53^. Thus, by stimulating BRAF^WT^ activity (e.g., RTK upregulation, RAS mutations, relief of negative feedback), we promote an open conformation which is susceptible to increased ternary complex formation and therefore degradation.

### SJF-0628 successfully inhibits cell growth in mutant-BRAF driven cancer cells

Next, we compared the effects on cell growth of inhibiting both the enzymatic and scaffolding roles of BRAF using SJF-0628, with those of an ATP-competitive inhibitor that targets only its catalytic function (vemurafenib and degradation-incompetent epimer, SJF-0661). In SK-MEL-28 cells (Class 1), vemurafenib and SJF-0661 inhibited cell growth with an EC_50_ of 215 ± 1.09 nM and 243 ±1.09 nM respectively, while SJF-0628 showed an EC_50_ of 37 ± 1.2 nM (Fig. 4a). The 6-fold increase in potency of SJF-0628 occurred despite the compounds having similar *in vitro* binding (vemurafenib = 27 nM, SJF-0628 = 39 nM, SJF-0661= 64 nM) (Extended Data Fig. 9a). In SK-MEL-239 C4 cells (BRAF^WT^/ BRAF-p61^V600E^), while vemurafenib and SJF-0661 had a minimal effect, SJF-0628 induced ∼80% decrease in cell growth with an EC_50_ of 218 nM ± 1.06 (Fig. 4b This result shows that targeted degradation can be used to overcome acquired resistance to BRAF inhibitor-based therapies. A 5-day treatment in SK-MEL-246 cells (G469A) SJF-0628 efficaciously inhibited cell growth with an EC_50_ of 45 ± 1.11nM and the epimer showed an EC_50_ 278± 1.07nM. However, vemurafenib caused some inhibition of SK-MEL-246 cellular growth at concentrations above 1μM, which was not sustained at 10 μM (Fig. 4c). In H1666 cells (G466V Class 3), SJF-0628 was able to induce 65% cell growth inhibition while vemurafenib showed less than 50% (Fig. 4d). Despite causing >70% inhibition of p-ERK (Extended Data Fig. 3d), SJF-0628 showed minimal inhibition of cell growth in CAL-12-T cells (G466V, Class 3) (Fig. 4e).

**Figure 4.**
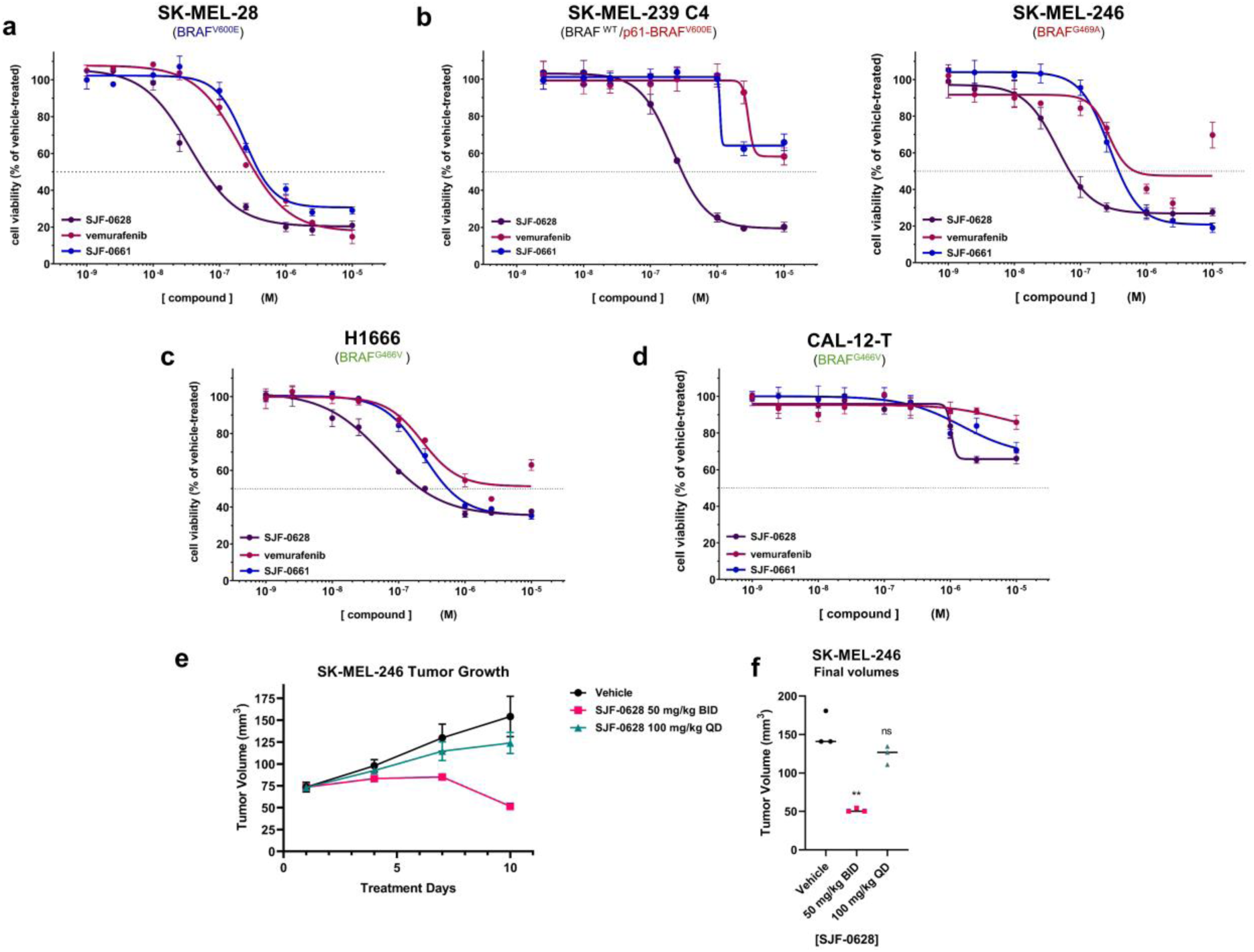
SJF-0628 outperforms vemurafenib in inhibiting growth of cell lines expressing mutant BRAF. **a,** Cell proliferation assay in SK-MEL-28 cells treated with increasing amounts of vemurafenib, SJF-0628, or SJF-0661 for 3 days (mean ± s.d., n=3). EC_50_ = 215 ± 1.09 nM, 37 ± 1.2 nM, and 243 ± 1.09 nM respectively. **b,** Cell proliferation assay in vemurafenib resistant SK-MEL-239-C4 cells treated with increasing amounts vemurafenib, SJF-0628, or SJF-0661 for 5 days (mean ± s.d., n=3). **c,** Cell proliferation assay in SK-MEL-246 (Class 2) cells treated with increasing amounts vemurafenib, SJF-0628, or SJF-0661 for 5 days (mean ± s.d., n=3) **d,** SJF-0628 EC50=218 nM±1.06 c,H1666 cells treated with SJF-0628, vemurafenib, or SJF-0661 for 5 days (mean ± s.d., n=3). **d,** Treatment of CAL-12-T cells with vemurafenib, SJF-0628, or SJF-0661 for 5 days shows minimal effect on cell viability (mean ± s.d., n=3). **e,** Results of an efficacy study in SK-MEL-246 tumor xenografts implanted in female athymic mice showing tumor regression with 50 mg/kg IP twice daily. **f,** Scatter plot result of final volumes (** *P* value < 0.01). *P* value calculated by unpaired t-test.

SJF-0628 causes potent degradation of all BRAF mutant classes and has minor effects on WT RAF. This result suggests that the PROTAC will be effective in treating tumors driven by these mutations with minimal on target toxicity, thus we tested the effects of SJF-0628 in an A375 (BRAF^V600E^) murine xenograft model. Mice treated with SJF-0628 (three days; 50 mg/kg or 150 mg/kg) showed marked degradation of BRAF in the xenograft at both concentrations (D_MAX_ > 90 %) (Extended Data Fig. 9b). Because SJF-0628 successfully induced degradation *in vivo*, we tested its effect on tumor growth in the Class 2 SK-MEL-246 melanoma xenograft model (BRAF^G469A^, Class 2). Strikingly, while once daily 100 mg/kg treatment showed a minor response, twice daily treatment of 50 mg/kg induced tumor shrinkage beyond the initial tumor size within 10 days (Fig. 4e,f). We did not observe a significant body weight loss with either dose (Fig. S9C). Thus, through targeted degradation, SJF-0628 is successfully able to exhibit a significant antitumor effect.

## Discussion

Each BRAF mutation alters how the protein signals in distinct ways, therefore, careful consideration must be taken to select the appropriate inhibitor. Despite tremendous effort to create therapies that target diverse BRAF mutations, all three currently FDA-approved drugs target Class 1 BRAF mutants alone. Furthermore, resistance to current drugs inevitably occurs. Thus, we have developed a vemurafenib-based PROTAC, SJF-0628, that outperforms vemurafenib in inhibiting MAPK signaling and growth of BRAF^V600E-^driven cancer cells. Importantly, the mutant BRAF-targeting PROTACs described here spare WT RAF, thus widening the potential therapeutic window of this new class of anti-tumor drugs.

We find that measuring ternary complexes (BRAF:PROTAC:VHL) in cells or cell lysates to be more predictive of degradation than *in vitro* studies with purified proteins. Our *in vitro* pull-down assays likely contained highly dimerized BRAF in the active conformation, resulting in artificially high levels of ternary complex formation. However, in cells BRAF exists in a closed inactive conformation, which is less conducive to ternary complex formation allowing it to escape degradation. So by promoting its activation, BRAF^WT^ adopts an open conformation, similar to mutant BRAF, and is thus susceptible to SJF-0628-induced degradation. This finding suggests that intracellular protein conformation can affect PROTAC-induced degradation. As PROTACs require induced protein-protein interactions to function, it is important to consider protein interactions within the cellular context that might encourage or discourage degradation. Importantly for the development of new PROTACs, we show that we can control induced protein degradation (i.e., BRAF^WT^) without modifying the PROTAC itself but by manipulating its signaling pathway.

While preparing this manuscript, Han *et al.* described cereblon-based PROTACs that degrade BRAF^V600E^ but spares BRAF^WT^. However, their PROTACs do not allow for additional MAPK inhibition and cause less cell proliferation inhibition than the parent inhibitor. Beyond this initial observation, we also successfully target vemurafenib-resistant BRAF mutations. This includes mutants that have both acquired (p61 V600E) and intrinsic (Class 2) resistance to vemurafenib. Furthermore, we show that SJF-0628 can be used to successfully target Class 2 mutants *in vivo*. In addition, we make Class 3 BRAF mutants, which cannot be targeted with traditional small molecule inhibitors, therapeutically accessible through targeted degradation. Indeed, this is the first demonstration of PROTAC induced degradation of a pseudokinase. Thus, using PROTACs, we are able to expand the druggable space to a class of proteins with immense cancer relevance (HER3, ROR2, etc.). In summary, this study demonstrates that the PROTAC technology is an attractive strategy for targeting difficult oncoproteins such as mutant BRAF.

## Acknowledgments

We thank all members of the Crews and Rosen labs for helpful discussions. We are thankful for Jemilat Salami-Oyenuga, Ashton Lai, George Burslem, Anna Wadhwa, Hanqing Dong, Kanak Raina and Alicia Morgan for early screening of BRAF-targeting PROTACs, technical expertise, and meaningful discussion. C.M.C. gratefully acknowledges support from the NIH (R35CA197589) and the American Cancer Society. S.A. is supported by the HHMI Gilliam Fellowship. J.H. is supported by an R50 Research Specialists award from the NCI (R50 CA211252-02).

## Contributions

C.M.C. and N.R. conceived the project and supervised experiments. SJF designed and synthesized the PROTACS. S.A., Y.G., Z.Y., J.H., K.T.G.S, and L.V planned and carried out the biological experiments. SA wrote the manuscript with assistance from all authors.

## Materials and Methods

### Cell Culture and Reagents

SK-MEL-28 (E-MEM), A-431 (D-MEM), and SKBR3(RPMI) cells was obtained from ATCC. Inducible 293-Trex (DMEM), CAL-12T (DMEM), H1666 (RPMI), and SK-MEL-30 cells (RPMI) were obtained from Arvinas. We thank the Kupfer for HCT-116 cells (DMEM), Frank Slack for H23 cells, A. Houghton and P. Chapman for SKMEL 246 cells (DMEM) and SK-MEL-239 C4 cells (DMEM;1 µM vemurafenib), Joyce Lui for OVCAR-8 cells, and the Trevor Bivona for HCC364 cells (10 µM vemurafenib). Inducible expression NIH3T3 cells were maintained in DMEM;50 μg ml^−1^ hygromycin and 0.2 μg ml^−1^ puromycin). All cell lines tested negative for mycoplasma. All media was supplemented with 10% fetal bovine serum and 1% penicillin– streptomycin and grown in a humidified incubator at 37 °C and 5% CO_2_.

### Compounds, Agarose and Recombinant Protein

From Selleckchem: PLX-4032 (S1267), trametinib (S2673), cobimetinib (S8041), GDC-0623 (S7553), MLN-4924 (S7109), SCH772984 (S7101) and PD0325901 (S1036). Epoxomicin was synthesized in the Crews lab. Full length BRAF was purchased from Proteros.

## Experimental

### PROTAC treatment and Immunoblotting

Cells were plated in 6 well dishes (5×10^5^-8×10^5^ cells) and allowed to attach overnight. Cells were treated with SJF-0628 or SJF-0661 for 20-24 hours (unless otherwise stated). The plates were then placed on ice and washed 1x with chilled PBS and lysed in buffer containing 25mM Tris-HCl [pH 7.4], 0.25% sodium deoxycholate, 150 mM NaCl, 1 % Triton X-100, supplemented with protease inhibitors (1x Roche protease inhibitor cocktail) and phosphatase inhibitors (10 mM NaF, 1 mM Na_3_OV_4_, and 20 mM β-glycerophosphate). Lysates were then cleared at 15,000 RPM for 10 minutes at 4 °C. Protein concentrations of the supernatants were then quantified using a Pierce BCA Protein Assay. 12-40 µgs of protein were separated using a gradient (4-20%) Criterion TGX precast gel and transferred unto a nitrocellulose membrane. The membranes were then blocked in 5% non-fat milk in TBST (Tris-buffered Saline with Tween 20) for 1 hour before probing with the indicated primary antibody overnight. Membranes were imaged using Bio-Rad Image Lab software using ECL prime detection reagent (GE Healthcare, RPN2232 or ThermoScientific, 34095)

### Antibodies

Primary antibodies from Cell-Signaling: V5 (no. 13202), BRAF (no. 14814s), CRAF (no.53745), anti-p202/p204-ERK1/2 (p-ERK1/2) (no. 4370), anti-ERK1/2 (no.4696s) anti-MEK1/2 (no. 8727), anti-p217/p221-MEK1/2 (p-MEK1/2) (no. 9154), GAPDH (no.2118S), VHL (no. 68547), anti-phospho-c-Raf (Ser338) (no. 9427), anti-HA (no. 2367), anti-ARAF(no. 75804s), anti-ubiquitin (no. 43124), RBX-1 (no. 11922S) anti-p-HER2/ErbB2 (Y1221/1222) (6B12) (no. 2243S); primary antibody from Millipore: anti-tubulin (16-232); primary antibody from Invitrogen: anti-Cullin 2(700179); primary antibody from Lifespan Biosciences: anti-KRAS (LS-C175665). Secondary antibodies were from ThermoFisher: anti-rabbit HRP (31460) and anti-mouse HRP (31444).

### Cell proliferation

Cells (2,5000 to 5,000) were seeded in 96 well plates and treated with compound for the indicated lengths of time (between 72 – 96 hours). 2 mg/ml MTS (Promega Corp., Madison, WI : G5421) and 25 μM phenazine methosulfate (Sigma, St. Louis, MO) were combined 19:1 and then added to cells (1 volume combined reagent: 5 volumes medium) and incubated for 1-3 hours. Mitochondrial reduction of MTS to the formazan derivative was monitored by measuring the medium’s absorbance at 480 nm using a Perkin Elmer Envision Plate reader.

### Protein Purification

For the expression of GST-tagged VHL:Elongin B:Elongin C (herein referred to as GST-VBC), wild-type human VHL, Elongin B, and Elongin C were co-expressed in *E. coli*. BL21(DE3) cells were co-transformed with pBB75-Elongin C and pGEX4T-2-VHL-rbs-Elongin B and selected in LB medium containing carbenicillin (100 µg mL−1) and kanamycin (25 µg mL−1) at 37 °C until OD 600 = 0.8, at which point the culture was chilled to 16 °C and induced with 0.4 mM IPTG for 16 h. Cells were homogenized and lysed using a Branson digital sonifier with lysis buffer composed of 50 mM Tris [pH 8.0], 200 mM NaCl, 5% glycerol, 5 mM DTT containing a 1 X protease inhibitor cocktail tablet (Roche). Clarified cell lysate was applied to glutathione sepharose 4B beads (GE Life Science) and gently rotated for 2 h at 4 °C. Beads were washed with four column volumes of lysis buffer, followed by four column volumes of elution buffer (50 mM Tris pH 8.0, 200 mM NaCl, 10 mM glutathione). Eluted protein was assessed for identity and purity via Coomassie staining of sample run on an SDS-PAGE gel and pure elutions were pooled, concentrated, and diluted in ion-exchange buffer A (30 mM Tris pH 8.0, 5% glycerol, 1 mM DTT) until the salt concentration was 50 mM, before loading onto a Mono Q 5/50 GL column (GE Life Sciences). The protein was subjected to a linear gradient of NaCl (0–500 mM NaCl) using ion-exchange buffer B (30 mM Tris 8.0, 1 M NaCl, 5% glycerol, 1 mM DTT). Fractions were then assessed for purity via Coomassie, pooled, concentrated, and run on a Superdex-200 column (GE Life Sciences) using size-exclusion buffer (30 mM Tris pH 8.0, 100 mM NaCl, 10% glycerol, 1 mM DTT). Pure fractions of GST-VHL were pooled, concentrated to ∼5 mg mL−1, aliquoted, and flash-frozen before storing at −80 °C.

### Ternary Complex Assays

Glutathione sepharose 4B was washed twice with water and then blocked for one hour at room temperature with 10% BSA in TBST. The beads were then washed again twice with TBST and once with wash buffer (50mM HEPES pH 7.5, 150mM NaCl, 1mM DTT, 0.01% NP40, 5 mM MgCl_2_, 10% glycerol) and then purified GST-VBC was immobilized for two hours at 4°C at 2.5 pmole per μL of beads. The beads were then washed thrice with wash buffer, resuspended and BRAF WT or V600E protein was added at 500 nM per 50 μL reaction with 5 μL of beads. The bead:BRAF mixture was then aliquoted to separate tubes and PROTAC was added at the indicated concentration (PROTACs were intermediately diluted in 50% DMSO) and this was incubated at 4°C for two hours. The beads were washed 4 times with 1 mL column of TBST and then eluted with SDS loading buffer.

For experiments in which the input substrate is a whole cell lysate, the sample was prepared as follows: 15 mm dishes of confluent NIH3T3 cells were doxycycline induced overnight, after which the cells were washed with DPBS and lysed using lysis buffer (50mM Tris-HCl, pH 7.4, 015M NaCl, 1mM EDTA,1%NP40, 10%glycerol). The lysate was cleared by centrifugation and then added to the beads as an input substrate, as above. For MEK inhibitor comparison, NIHT3 cells were pre-treated with 1µM of cobimetinib for 3 hours.

### Cellular Immunoprecipitation and Ubiquitination Assay

Doxycycline-induced NIH3T3 cells or 293 T-Rex cells that express indicated BRAF isoform were seeded in 10 cm dishes overnight. Cells were then treated for 1 hour with PROTAC or DMSO. Cells were then placed on ice, washed with ice-cold 1X PBS and lysed in 500 µL modified 1X lysis buffer (50 mM Tris-HCl, pH 7.4, 0.15M NaCl, 1mM EDTA, 1%NP40, 10% glycerol) containing 5 mM 1,10-phenanthroline monohydrate, 10 mM N-ethylmaleimide, 20 µM PR-619, and 1X protease inhibitor cocktail (Roche). Lysates were spun down at 14,000 × g at 4 °C for 10 min. Equal amounts of lysate was aliquoted onto 20 µL (bed volume) of anti-V5-beads (Sigma, A7345). V5-containing proteins were immunoprecipitated from lysates for 2 hours at 4 °C with gentle rotation, after which samples were spun down at 6000 × g at 4 °C for 2 min and the beads were washed 4 times with DPBS. Beads were resuspended in 1X lithium dodecyl sulfate (LDS) sample buffer containing 5% 2-mercaptoethanol (ß-ME). Immunoprecipitated protein was eluted off the beads by heating at 95 °C for 5 min and the supernatant was run on an SDS-PAGE gel and evaluated for the presence of immunoprecipitated V5-tagged proteins, as well as Cullin 2. Input refers to the normalized input lysate loaded onto V5-sepharose beads.

TUBE1 immunoprecipitation experiments were carried out exactly as described above, except that 1 equal lysate was loaded onto 20 µL TUBE1 agarose (LifeSensors) resin per sample and washed with TBST.

### Radiolabeled Kinase Assays

Kinase assays were performed by Reaction Biology Corps by their protocol in duplicate using Km amounts of ATP calculated for each kinase.

### Elisa Kinase inhibition Assay

Kinase assays were performed by Carna Biosciences by their protocol in duplicate using K_m_ amounts of ATP calculated for each kinase.

### RT-PCR

Cells were seeded in 12 well plates and treated as described. RNA was isolated with the RNeasy Mini Kit (QIAGEN) and 1 μg of total RNA was reverse transcribed using the High Capacity cDNA Reverse Transcription Kit (Applied Biosystems). SYBR Green PCR master mix (Kapa Biosystems) was used for qRT-PCR samples were performed and analyzed in triplicate. Relative RNA expression levels were calculated using the ddCt method and normalized to control samples and beta-tubulin was used for normalization. Primers used in this study are as follows:

Beta_Tub_F: 5ʹ-TGGACTCTGTTCGCTCAGGT-3ʹ

Beta_Tub_R: 5ʹ-TGCCTCCTTCCGTACCACAT-3ʹ

BRAF_F: 5ʹ-GAGGCGTCCTTAGCAGAGAC-3ʹ

BRAF_R: 5ʹ-AAGGAGACGGACTGGTGAGAAF-3ʹ

### Quantitation and Statistical Analysis of Western Blots

Western blot data was quantified by using the band feature in Image Lab, and values were averaged and analyzed in GraphPad Prism. DC_50_ and D_MAX_ values were fitted using a three parameter [inhibitor] versus response and reported directly from the Prism output. Mean± s.t.d reported and unpaired t-tests were performed in GraphPad Prism.

### Animal Studies

#### A375 xenograft study

5 million A375 cells were subcutaneously implanted in female nu/nu mice. Tumors were randomized after a period of 10 days into groups with an average tumor size of 350 mm^3^, and treated with vehicle (5% DMSO, 5% EtOH and 20% Solutol HS15 in D5W), 50 mg/kg SJF-0628, or 150 mg/kg SJF-0628 (4 mice per arm) intraperitoneally once a day for 3 days. Mice were sacrificed eight hours after the final dose. Tissues were harvested, flash frozen and lysed in 1X cell lysis buffer (Cell Signaling #9803) supplemented with protease and phosphatase inhibitors. Harvested tumors were disrupted using metal beads in a Tissuelyser. Homogenates were normalized for protein content and analyzed using SDS-PAGE and western blotting.

#### SK-MEL-246 xenograft study

10 million SK-MEL-246 cells were subcutaneously implanted in female nu/nu mice. The tumor volumes and mice weights were measured twice a week after the implantation. The i.p. treatments with vehicle (5% DMSO, 5% EtOH and 20% Solutol HS15 in D5W), 50 mg/kg (BID) or 100mg/kg SJF-0628 (QD), (3 mice/group) were started when the tumor volumes reached an average of 100 mm^3^. All studies were performed in compliance with institutional guidelines under an IACUC approved protocol. Investigators were not blinded when assessing the outcome of the *in vivo* experiments.

## Experimental-Chemistry

### 1. Chemical synthesis

#### General comments

Unless otherwise indicated, common reagents or materials were obtained from commercial source and used without further purification. Tetrahydrofuran (THF), dimethylformamide (DMF), and Dichloromethane (DCM) were dried by a PureSolv^TM^ solvent drying system. Flash column chromatography was performed using silica gel 60 (230-400 mesh). Analytical (TLC) and preparative (PTLC) thin layer chromatography was carried out on Merck silica gel plates with QF-254 indicator and visualized by UV or iodine. ^1^H and ^13^C NMR spectra were recorded on an Agilent DD_2_ 500 (500 MHz ^1^H; 125 MHz ^13^C) or Agilent DD_2_ 600 (600 MHz ^1^H; 150 MHz ^13^C) or Agilent DD_2_ 400 (400 MHz ^1^H; 100 MHz ^13^C) spectrometer at room temperature. Chemical shifts were reported in ppm relative to the residual CDCl_3_ (δ 7.26 ppm ^1^H; δ 77.00 ppm ^13^C), CD_3_OD (δ 3.31 ppm ^1^H; δ 49.00 ppm ^13^C), or *d^6^*-DMSO (δ 2.50 ppm ^1^H; δ 39.52 ppm ^13^C). NMR chemical shifts were expressed in ppm relative to internal solvent peaks, and coupling constants were measured in Hz. (bs = broad signal). Mass spectra were obtained using electrospray ionization (ESI) on a time of flight (TOF) mass spectrometer. Compounds **1**^55^, **VHL** ligands^56^ 5 and 6 were prepared according with the literature or acquired commercially.

**Scheme 1.**
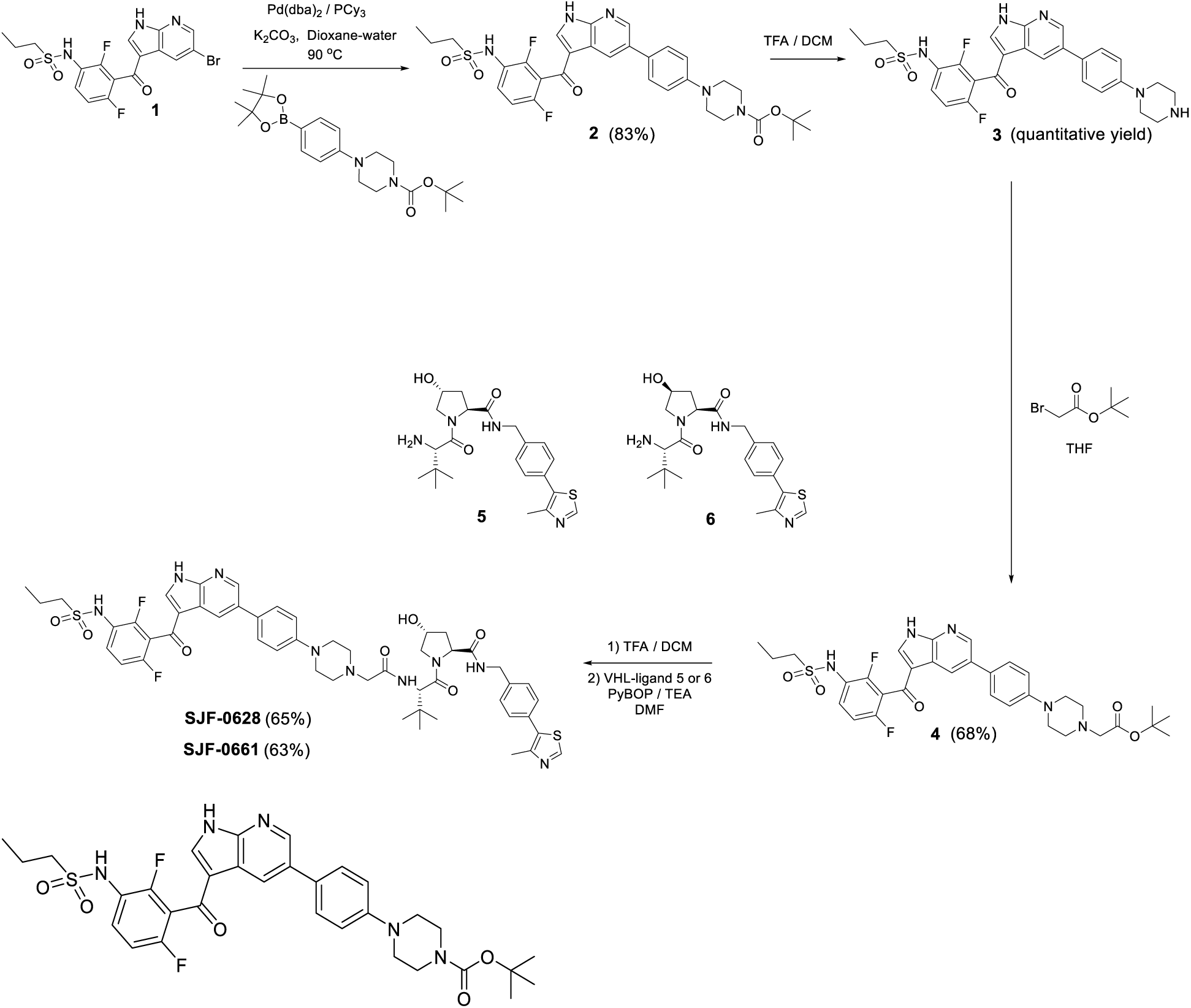
Synthetic Approach for SJF-0628 and SJF-0661

#### *tert*-Butyl 4-(4-(3-(2,6-difluoro-3-(propylsulfonamido)benzoyl)-1H-pyrrolo[2,3-b]pyridin-5-yl)phenyl) -piperazine-1-carboxylate (2)

To a solution of N-[3-(5-bromo-1H-pyrrolo[2,3-b]pyridine-3-carbonyl)-2,4-difluoro-phenyl]propane-1-sulfonamide (1) (70.8 mg, 0.155 mmol) in Dioxane (6 ml) was added tert-butyl 4-(4-(4,4,5,5-tetramethyl-1,3,2-dioxaborolan-2-yl)phenyl)piperazine-1-carboxylate (60.0 mg, 0.155 mmol), K_2_CO_3_ (64.2 mg, 0.465 mmol), Tricyclohexyl phosphine (4.33 mg, 0.0155 mmol)and water (2 mL). Then the reaction mixture was de-gassed under vacuum and purged with argon (5×), Pd(dba)_2_ (4.44 mg, 0.00773 mmol) was added into and the reaction mixture was heated at 90 °C for 3 h. By TLC small amounts of SM (Hex:EtOAc, 3:7), the reaction mixture was filtered in vacuum over a celite pad, filtrate was poured into an aqueous saturated solution of NaCl (20 mL) and the product was extracted with EtOAc (2×20 mL). The EtOAc layers were combined, dried (Na_2_SO_4_) and concentrated in vacuum. The crude material was diluted in DCM and purified by flash chromatography (SiO_2_-12g, Hexane:EtOAc, 9:1 to 100% EtOAc in 15 min) to give 82 mg (83%) of product. ^1^H NMR (500 MHz, DMSO-d6) δ 12.92 (bs, 1H), 9.76 (bs, 1H), 8.66 (d, J = 2.2 Hz, 1H), 8.55 (s, 1H), 8.19 (s, 1H), 7.75 – 7.46 (m, 3H), 7.28 (t, J = 8.7 Hz, 1H), 7.10 (d, J = 8.7 Hz, 2H), 3.49 (t, 4H), 3.19 (t, J = 5.2 Hz, 4H), 3.16 – 3.05 (m, 2H), 1.74 (h, J = 7.5 Hz, 2H), 1.43 (s, 9H), 0.96 (t, J = 7.4 Hz, 3H). ^13^C NMR (151 MHz, DMSO-d6) δ 180.61, 156.04 (dd, J = 246.5, 6.9 Hz), 153.88, 152.34 (dd, J = 249.4, 8.6 Hz), 150.35, 148.44, 143.61, 138.60, 131.47, 128.92 – 128.75 (m), 128.19 (d, J = 161.3 Hz), 126.05, 121.94 (dd, J = 13.6, 3.6 Hz), 118.96 – 117.87 (m), 117.60, 116.33, 115.61, 112.36 (dd, J = 22.6, 3.2 Hz), 79.03, 53.42, 48.08, 43.72, 42.58, 28.09, 16.87, 12.64 . LC-MS (ESI); m/z [M+H]^+^: Calcd. for C_32_H_36_F_2_N_5_O_5_S, 640.2405. Found 640.2541.

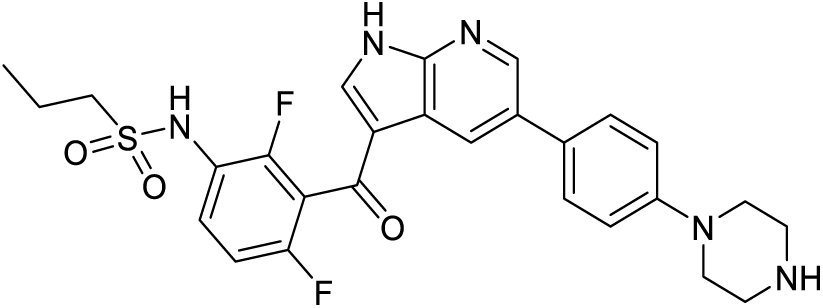

#### N-(2,4-difluoro-3-(5-(4-(piperazin-1-yl)phenyl)-1H-pyrrolo[2,3-b]pyridine-3-carbonyl)phenyl)propane-1-sulfonamide (3)

A solution of tert-butyl 4-[4-[3-[2,6-difluoro-3-(propylsulfonylamino)benzoyl]-1H-pyrrolo[2,3-b]pyridin-5-yl]phenyl]piperazine-1-carboxylate (**2**) (28.0 mg, 0.0438 mmol) in a mixture of DCM/TFA (3:1, 4 mL) was stirred for 1h at room temperature (by TLC no SM). The solvent was removed under vacuum and the residue was dried under high vacuum for 2h (23 mg of product, quantitative yield). Crude product was used in the next step without any further purification. LC-MS (ESI); m/z [M+H]^+^: Calcd. for C_27_H_28_F_2_N_5_O_3_S, 540.1880. Found 540.1949.

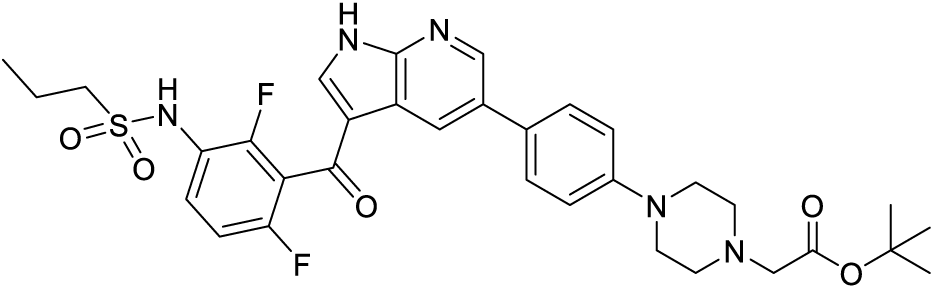

#### *tert*-Butyl 2-(4-(4-(3-(2,6-difluoro-3-(propylsulfonamido)benzoyl)-1H-pyrrolo[2,3-b]pyridin-5-yl)-phenyl) piperazin-1-yl)acetate (4)

To a solution of N-[2,4-difluoro-3-[5-(4-piperazin-1-ylphenyl)-1H-pyrrolo[2,3-b]pyridine-3-carbonyl]-phenyl]propane-1-sulfonamide (**3**) (23.0 mg, 0.0426 mmol) and TEA (0.0594 mL, 0.426 mmol) in DMF (1 ml) was added tert-butyl 2-bromoacetate (9.15 mg, 0.0469 mmol) and the resulting solution stirred for 3 h at rt. The reaction mixture was evaporated under vacuum. Crude product was purified by PTLC (DCM:MeOH:NH_4_OH, 90:9:1, 2×) to give 19 mg of pure product (69% yield). ^1^H NMR (500 MHz, DMSO-d6) δ 12.92 (bs, 1H), 9.76 (bs, 1H), 8.65 (t, J = 2.6 Hz, 1H), 8.55 (s, 1H), 8.18 (s, 1H), 7.76 – 7.42 (m, 3H), 7.28 (t, J = 8.3 Hz, 1H), 7.07 (d, J = 6.5 Hz, 2H), 3.33 (s, 2H), 3.30 – 3.16 (m, 4H), 3.16 – 3.04 (m, 2H), 2.81 – 2.55 (m, 4H), 1.84 – 1.67 (m, 2H), 1.43 (s, 9H), 0.96 (t, J = 7.2 Hz, 3H). ^13^C NMR (151 MHz, DMSO-d6) δ 180.57, 169.22, 156.02 (dd, J = 246.3, 7.0 Hz), 152.33 (dd, J = 249.4, 8.6 Hz), 150.42, 148.37, 143.55, 138.50, 131.53, 128.75 (d, J = 9.6 Hz), 128.17, 127.58, 125.93, 121.92 (dd, J = 13.6, 3.6 Hz), 118.25 (t, J = 23.6 Hz), 117.58, 115.80, 115.58, 112.33 (dd, J = 21.9, 3.0 Hz), 80.23, 59.21, 53.44, 51.81, 47.93, 27.82, 16.8, 12.62 . LC-MS (ESI); m/z [M+H]^+^: Calcd. for C_33_H_38_F_2_N_5_O_5_S, 654.2561. Found 654.2675.

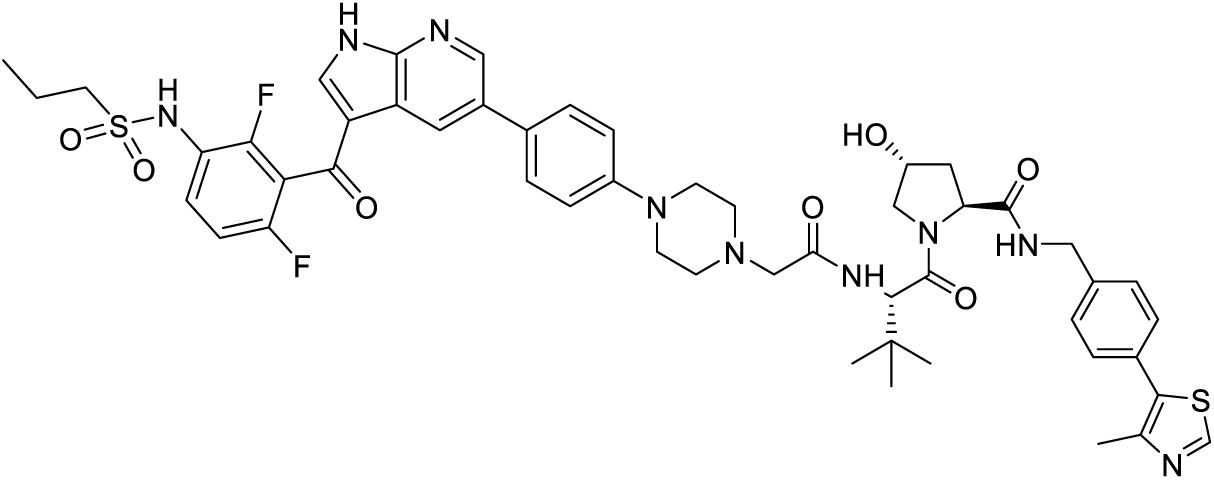

#### (2S,4R)-1-((S)-2-(2-(4-(4-(3-(2,6-difluoro-3-(propylsulfonamido)benzoyl)-1H-pyrrolo[2,3-b]pyridin-5-yl)phenyl)piperazin-1-yl)acetamido)-3,3-dimethylbutanoyl)-4-hydroxy-N-(4-(4-methylthiazol-5-yl)benzyl)pyrrolidine-2-carboxamide (SJF-0628)

A solution oftert-butyl 2-(4-(4-(3-(2,6-difluoro-3-(propylsulfonamido)benzoyl)-1H-pyrrolo[2,3-b]pyridin-5-yl)phenyl)piperazin-1-yl)acetate (**4**) (20.0 mg, 0.0306 mmol) in a mixture of TFA (2 ml, 13.46 mmol) and DCM (2 ml) was stirred for 5 h. Then the solvent was removed under vacuum and crude product was dried under high vacuum for 1 h. Crude product was used in the next step without any further purification (18.3 mg, quantitative yield). LC-MS (ESI); m/z: [M+H]^+^ Calcd. for C_29_H_30_F_2_N_5_O_5_S, 598.1935. Found 598.1953. To a solution of crude product from above; 2-(4-(4-(3-(2,6-difluoro-3-(propylsulfonamido)benzoyl)-1H-pyrrolo[2,3-b]pyridin-5-yl)-phenyl) - piperazin-1-yl)acetic acid (18.3 mg, 0.0306 mmol) and (2S,4R)-1-[(2S)-2-amino-3,3-dimethyl-butanoyl]-4-hydroxy-N-[[4-(4-methylthiazol-5-yl)phenyl]methyl]pyrrolidine-2-carboxamide; hydro-chloride (**5**) (17.2 mg, 0.0367 mmol) in DMF (1 ml) was added TEA (0.106 mL, 0.762 mmol) and PyBOP (19.1 mg, 0.0367 mmol) at room temperature. The reaction mixture was stirred for 4 h at the same temperature. TLC (DCM:MeOH:NH_4_OH, 90:9:1) shows no starting material (acid). The reaction mixture was evaporated to dryness under high vacuum. Crude product was diluted with EtOAc (10 mL) and washed with a saturated-aqueous solution of NaHCO_3_ (2×5 mL), organic extract was dried (Na_2_SO_4_), and evaporated under vacuum. Crude product was purified by PTLC (DCM:MeOH:NH_4_OH, 90:9:1, 2×) to give 20 mg of product (65% yield). ^1^H NMR (500 MHz, DMSO-d6) δ 12.92 (bs, 1H), 9.76 (bs, 1H), 8.91 (s, 1H), 8.66 (s, 1H), 8.65 – 8.45 (m, 2H), 7.85 (d, J = 9.1 Hz, 1H), 7.74 – 7.52 (m, 3H), 7.49 – 7.32 (m, 4H), 7.28 (t, J = 8.6 Hz, 1H), 7.09 (d, J = 8.5 Hz, 2H), 5.16 (d, J = 3.0 Hz, 1H), 4.54 (d, J = 9.6 Hz, 1H), 4.52 – 4.32 (m, 3H), 4.26 (dd, J = 15.7, 5.4 Hz, 1H), 3.76 – 3.58 (m, 2H), 3.27 (s, 4H), 3.21 – 2.95 (m, 4H), 2.68 (s, 4H), 2.40 (s, 3H), 2.07 (dd, J = 12.9, 7.7 Hz, 1H), 1.92 (ddd, J = 13.1, 9.0, 4.6 Hz, 1H), 1.75 (h, J = 7.5 Hz, 2H), 0.97 (s, 9H), 0.96 (t, J = 7.4 Hz, 3H). ^13^C NMR (126 MHz, DMSO-d6) δ 180.56, 171.77, 169.28, 168.48, 156.02 (dd, J = 246.6, 7.0 Hz), 152.37 (dd, J = 240.8, 8.8 Hz), 151.30, 150.27, 148.38, 147.68, 143.55, 139.42, 138.47, 131.48, 131.12, 129.68, 129.10 – 128.66 (m), 128.63, 128.33, 127.57, 127.51, 125.96, 121.92 (dd, J = 13.7, 3.6 Hz), 118.60 – 117.95 (m), 117.57, 115.84, 115.59, 112.32 (dd, J = 23.0, 2.7 Hz), 68.91, 60.59, 58.81, 56.56, 55.86, 53.46, 52.75, 48.16, 41.67, 37.87, 35.80, 26.26, 16.83, 15.90, 12.61. LC-MS (ESI); m/z [M+H]^+^: Calcd. for C_51_H_58_F_2_N_9_O_7_S_2_, 1010.3868. Found 1010.4036.

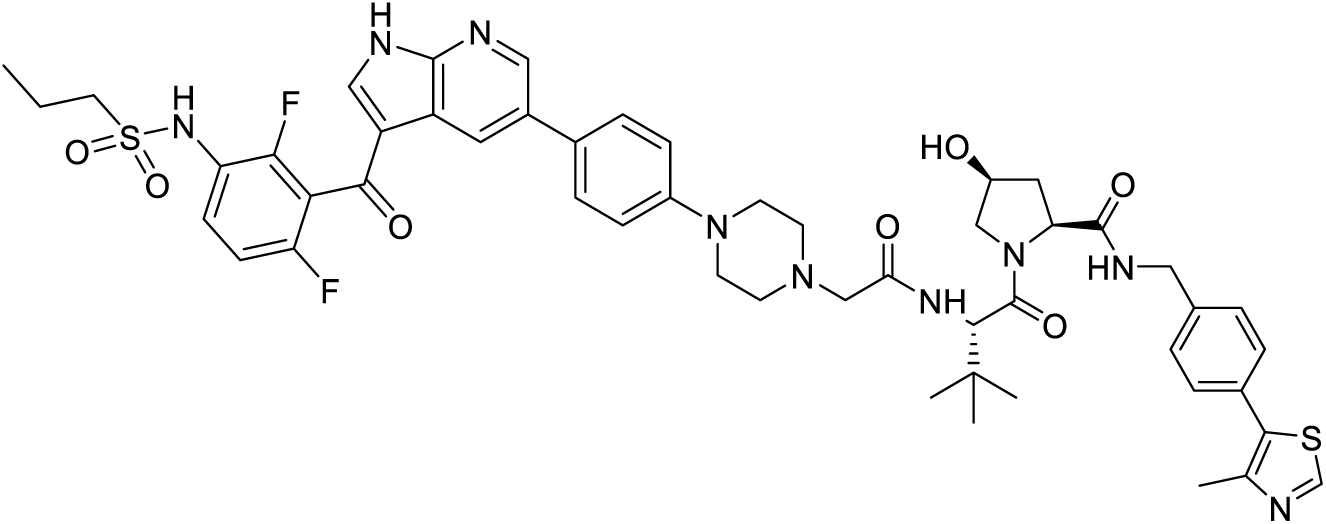

#### (2S,4S)-1-((S)-2-(2-(4-(4-(3-(2,6-difluoro-3-(propylsulfonamido)benzoyl)-1H-pyrrolo[2,3-b]pyridin-5-yl)phenyl)piperazin-1-yl)acetamido)-3,3-dimethylbutanoyl)-4-hydroxy-N-(4-(4-methylthiazol-5-yl)benzyl)pyrrolidine-2-carboxamide (SJF-0661)

A solution of tert-butyl 2-(4-(4-(3-(2,6-difluoro-3-(propylsulfonamido)benzoyl)-1H-pyrrolo[2,3-b]pyridin-5-yl)phenyl)piperazin-1-yl)acetate (**4**) (20.0 mg, 0.0306 mmol) in a mixture of TFA (1 mL) and Dichloromethane (2 ml) was stirred for 5 h. Then the solvent was removed under vacuum and crude product was dried under high vacuum for 1 h. Crude product was used in the next step without any further purification (5.7 mg, quantitative yield). HRMS (ESI); m/z: [M+H]+ Calcd. for C_29_H_30_F_2_N_5_O_5_S, 598.1935. Found 598.1755. To a solution of crude product from above; 2-(4-(4-(3-(2,6-difluoro-3-(propylsulfonamido)benzoyl)-1H-pyrrolo[2,3-b]pyridine-5-yl)phenyl) piperazin-1-yl)acetic acid (5.70 mg, 0.00954 mmol) and (2S,4S)-1-((S)-2-amino-3,3-dimethylbutanoyl)-4-hydroxy-N-(4-(4-methylthiazol-5-yl)benzyl)pyrrolidine-2-carboxamide (**6**) (5.35 mg, 0.0124 mmol) in DMF (1 ml) was added TEA (0.1 mL, 0.762 mmol) and PyBOP (5.96 mg, 0.0114 mmol) at room temperature. The reaction mixture was stirred for 4 h at the same temperature. TLC (DCM:MeOH:NH4OH, 90:9:1) shows no starting material (acid). The reaction mixture was evaporated to dryness under high vacuum. Crude product was filtered over a silica-carbonate cartridge (100 mg) using DCM:MeOH (9:1) as a eluent and evaporated under vacuum.. Crude product was purified by PTLC (DCM:MeOH:NH_4_OH, 90:9:1, 2×) to give 6.1 mg of product (63% yield). ^1^H NMR (500 MHz, DMSO-d6) δ 12.93 (bs, 1H), 9.76 (bs, 1H), 8.93 (s, 1H), 8.69 (t, J = 5.8 Hz, 1H), 8.66 (s, 1H), 8.56 (bs, 1H), 8.18 (s, 1H), 7.81 (d, J = 8.6 Hz, 1H), 7.65 – 7.50 (m, 3H), 7.51 – 7.32 (m, 4H), 7.28 (t, J = 8.7 Hz, 1H), 7.09 (d, J = 8.2 Hz, 2H), 5.48 (d, J = 7.2 Hz, 1H), 4.49 (d, J = 9.2 Hz, 1H), 4.46 – 4.33 (m, 2H), 4.32 – 4.18 (m, 2H), 3.96 – 3.88 (m, 1H), 3.54 – 3.42 (m, 1H), 3.30 – 3.19 (m, 4H), 3.17 – 3.00 (m, 4H), 2.78 – 2.56 (m, 4H), 2.41 (s, 3H), 2.39 – 2.30 (m, 1H), 1.82 – 1.68 (m, 3H), 0.98 (s, 9H), 0.95 (t, 3H). ^13^C NMR (151 MHz, DMSO-d6) δ 180.60, 172.29, 169.55, 168.84, 156.04 (dd, J = 246.5, 6.9 Hz), 152.38 (dd, J = 249.4, 8.3 Hz), 151.40, 150.29, 148.40, 147.75, 143.57, 139.16, 138.53, 131.50, 131.12, 129.79, 128.78 (d, J = 7.4 Hz), 128.70, 128.36, 127.54, 125.99, 121.93 (dd, J = 13.4, 3.6 Hz), 118.52 – 117.94 (m), 117.59, 115.87, 115.60, 112.35 (dd, J = 23.0, 3.9 Hz), 69.09, 60.50, 58.62, 56.01, 55.63, 53.46, 52.70, 48.17, 41.83, 36.90, 35.21, 26.25, 16.85, 15.93, 12.63 . LC-MS (ESI); m/z [M+H]^+^: Calcd. for C_51_H_58_F_2_N_9_O_7_S_2_, 1010.3868. Found 1010.3542.

**Scheme 2.**
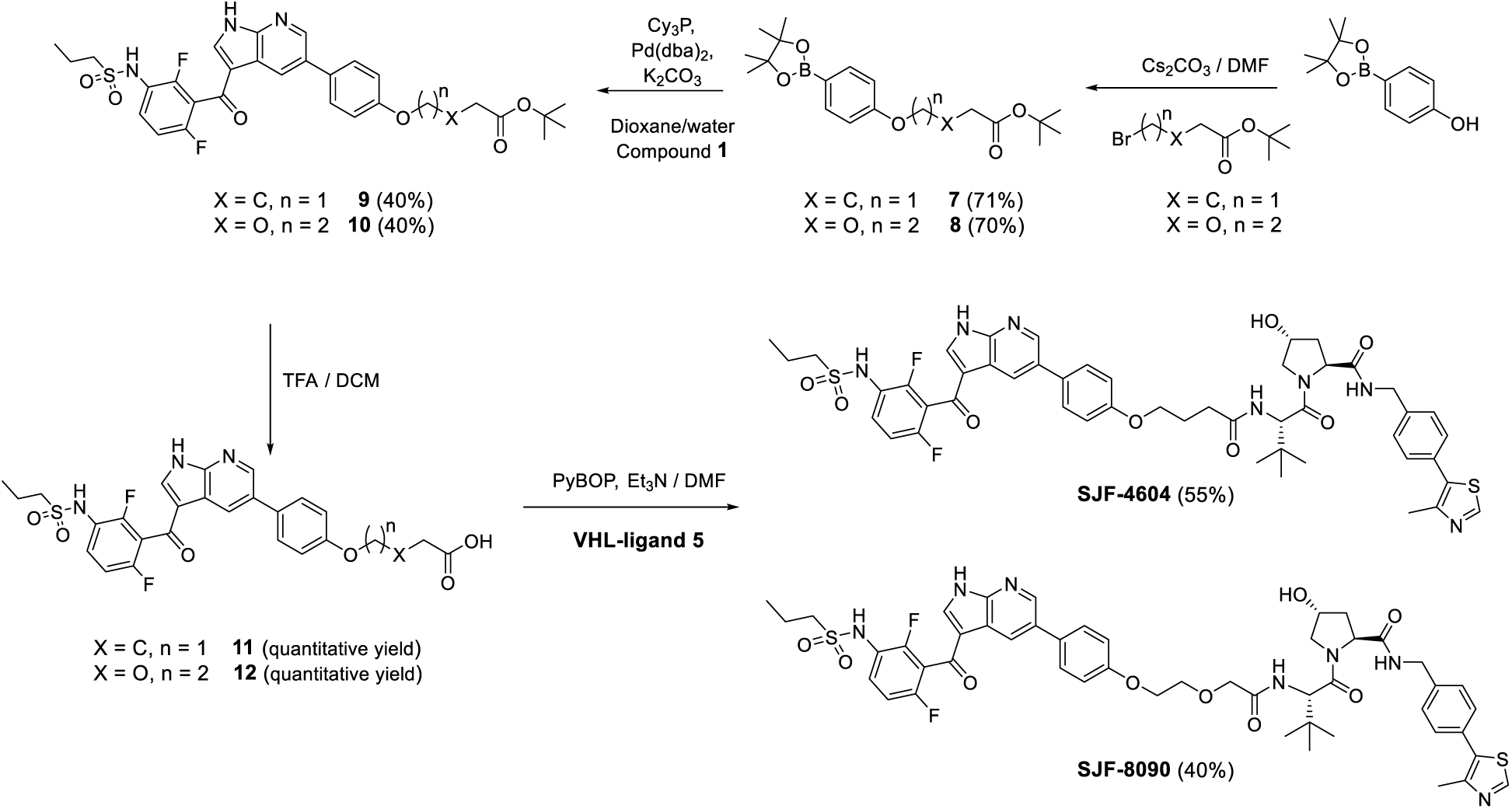
Synthetic Approach for SJF-4604 and SJF-8090

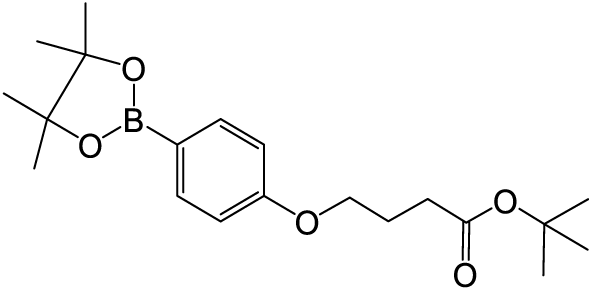

#### *tert*-Butyl 4-(4-(4,4,5,5-tetramethyl-1,3,2-dioxaborolan-2-yl)phenoxy)butanoate (7)

To a mixture of 4-(4,4,5,5-tetramethyl-1,3,2-dioxaborolan-2-yl)phenol (209.12 mg, 0.95 mmol) and *tert*-butyl 4-bromobutanoate (212 mg, 0.95 mmol) in N,N-Dimethylformamide (2 mL) was added Cs_2_CO_3_ (402.47 mg, 1.24 mmol). Reaction mixture was heated at 65 °C for 12 h (overnight). By TLC small amounts of starting material (Hex:EtOAc, 7:3). Reaction mixture was diluted with EtOAc (10 mL), washed with water (4×10 mL), dried (Na_2_SO_4_) and evaporated under vacuum. Crude product was purified by flash CC (SiO_2_-25g, Hex:EtOAc, gradient 9:1 to 4:6 in 15 min) to give 198 mg (57% yield) of product as an oil: ^1^H NMR (500 MHz, DMSO-d6) δ 7.59 (d, J = 8.2 Hz, 2H), 6.91 (d, J = 7.9 Hz, 2H), 3.99 (t, J = 6.3 Hz, 2H), 2.35 (t, J = 7.3 Hz, 2H), 1.92 (p, J = 6.7 Hz, 2H), 1.39 (s, 9H), 1.27 (s, 12H). ^13^C NMR (101 MHz, dmso) δ 172.25, 161.56, 136.66, 120.43, 114.37, 83.77, 80.12, 66.81, 31.72, 28.20, 25.12, 24.71. LC-MS (ESI); m/z [M+Na]^+^: C_20_H_31_BO_5_Na, 385.2162. Found 385.2194.

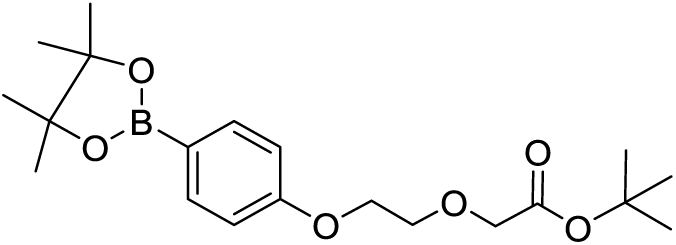

#### *tert*-Butyl 2-(2-(4-(4,4,5,5-tetramethyl-1,3,2-dioxaborolan-2-yl)phenoxy)ethoxy)acetate (8)

To a mixture of 4-(4,4,5,5-tetramethyl-1,3,2-dioxaborolan-2-yl)phenol (100 mg, 0.45 mmol) and *tert*-butyl 2-(2-bromoethoxy)acetate (108.65 mg, 0.91 mmol) in N,N-Dimethylformamide (2 mL) was added Cs_2_CO_3_ (296 mg, 0.86 mmol). Reaction mixture was heated at 60 °C for 2 h. By TLC no starting materia (less polar product, Hex:EtOAc, 7:3). Crude product was purified by flash CC (SiO_2_-25g, Hex:EtOAc, 9:1 to 4:6 in 15 min) to give 134 mg of product as a waxy solid (70% yield). ^1^H NMR (400 MHz, Chloroform-d) δ 7.74 (d, J = 8.5 Hz, 2H), 6.91 (d, J = 8.5 Hz, 2H), 4.19 (dd, J = 5.7, 3.8 Hz, 2H), 4.09 (s, 2H), 3.93 (dd, J = 5.8, 3.7 Hz, 2H), 1.48 (s, 9H), 1.33 (s, 12H). ^13^C NMR (151 MHz, cdcl3) δ 169.71, 161.35, 136.62, 121.07, 114.05, 83.70, 81.87, 69.94, 69.36, 67.37, 28.26, 25.01. HRMS (ESI); m/z: [M+Na]^+^ Calcd. for C_20_H_31_BO_6_Na, 401.2111. Found 401.2102.

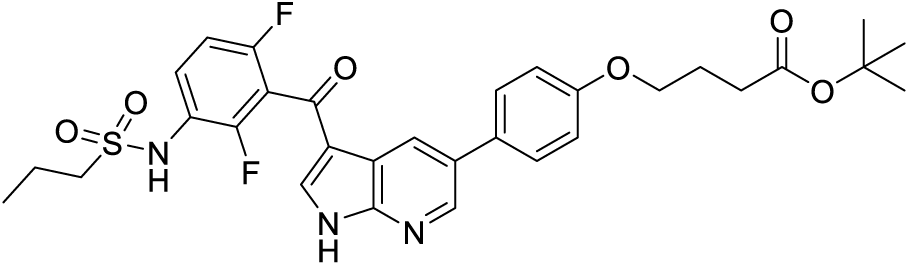

#### *tert*-Butyl 4-(4-(3-(2,6-difluoro-3-(propylsulfonamido)benzoyl)-1H-pyrrolo[2,3-b]pyridin-5-yl)phenoxy)-butanoate (9)

To a solution of *tert*-butyl 4-[4-(4,4,5,5-tetramethyl-1,3,2-dioxaborolan-2-yl)phenoxy] -butanoate (65 mg, 0.18 mmol) in Dioxane (3 ml) in a microwave vial was added N-[3-(5-bromo-1H-pyrrolo[2,3-b]pyridine-3-carbonyl)-2,4-difluoro-phenyl]propane-1-sulfonamide (82.2 mg, 0.18 mmol). Then the reaction mixture was de-gassed under vacuum and purged with argon (5×). Then Tricyclohexylphosphine (5.03 mg, 0.0179 mmol) and Pd(dba)2 (5.16 mg, 0.01 mmol) were added into and the vial was caped and sealed under a stream of argon, then the reaction mixture was heated at 100 °C in a microwave reactor for 2 h. By TLC no starting material (Hex:EtOAc, 3:7), the reaction mixture was filtered in vacuo over a celite pad. Filtrate was poured onto an aqueous saturated solution of NaCl (20 mL) and the product was extracted with EtOAc (2×20 mL). The EtOAc layers were combined, dried (Na_2_SO_4_) and concentrated in vacuo. The crude material was diluted in DCM and purified by flash chromatography (SiO_2_-12g, Hexane:EtOAc, 8:2 to 100% EtOAc in 15 min) to give 49 mg (40%) of product as a pale solid. ^1^H NMR (500 MHz, DMSO-d6) δ 12.93 (s, 1H), 9.74 (s, 1H), 8.64 (d, J = 2.0 Hz, 1H), 8.55 (bs, 1H), 8.19 (s, 1H), 7.65 (d, J = 8.5 Hz, 2H), 7.57 (td, J = 9.0, 5.6 Hz, 1H), 7.26 (t, J = 8.6 Hz, 1H), 7.05 (d, J = 8.6 Hz, 2H), 4.03 (t, J = 6.3 Hz, 2H), 3.15 – 3.06 (m, 2H), 2.38 (t, J = 7.3 Hz, 2H), 1.95 (p, J = 6.8 Hz, 2H), 1.78 – 1.67 (m, 2H), 1.39 (s, 9H), 0.94 (t, J = 7.4 Hz, 3H). ^13^C NMR (126 MHz, dmso) δ 181.02, 172.32, 158.70, 157.46, 157.41, 156.46 (d, J = 240.0 Hz), 152.76 (dd, J = 249.2, 8.3 Hz), 148.97, 144.17, 131.74, 130.92, 129.18 (d, J = 12.4 Hz), 128.71, 126.86, 122.34 (d, J = 13.2 Hz), 119.81 – 117.27 (m), 117.96, 116.05, 115.59, 112.76 (d, J = 27.7 Hz). 80.13, 67.09, 53.89, 31.80, 28.22, 24.81, 17.27, 13.05. LC-MS (ESI); m/z: [M+H]^+^ Calcd. for C_31_H_34_F_2_N_3_O_6_S, 614.2136. Found 614.2281.

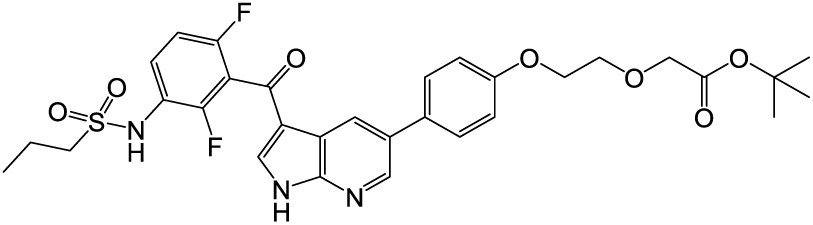

#### *tert*-Butyl 2-(2-(4-(3-(2,6-difluoro-3-(propylsulfonamido)benzoyl)-1H-pyrrolo[2,3-b]pyridin-5-yl)-phenoxy)ethoxy)acetate (10)

To a solution of *tert*-butyl 2-[2-[4-(4,4,5,5-tetramethyl-1,3,2-dioxaborolan-2-yl)phenoxy]ethoxy]acetate (54 mg, 0.14 mmol) and N-[3-(5-bromo-1H-pyrrolo[2,3-b]pyridine-3-carbonyl)-2,4-difluoro-phenyl]propane-1-sulfonamide (72 mg, 0.16 mmol) in Dioxane (6 ml) was degassed under vacuum and purged with argon (5×). Then K_2_CO_3_ (210 mg, 1.52 mmol) and Water (3 ml) was added. The reaction mixture was degassed again under vacuum and purged with argon (5×). Then Tricyclohexylphosphine (14.27 mg, 0.05 mmol) and Pd(dba)_2_ (14.6 mg, 0.025 mmol) were added into and degassed again under vacuum and purged with argon (5×). The reaction mixture was heated with vigorous stirring at 80 °C and stirred for 12 h. By TLC (DCM:MeOH:NH_4_OH, 90:9:1, 3X) full conversion (SM and product have a very similar r.f.). The reaction mixture was diluted with EtOAc (50 mL) and filtered over a celite pad. Filtrate was washed with an aqueous saturated solution of NaCl (30 mL) and product was extracted with EtOAc (2×50 mL). EtOAc layers combined, dried (Na_2_SO_4_) and concentrated under vacuo. Crude product was purified by flash CC (SiO_2_-12g, Hex:EtOAc, 1:9 to 100% in 15 min) to give 40 mg of product (40% yield). ^1^H NMR (400 MHz, DMSO-d6) δ 12.95 (bs, 1H), 9.77 (bs, 1H), 8.67 (s, 1H), 8.58 (s, 1H), 8.21 (s, 1H), 7.68 (d, J = 8.3 Hz, 2H), 7.59 (q, J = 8.8 Hz, 1H), 7.28 (t, J = 8.7 Hz, 1H), 7.09 (d, J = 8.3 Hz, 2H), 4.18 (dd, J = 5.6, 3.3 Hz, 2H), 4.08 (s, 2H), 3.84 (dd, J = 5.5, 3.4 Hz, 2H), 3.19 – 3.06 (m, 2H), 1.74 (h, J = 7.5 Hz, 2H), 1.43 (s, 9H), 0.96 (t, J = 7.4 Hz, 3H). ^13^C NMR (151 MHz, DMSO-d6) δ 180.61, 169.33, 158.18, 156.05 (d, J = 246.4 Hz), 152.31 (d, J = 249.5 Hz), 148.56, 143.75, 138.66, 131.30, 130.58, 128.76 (d, J = 9.8 Hz), 128.29, 126.47, 121.94 (dd, J = 13.6, 3.6 Hz), 118.23 (dd, J = 24.6, 22.3 Hz), 117.54, 115.63, 115.17, 112.34 (dd, J = 22.8, 3.9 Hz), 80.76, 69.07, 68.19, 67.18, 53.45, 27.77, 16.85, 12.62. LC-MS (ESI); m/z: [M+H]^+^ Calcd. for C_31_H_34_F_2_N_3_O_7_S, 630.2085. Found 630.5872.

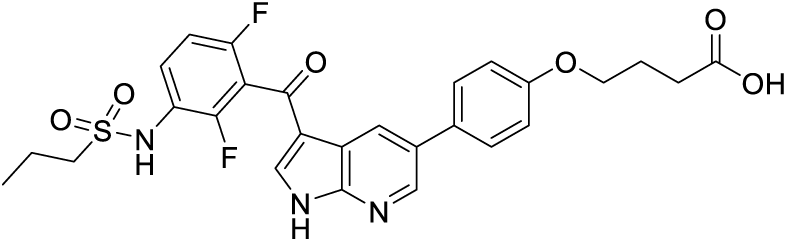

#### 4-(4-(3-(2,6-Difluoro-3-(propylsulfonamido)benzoyl)-1H-pyrrolo[2,3-b]pyridin-5-yl)phenoxy)butanoic acid (11)

A solution of *tert*-butyl 4-[4-[3-[2,6-difluoro-3-(propylsulfonylamino)benzoyl] -1H-pyrrolo[2,3-b]pyridin-5-yl]phenoxy]butanoate (16 mg, 0.03 mmol) in a mixture of TFA (1 ml, 13.46 mmol) and Dichloromethane (2 ml) was stirred at room temperature for 2 h. Then the solvent was removed under vacuum and crude product was dried under high vacuum for 2 h. Crude product was used in the next step without any further purification (14.5 mg, quantitative yield). LC-MS (ESI); m/z: [M+H]^+^ Calcd. for C_27_H_26_F_2_N_3_O_6_S, 558.1510. Found 558.1603.

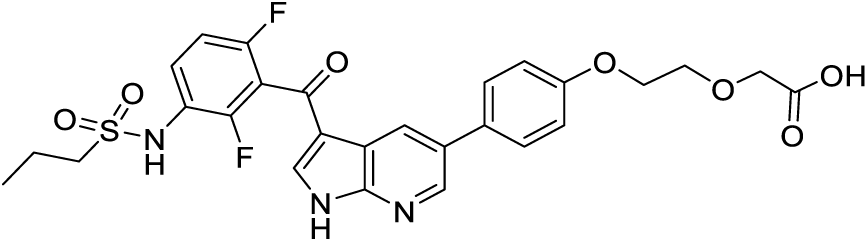

#### 2-(2-(4-(3-(2,6-Difluoro-3-(propylsulfonamido)benzoyl)-1H-pyrrolo[2,3-b]pyridin-5-yl)phenoxy)ethoxy) -acetic acid (12)

A solution of *tert*-butyl 2-[2-[4-[3-[2,6-difluoro-3-(propylsulfonylamino)benzoyl]-1H-pyrrolo[2,3-b]-pyridin-5-yl]phenoxy]ethoxy]acetate (8 mg, 0.013 mmol) in a mixture of TFA (1 ml, 13.46 mmol) and Dichloromethane (2 ml) was stirred at room temperature for 2 h. Then the solvent was removed under vacuum and crude product was dried under high vacuum for 2 h. Crude product was used in the next step without any further purification (7.2 mg, quantitative yield). LRMS (ESI); m/z: [M+H]^+^ Calcd. for C_27_H_26_F_2_N_3_O_7_S, 574.1459. Found 574.3837.

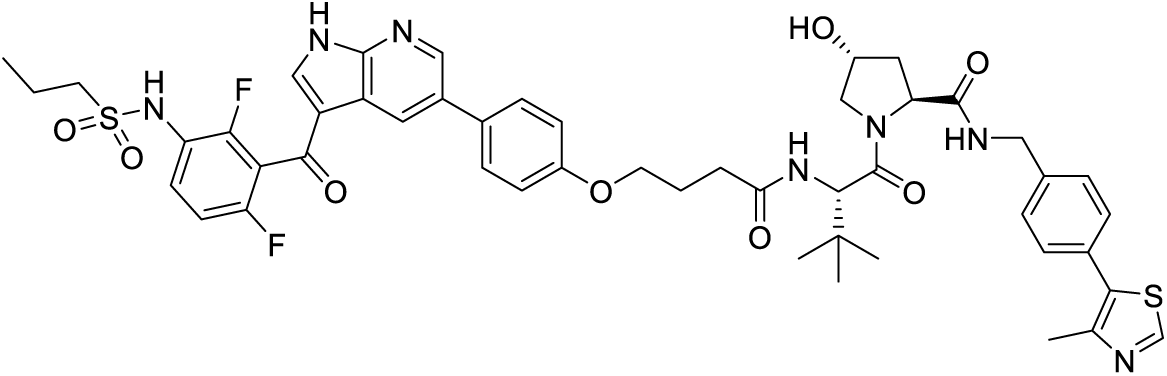

#### (2S,4R)-1-((S)-2-(4-(4-(3-(2,6-Difluoro-3-(propylsulfonamido)benzoyl)-1H-pyrrolo[2,3-b]pyridin-5-yl)phenoxy)butanamido)-3,3-dimethylbutanoyl)-4-hydroxy-N-(4-(4-methylthiazol-5-yl)benzyl) pyrrolidine-2-carboxamide (PROTAC SJF-4604)

To a solution of crude product from **11**; 4-[4-[3-[2,6-difluoro-3-(propylsulfonylamino)-benzoyl]-1H-pyrrolo[2,3-b]pyridin-5-yl]phen-oxy]butanoic acid (14.5 mg, 0.026 mmol) and **VHL-ligand 5** (2S,4R)-1-[(2S)-2-amino-3,3-dimethyl-butanoyl]-4-hydroxy-N-[[4-(4-methylthiazol-5-yl)-phenyl]methyl]pyrrolidine-2-carboxamide;hydrochloride (13 mg, 0.029 mmol) in DMF(2 ml) was added TEA (0.1 ml, 0.72 mmol) and PyBOP (14.9 mg, 0.029 mmol) at room temperature. The reaction mixture was stirred for 4 h at the same temperature. TLC (DCM:MeOH:NH_4_OH, 90:9:1) shows no starting materials. The DMF was removed under high vacuum. Crude product was filtered over a silica-carbonate cartridge using DCM:MeOH (9:1) as a eluent. Filtrate was evaporated under vacuum and crude product was purified by PTLC (DCM:MeOH:NH_4_OH, 90:9:1, 2×) to give 14 mg of product (55% yield). ^1^H NMR (400 MHz, DMSO-d6) δ 12.89 (bs, 1H), 9.72 (bs, 1H), 8.97 (s, 1H), 8.66 (d, J = 2.1 Hz, 1H), 8.58 (t, J = 5.7 Hz, 2H), 8.20 (s, 1H), 8.01 (d, J = 9.3 Hz, 1H), 7.66 (d, J = 8.3 Hz, 2H), 7.59 (td, J = 9.0, 5.8 Hz, 1H), 7.43 (d, J = 8.2 Hz, 2H), 7.38 (d, J = 8.2 Hz, 2H), 7.28 (t, J = 8.5 Hz, 1H), 7.07 (d, J = 8.7 Hz, 2H), 5.15 (d, J = 3.3 Hz, 1H), 4.58 (d, J = 9.3 Hz, 1H), 4.50 – 4.40 (m, 2H), 4.36 (bs, 1H), 4.22 (dd, J = 15.8, 5.3 Hz, 1H), 4.03 (t, J = 6.1 Hz, 2H), 3.76 – 3.61 (m, 2H), 3.17 – 3.05 (m, 2H), 2.44 (s, 3H), 2.49 – 2.31 (m, 2H), 2.13 – 1.85 (m, 4H), 1.74 (dq, J = 14.9, 7.4 Hz, 2H), 0.96 (s, 9H), 0.95 (t, 3H). ^13^C NMR (151 MHz, dmso) δ 181.03, 172.39, 172.03, 170.09, 156.43 (dd, J = 246.4, 6.9 Hz), 158.76, 152.75 (dd, J = 249.5, 8.5 Hz), 151.86, 148.95, 148.13, 144.17, 139.06, 131.76, 131.59, 130.81, 130.06, 129.18 (d, J = 14.4 Hz), 129.06, 128.69, 127.85, 126.86, 122.39 (dd, J = 13.8, 3.2 Hz), 118.,94 – 118.29 (m), 117.95, 116.04, 115.61, 112.75 (dd, J = 22.5, 3.3 Hz), 69.33, 67.55, 59.15, 56.90, 56.84, 53.87, 42.08, 38.40, 35.68, 31.74, 26.83, 25.48, 17.27, 16.38, 13.04. LC-MS (ESI); m/z [M+H]^+^: Calcd. for C_49_H_54_F_2_N_7_O_8_S_2_, 970.3443. Found 970.3176.

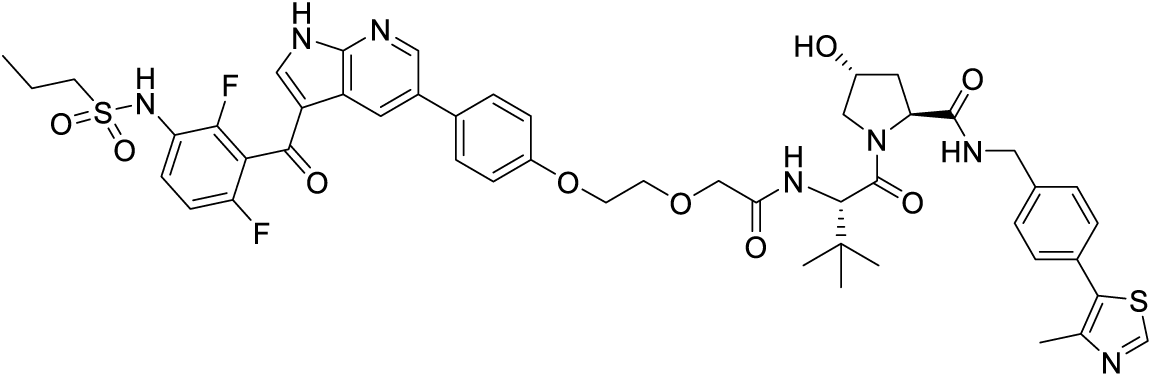

#### (2S,4R)-1-((S)-2-(2-(2-(4-(3-(2,6-Difluoro-3-(propylsulfonamido)benzoyl)-1H-pyrrolo[2,3-b]pyridin-5-yl)phenoxy)ethoxy)acetamido)-3,3-dimethylbutanoyl)-4-hydroxy-N-(4-(4-methylthiazol-5-yl)benzyl)-pyrrolidine-2-carboxamide (PROTAC SJF-8090)

To a solution of crude product from **12**; 2-[2-[4-[3-[2,6-difluoro-3-(propylsulfonylamino)benzoyl]-1H-pyrrolo-[2,3-b]pyridin-5-yl]phenoxy]ethoxy]acetic acid (7.28 mg, 0.01 mmol) and **VHL-ligand 5** (2S,4R)-1-[(2S)-2-amino-3,3-dimethyl-butanoyl]-4-hydroxy-N-[[4-(4-methylthiazol-5-yl)phenyl]methyl]pyrrolidine-2-carboxamide; hydrochloride (8.89 mg, 0.02 mmol) in DMF(2 ml) was added TEA (0.05 ml, 0.34 mmol) and PyBOP (7.93 mg, 0.02 mmol) at room temperature. The reaction mixture was stirred for 2 h at the same temperature. TLC (DCM:MeOH:NH_4_OH, 90:9:1) shows no starting materials. Reaction mixture was diluted with EtOAc (10 mL), washed with water (3×10 mL), dried (Na_2_SO_4_) and evaporated under vacuum to give 1 mg of crude product (product is partially soluble in water). Additional water extractions with EtOAc (5×30 mL) were performed. Organic extracts combined, dried (Na_2_SO_4_), and evaporated under high vacuum. Crude product was purified by PTLC (DCM:MeOH:NH_4_OH, 90:9:1, 2×) to give 5 mg of product (40 % total yield).^1^H NMR (500 MHz, DMSO-d6) δ 12.92 (bs, 1H), 9.73 (bs, 1H), 8.88 (s, 1H), 8.69 – 8.55 (m, 2H), 8.54 (bs, 1H), 8.19 (s, 1H), 7.67 – 7.54 (m, 3H), 7.53 (d, J = 9.6 Hz, 1H), 7.42 (d, J = 8.1 Hz, 2H), 7.36 (d, J = 8.1 Hz, 2H), 7.28 (t, J = 8.8 Hz, 1H), 7.16 (d, J = 8.7 Hz, 2H), 5.16 (d, J = 3.5 Hz, 1H), 4.62 (d, J = 9.6 Hz, 1H), 4.52 – 4.33 (m, 3H), 4.32 – 4.13 (m, 3H), 4.08 (s, 2H), 3.88 (t, J = 4.3 Hz, 2H), 3.76 – 3.57 (m, 2H), 3.19 – 3.03 (m, 2H), 2.37 (s, 3H), 2.11 – 1.99 (m, 1H), 1.97 – 1.84 (m, 1H), 1.81 – 1.64 (m, 2H), 0.97 (s, 9H), 0.96 (t, 3H). ^13^C NMR (151 MHz, dmso) δ 181.02, 172.25, 169.56, 168.95, 158.66, 151.69, 156.43 (dd, J = 246.3, 7.3 Hz), 152.75 (dd, J = 249.6, 8.6 Hz), 148.95, 148.09, 144.13, 139.84, 139.24 – 138.83 (m), 131.68, 131.50, 131.04, 130.07, 129.30, 129.08, 128.65, 127.87, 126.90, 122.37 (d, J = 15.4 Hz), 118.71 (d, J = 23.4 Hz), 117.93, 116.05, 115.69, 112.75 (dd, J = 23.3, 3.7 Hz), 70.02, 69.96, 69.33, 67.45, 59.18, 57.04, 56.15, 53.86, 42.14, 38.34, 36.28, 26.66, 17.27, 16.29, 13.04. LC-MS (ESI); m/z [M+H]^+^: Calcd. for C_49_H_54_F_2_N_7_O_9_S_2_, 986.3392. Found 986.3481.

---

**Extended Data Figure 1.**
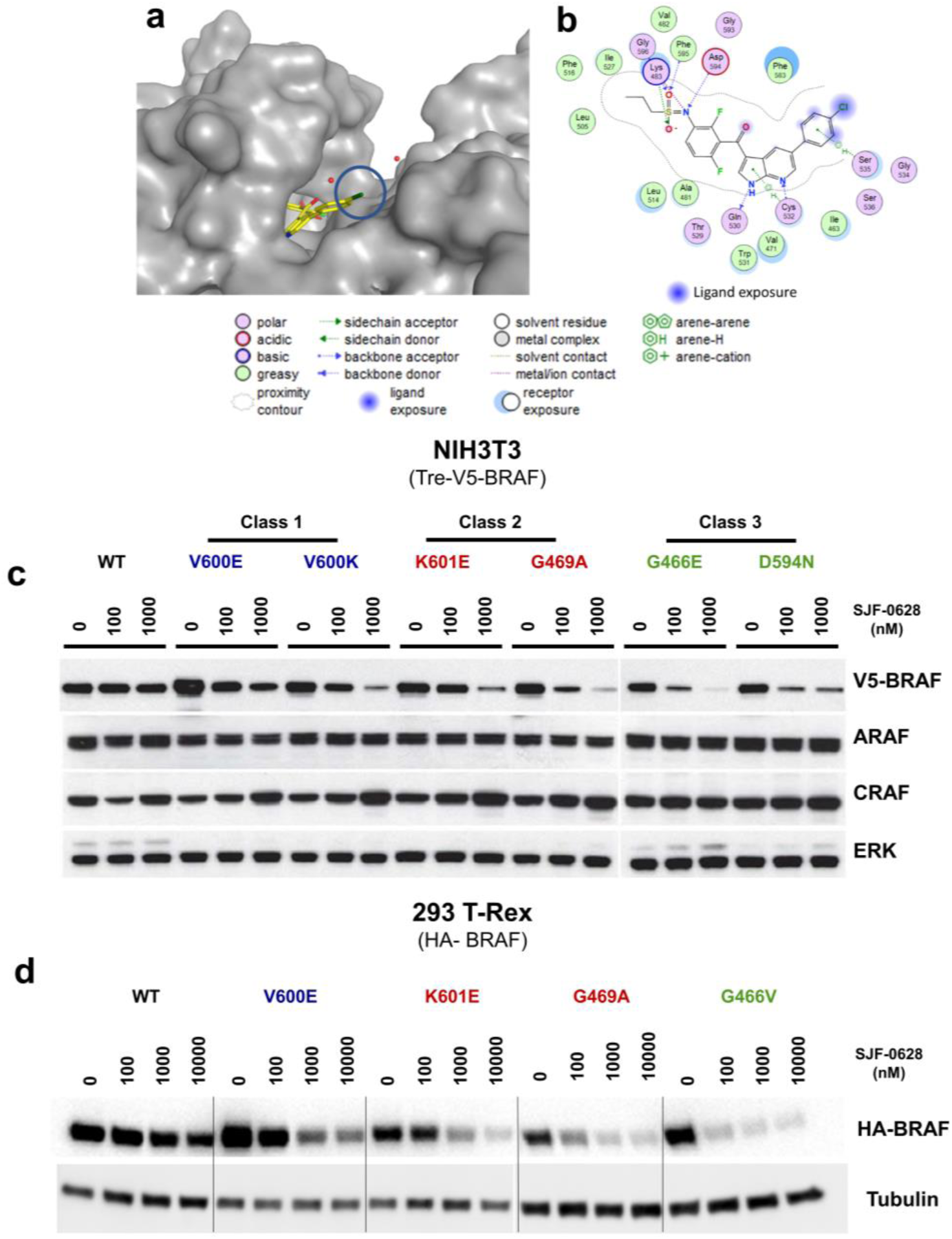
Vemurafenib based PROTAC, SJF-0628, induces mutant selective degradation of BRAF. **a,** Crystal structure of BRAF^V600E^ in complex with vemurafenib (PDB: 3OG7) **b**, Ligand interactions diagram showing important BRAF: vemurafenib interactions and solvent exposure. **c**, Inducible NIH3T3 cells expressing indicated V5-BRAF (doxycycline 500 ng/mL, 24 hours) treated with increasing amounts of SJF-0628 for 24 hours. **d**, 293 T-Rex cells (doxycycline 20 ng/mL, 24 hours) expressing indicated BRAF isoforms treated with increasing concentrations of SJF-0628 shows mutant selective degradation.

**Extended Data Figure 2.**
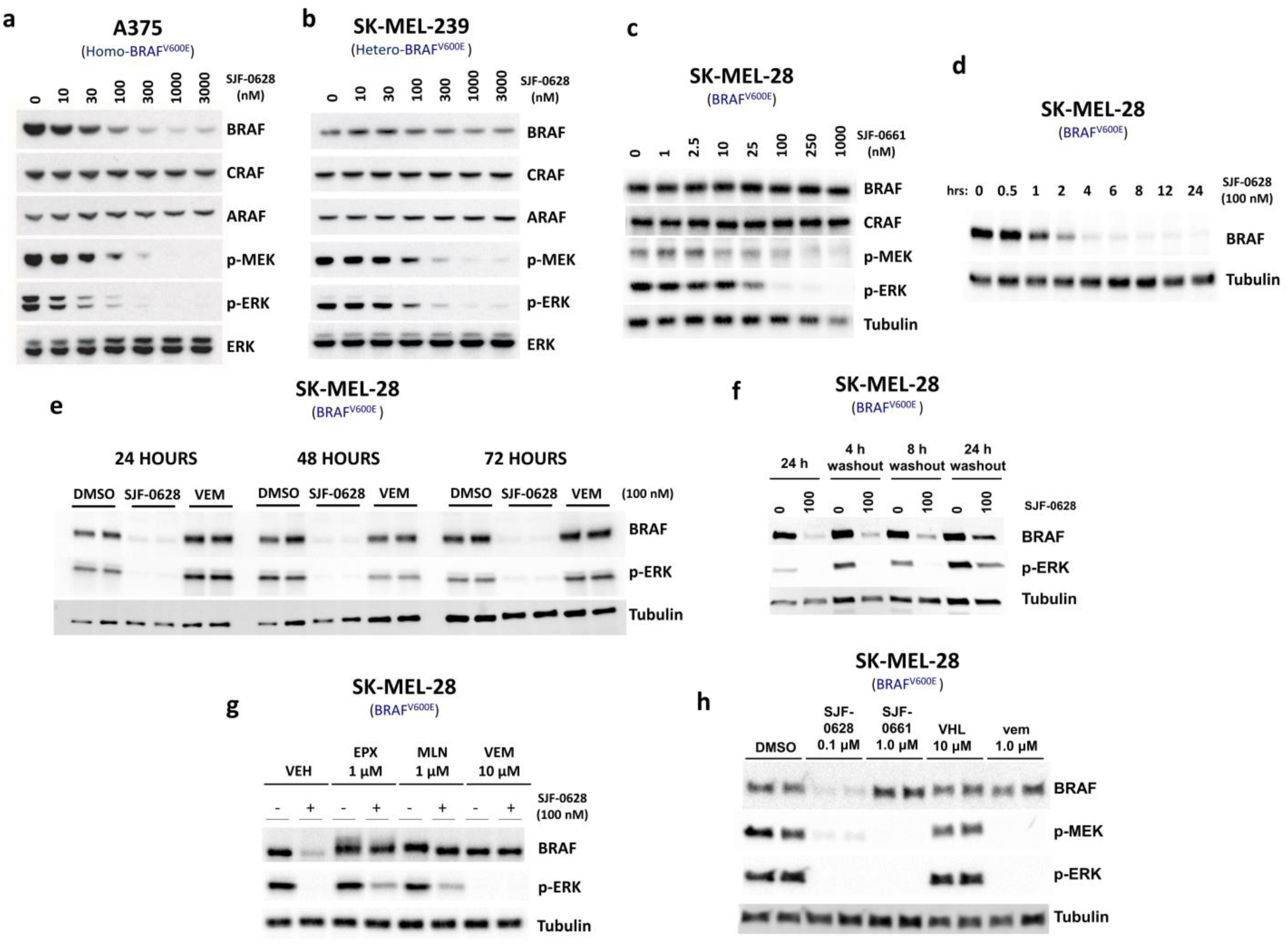
SJF-0628 induces a sustained and efficient degradation of BRAF^V600E^ via the proteasome. **a**, Treatment of A375 (homozygous BRAF^V600E^) cells with SJF-0628 shows BRAF^V600E^ degradation and inhibition of MEK and ERK phosphorylation. **b**, Treatment of SK-MEL-239 (heterozygous BRAF^V600E^) cells with SJF-0628 shows minimal BRAF degradation but marked inhibition of MEK and ERK phosphorylation. **c**,SK-MEL-28 cells treated with negative control epimer, SJF-0661, does not affect BRAF levels but inhibits ERK signaling. **d**, Representative immunoblot of SJF-0628 time course (100 nM) at indicated times in SK-MEL-28 cells (plotted in Fig. 1e) shows maximal degradation within 4 hours. **e**, Treatment of SK-MEL-28 cells with 100 nM of SJF-0628, and vemurafenib for indicated times shows sustained degradation and inhibition of MAPK signaling. **f**, SK-MEL-28 cells were treated with 100 nM for 24 hours, washed 3 times with DPBS and replenished with fresh media. Cells were then lysed either 4, 8 or 24 hours after media removal to determine level of BRAF and MAPK recovery. **g**, SK-MEL-28 cells treated with a proteasome inhibitor (EPX = epoxomicin), a neddylation inhibitor (MLN = MLN4924), or excess vemurafenib (VEM) for 2 hours, then subsequently treated with DMSO or PROTAC for 8 hours. **h**, SK-MEL-28 cells treated with indicated compound for 6 hours. VHL ligand alone does not cause MAPK inhibition.

**Extended Data Figure 3.**
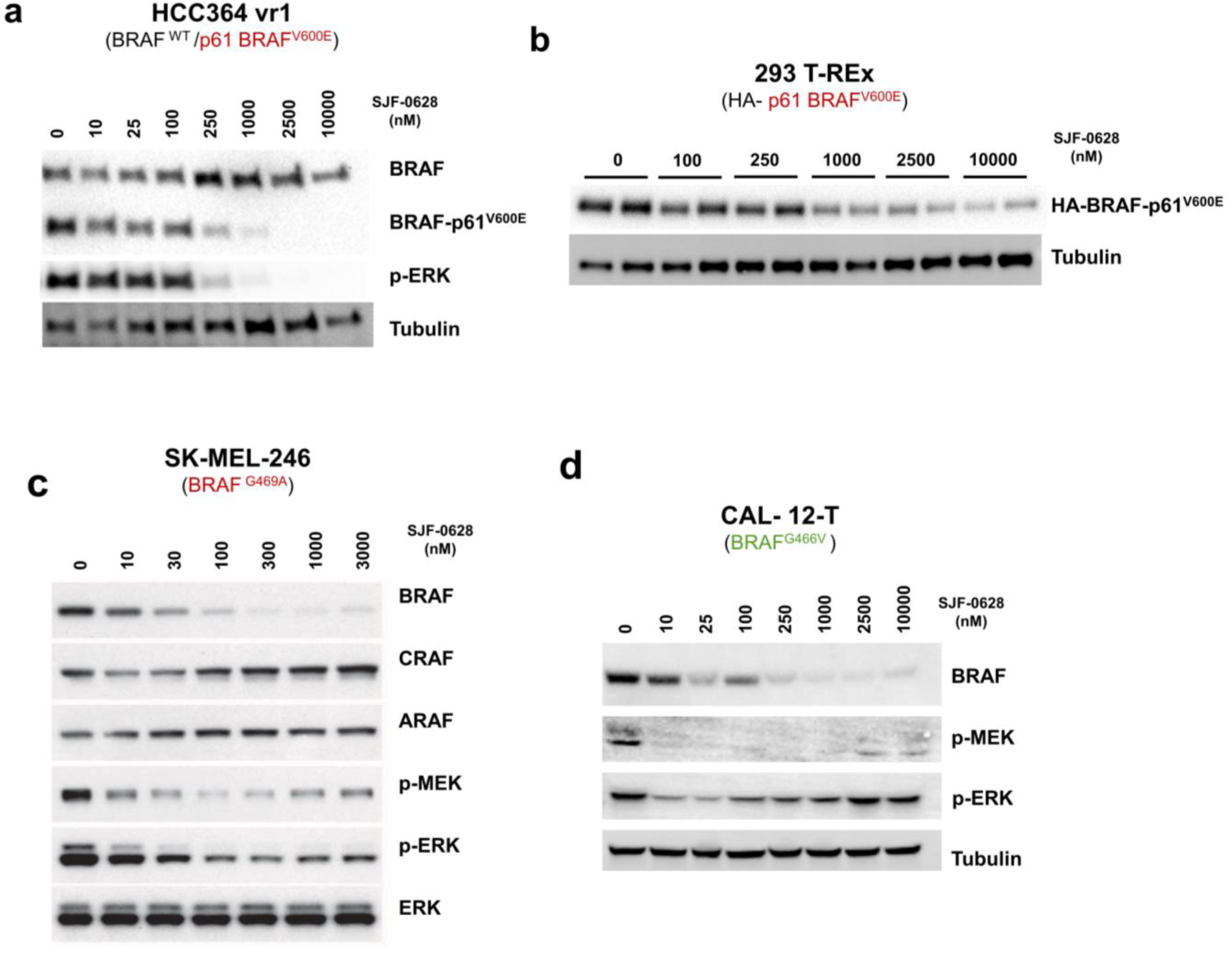
SJF-0628 induces degradation of acquired and intrinsic vemurafenib-resistant BRAF mutants. **a**, HCC364 vr1 (BRAF^WT^, p61-BRAF^V600E^) selectively induces degradation of p61-BRAF^V600E^ and spares BRAF^WT^. **b**, SJF-0628 treatment in 293 T-Rex cells expressing p61-BRAF^V600E^ shows dose dependent decrease in HA-p61^V600E^ protein levels. **c**, SK-MEL-246 (Class 2, BRAF ^G469A^) cells treated with increasing amount of SJF-0628 shows degradation of BRAF and inhibition of ERK signaling. **d**, CAL-12-T cells (homozygous BRAF^G466V^) treated with SJF-0628 shows BRAF degradation, but incomplete suppression of ERK signaling.

**Extended Data Figure 4.**
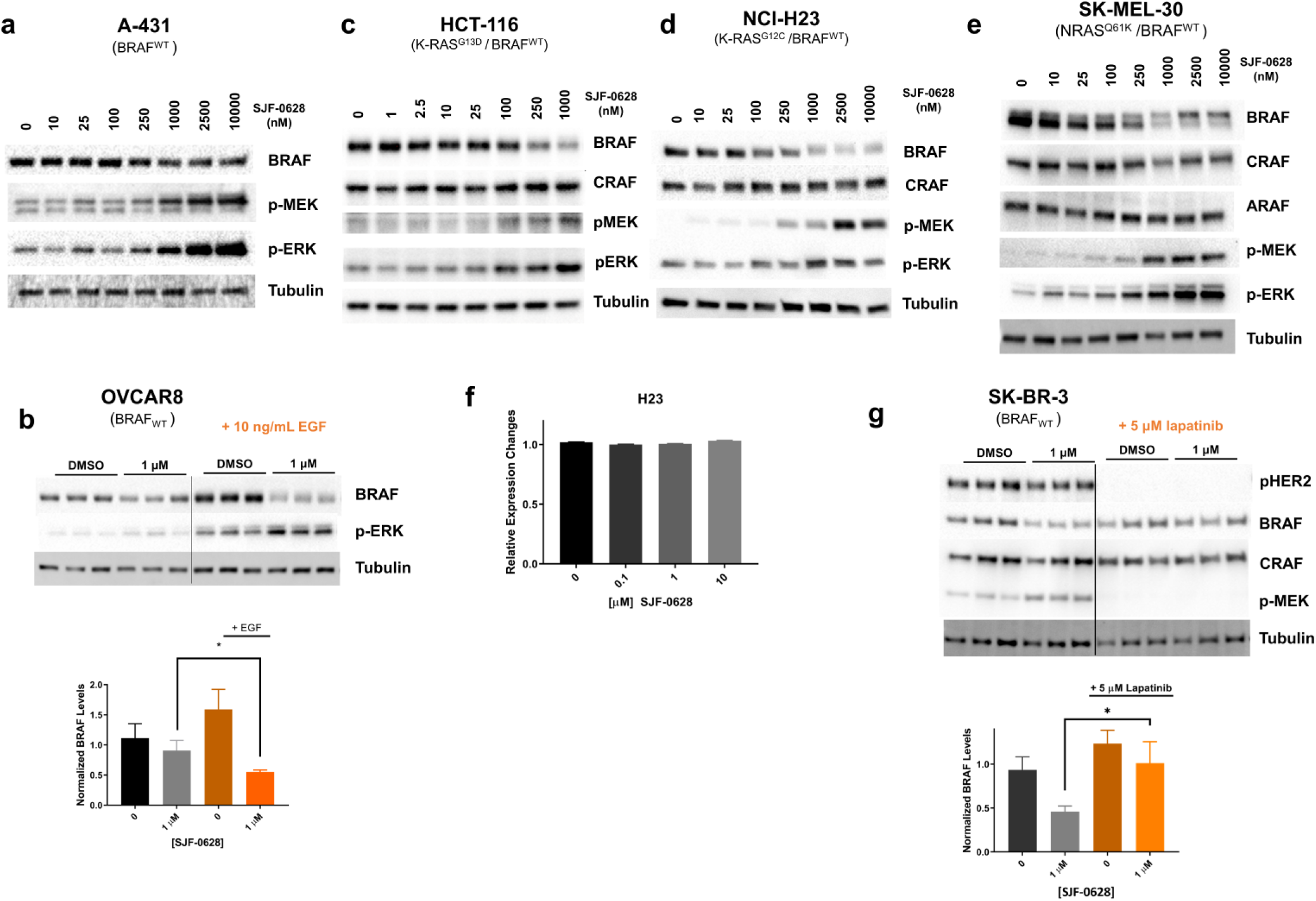
BRAF^WT^ is sensitized to PROTAC induced degradation in the presence of upstream drivers. **a**, Treatment of A-431 cells (HER1 amplification) which expresses BRAF^WT^ and RAS^WT^ treated with increasing amounts of SJF-0628 show some degradation of BRAF^WT^(∼30%). **b**, Serum-starved OVCAR8 cells stimulated with 10 ng/mL of EGF promotes SJF-0628 induced degradation of BRAF^WT^ **c**-**e**, SJF-0628 treatment in HCT116, H23, and SK-MEL-30 cells bearing a RAS mutation shows 40%-60% degradation of BRAF^WT^ at high concentrations and paradoxical activation of MAPK signaling. **f,** Quantitative real time PCR of H23 cells treated with SJF-0628 for 20 hours (mean ± s.d.,n=3). **g**, Lapatinib treatment in SKBR3 cells hinders PROTAC induced degradation (\**P* value <0.05). *P* value calculated by unpaired t-test.

**Extended Data Figure 5.**
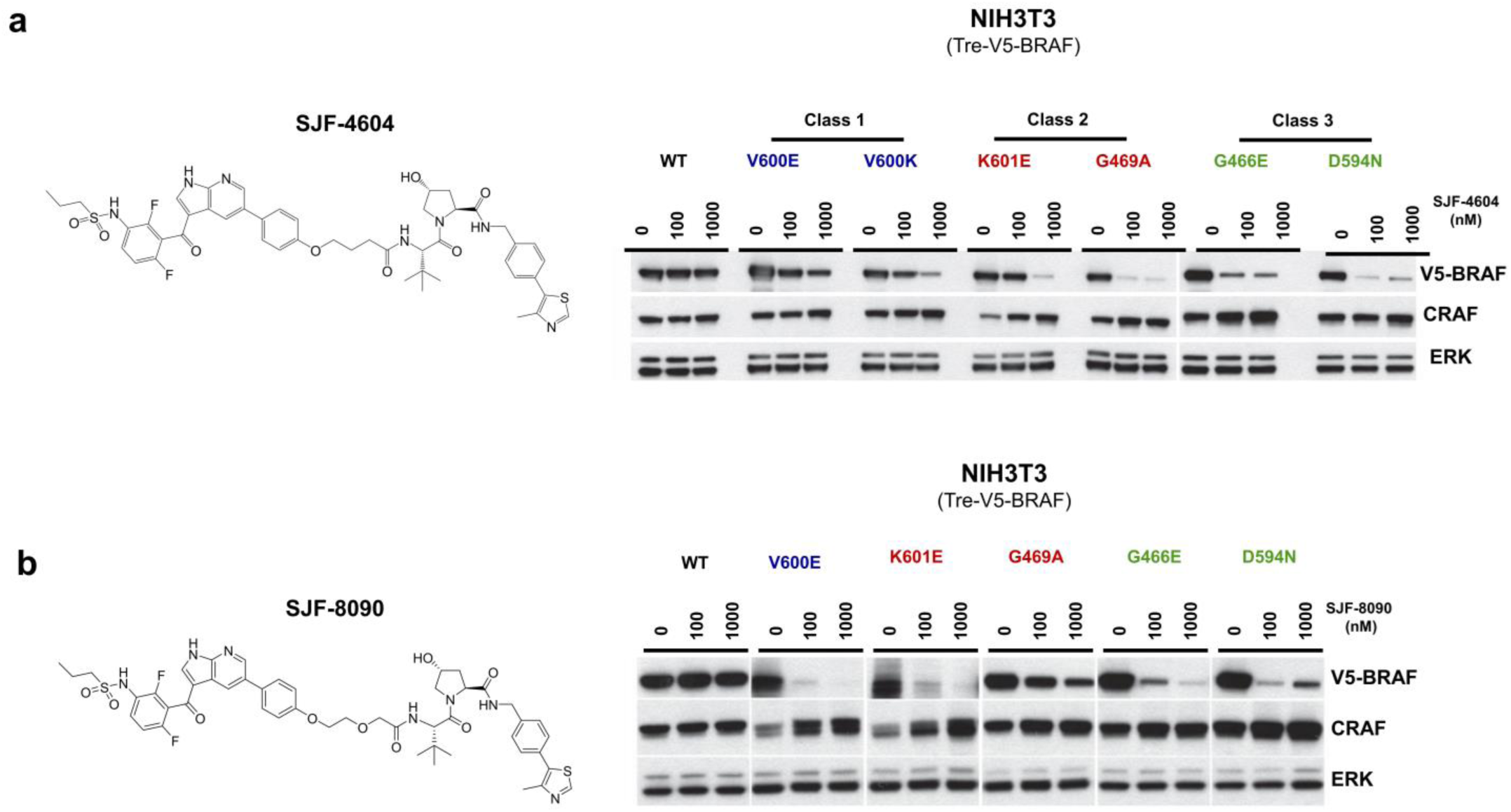
Vemurafenib based PROTACs spare BRAF^WT^ despite linker length and composition. **a-b**, Structures and results of vemurafenib based PROTACs SJF-4604 and SJF-8090 treatment in inducible NIH3T3 cells expressing indicated BRAF protein for 24 hours. Both PROTACs show mutant selective degradation.

**Extended Data Figure 6.**
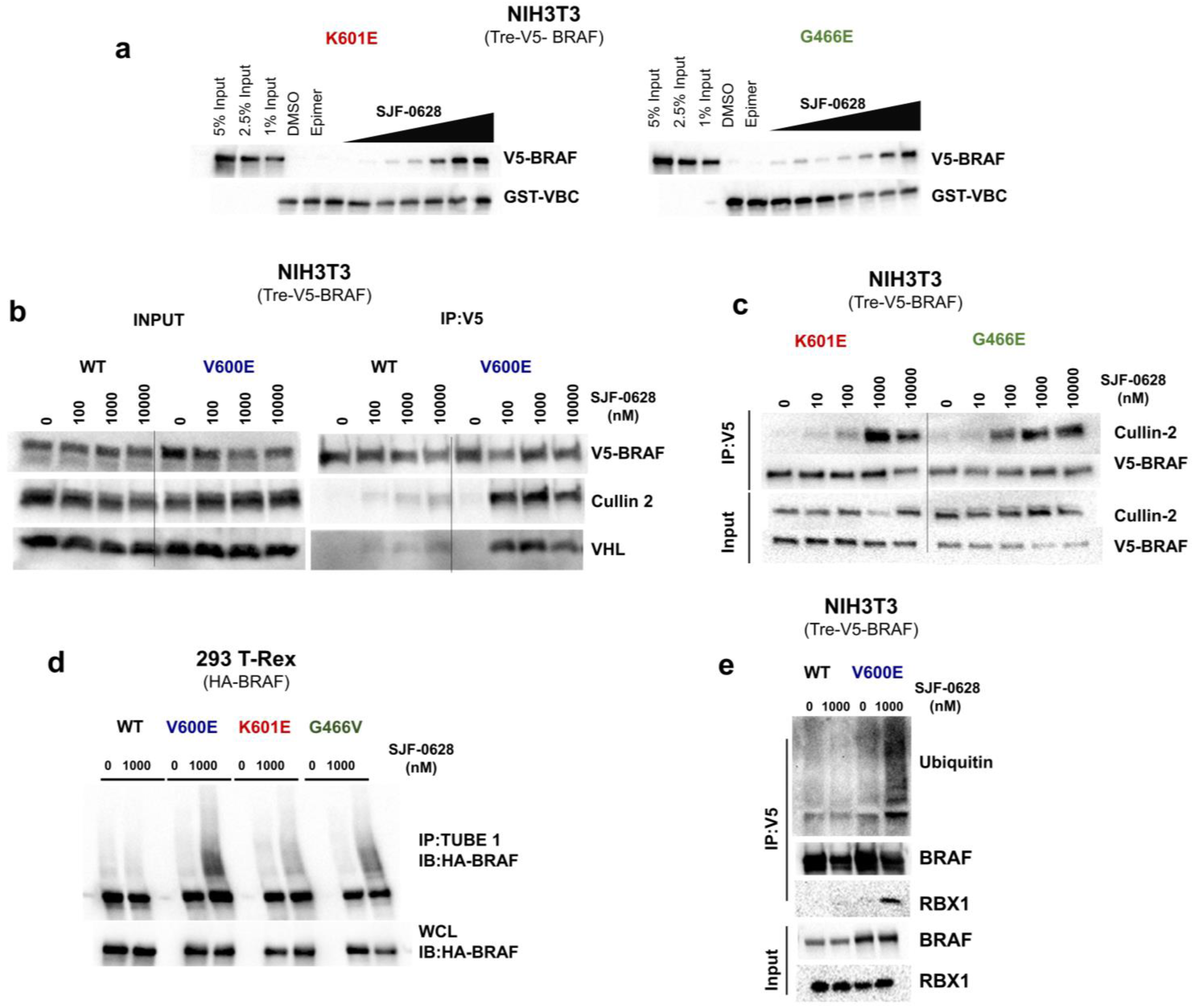
All three mutant BRAF classes bind SJF-0628 and form a stable ternary complex which promotes its ubiquitination. **a**, Cell lysate-based trimer assay of VBC immobilized on glutathione beads incubated with NIH3T3 cell lysates expressing BRAF^K601E^ and BRAF^G466E^ with vehicle, 500 nM SJF-0661, or increasing amounts of SJF-0628. **b**, V5-BRAF immunoprecipitation in NIH3T3 cells (WT and V600E) stably expressing human VHL treated with increasing concentrations SJF-0628 for 45 mins. **c,** V5-BRAF immunoprecipitation in NIH3T3 cells expressing BRAF^K601E^ or BRAF^G466E^ V600E treated with SJF-0628 for 1 hour. **d**, Tandem Ubiquitin Binding Entities 1 (TUBE1) Assay. Pulldown of tetra-ubiquitinated protein in 293 T-REx cells expressing indicated BRAF treated with SJF-0628 for 1 hour. Immunoblot for HA-BRAF. **e**, V5-BRAF immunoprecipitation in NIH3T3 cells expressing BRAF^WT^ and BRAF^V600E^ treated with SJF-0628 for 1 hour.

**Extended Data Figure 7.**
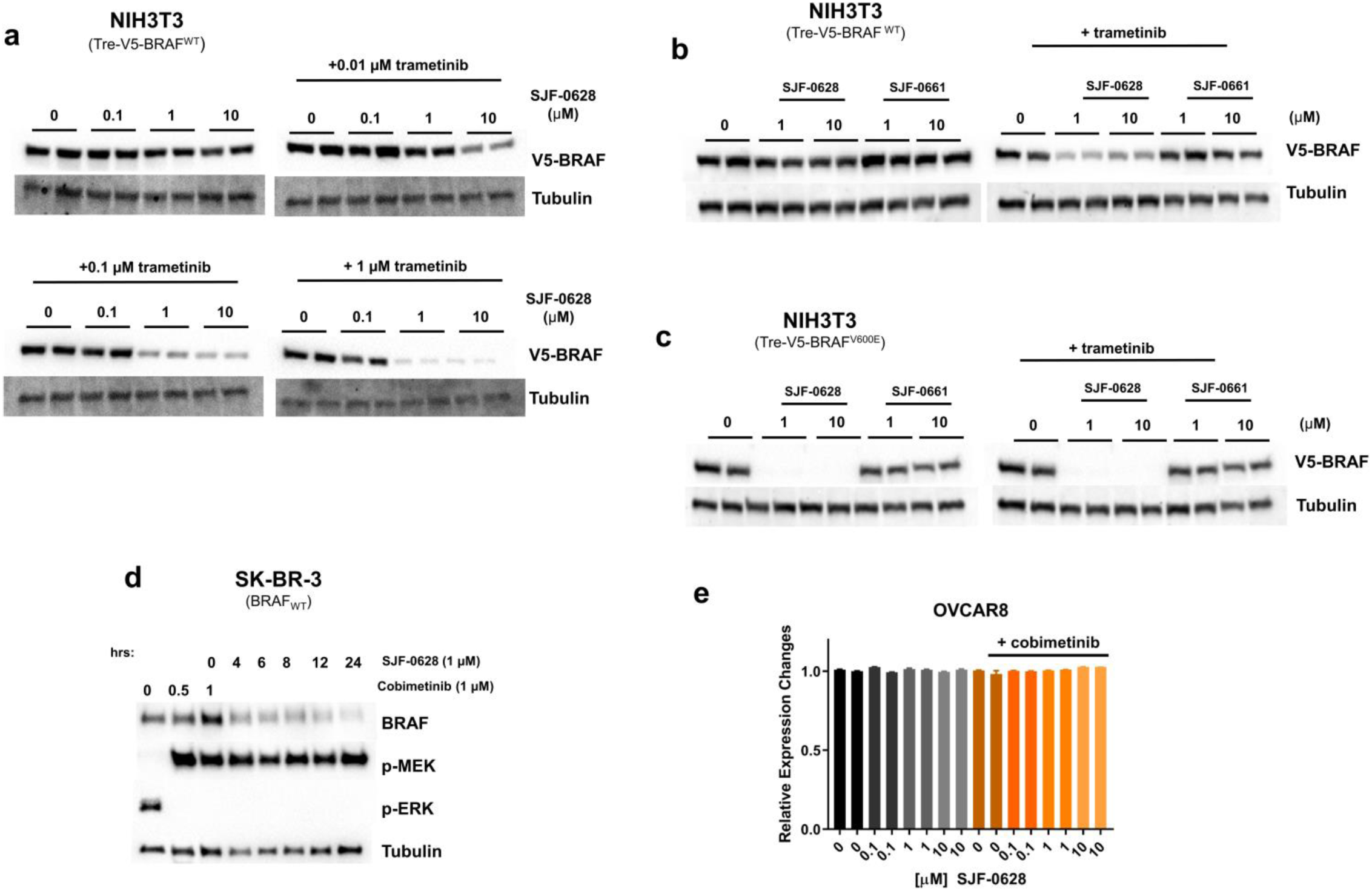
Trametinib or cobimetinib pre-treatment promotes dose and time dependent degradation of BRAF^WT^. **a**, NIH3T3 cells expressing inducible V5-BRAF^WT^ pre-treated with increasing concentrations of trametinib and subsequently treated with increasing concentrations of SJF-0628. **b-c**, NIH3T3 cells expressing BRAF^WT^ and BRAF^V600E^ pre-treated with trametinib followed by SJF-0628 or SJF-0661 treatment. **d,** Time course of SK-BR-3 cells pre-treated with 1 μM of cobimetinib for 1 hour then treated with 1 μM SJF-0628. **e**, mRNA expression changes of BRAF pre-treated with 1 μM of cobimetinib for 1 hour, then treated with 1 μM SJF-0628 for 20 hours(mean ± s.d.,n=3).

**Extended Data Figure 8.**
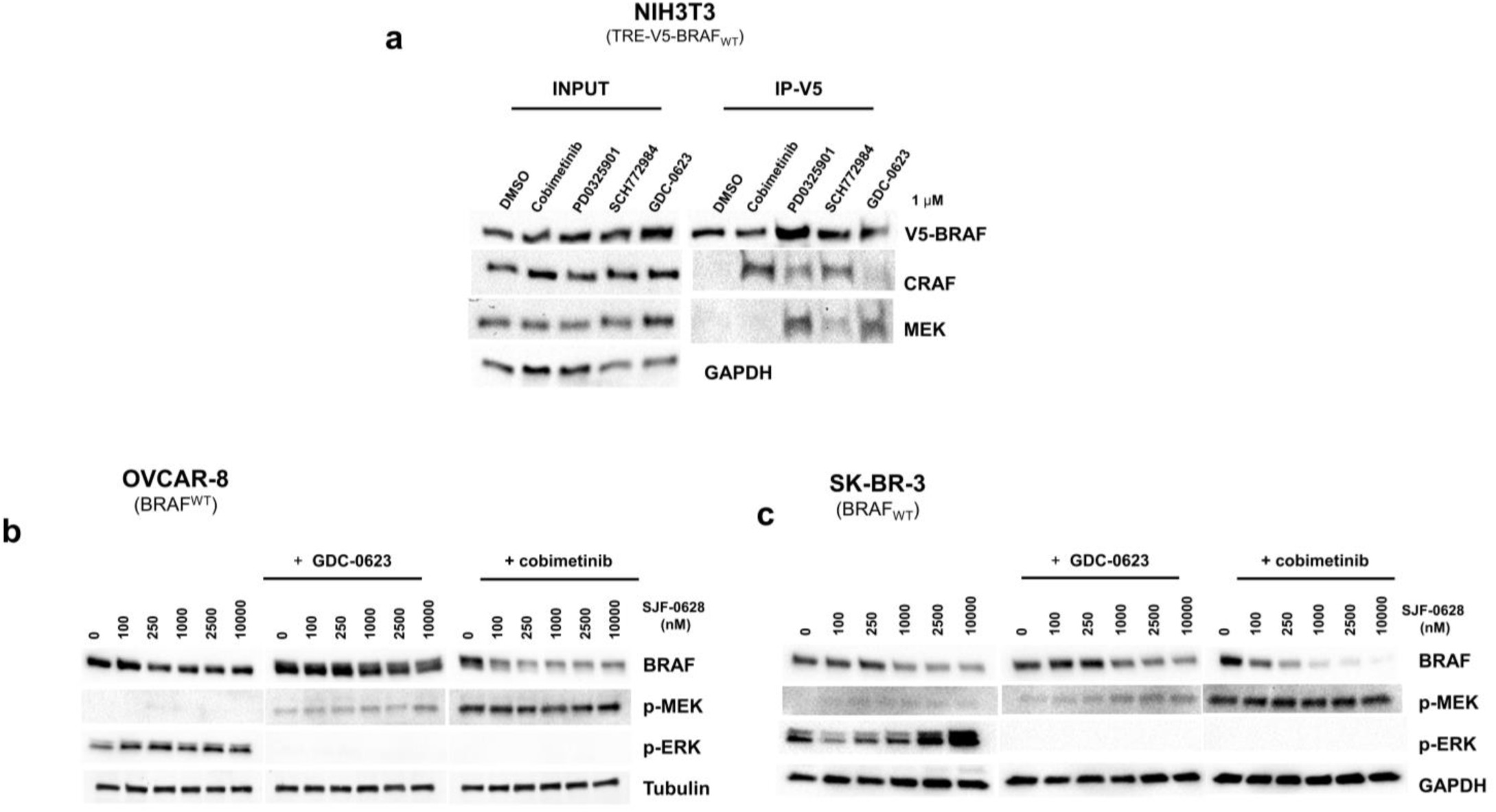
MAPK inhibitors that increase BRAF kinase activity promote SJF-0628 induced BRAF^WT^ degradation. **a**, Immunoprecipitation of V5-BRAF^WT^ from NIH3T3 cells treated with 1uM of the indicated MAPK pathway inhibitor. Cobimetinib treatment showed minimal BRAF association with MEK, but increased RAF dimerization while GDC-0623 showed minimal RAF dimerization, and increased BRAF: MEK association. **b-c**, OVCAR-8 cells and SK-BR-3 cells pre-treated with GDC-0623 and cobimetinib (500nM, 3 hours) then treated with SJF-0628 for 20 hours.

**Extended Data Figure 9.**
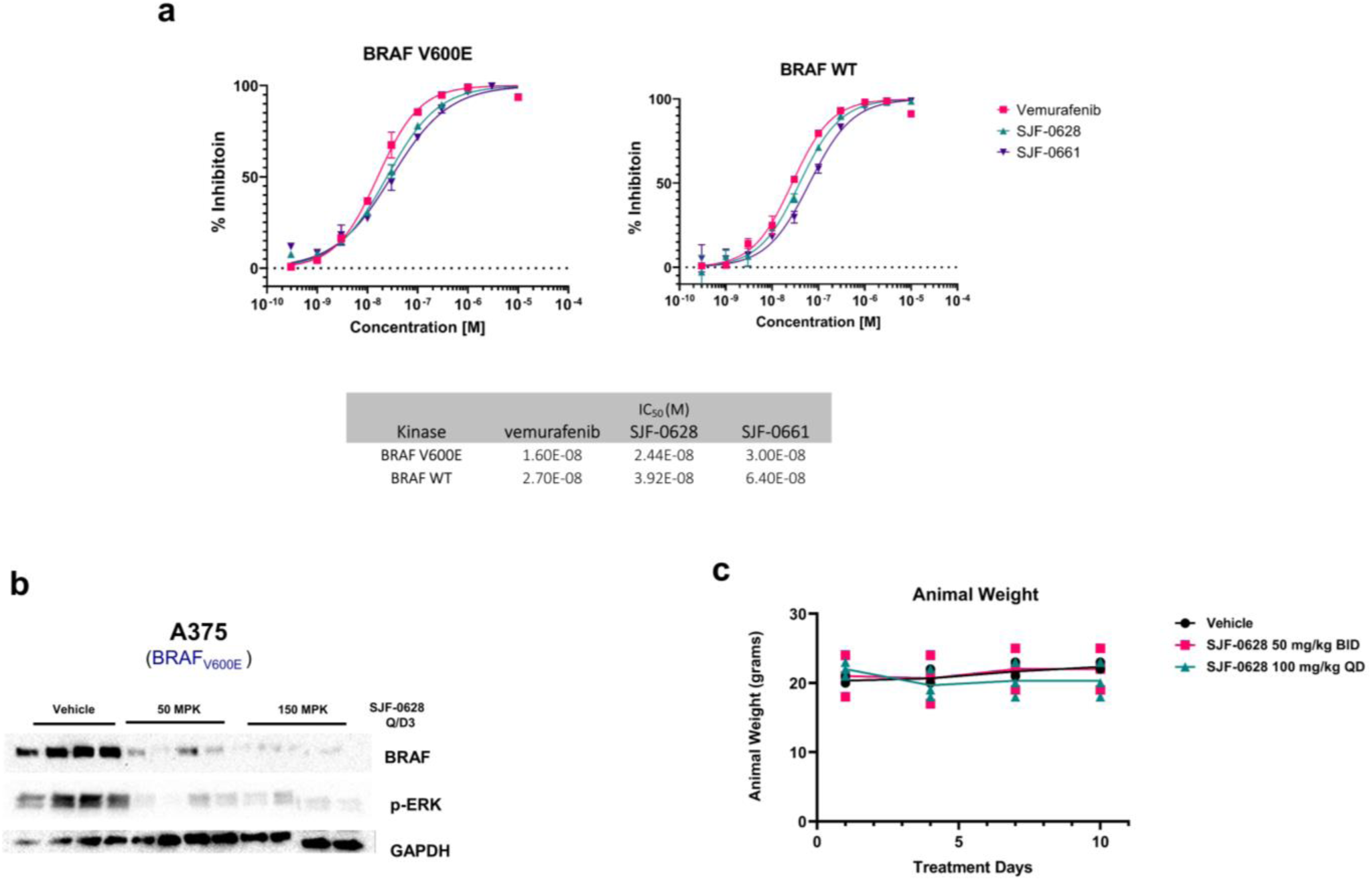
SJF-0628 inhibits BRAF^WT^ and BRAF^V600E^ with a similar affinity and induces degradation of BRAF^V600E^ *in vivo*. **a**, BRAF^WT^ and BRAF^V600E^ binding affinity and curves for ELISA kinase inhibition assay with SJF-0628, SJF-0661 and vemurafenib (n=2). **b,** BRAF^V600E^ degradation in A375 xenograft in female Balb/c nude mice treated with indicated concentrations of SJF-0628, QDx3. Tumors were harvested 8 hours after last treatment. **c,** Average mice body weight in SK-MEL-246 efficacy study.

## References

1. Zhang, W. & Liu, H. T. MAPK signal pathways in the regulation of cell proliferation in mammalian cells. Cell Research 12, 9–18, doi:10.1038/sj.cr.7290105 (2002).

2. Roberts, P. J. & Der, C. J. Targeting the Raf-MEK-ERK mitogen-activated protein kinase cascade for the treatment of cancer. Oncogene 26, 3291–3310, doi:10.1038/sj.onc.1210422 (2007).

3. Kung, J. E. & Jura, N. Structural Basis for the Non-catalytic Functions of Protein Kinases. Structure 24, 7–24, doi:10.1016/j.str.2015.10.020 (2016).

4. Haling, Jacob R. et al. Structure of the BRAF-MEK Complex Reveals a Kinase Activity Independent Role for BRAF in MAPK Signaling. Cancer Cell 26, 402–413, doi:https://doi.org/10.1016/j.ccr.2014.07.007 (2014).

5. Garnett, M. J., Rana, S., Paterson, H., Barford, D. & Marais, R. Wild-type and mutant B-RAF activate C-RAF through distinct mechanisms involving heterodimerization. Mol Cell 20, 963–969, doi:10.1016/j.molcel.2005.10.022 (2005).

6. Flemming, A. Targeting mutant BRAF in metastatic melanoma. Nature Reviews Drug Discovery 9, 841–841, doi:10.1038/nrd3304 (2010).

7. Barras, D. BRAF Mutation in Colorectal Cancer: An Update. Biomark Cancer 7, 9–12, doi:10.4137/BIC.S25248 (2015).

8. Nguyen-Ngoc et al. - 2015 - BRAF Alterations as Therapeutic Targets in Non–Small-Cell Lung Cancer.pdf.

9. BRAF V600E and Hairy-Cell Leukemia. Cancer Discovery 1, 96–96, doi:10.1158/2159-8290.Cd-rw042711-06 (2011).

10. Holderfield, M., Deuker, M. M., McCormick, F. & McMahon, M. Targeting RAF kinases for cancer therapy: BRAF-mutated melanoma and beyond. Nat Rev Cancer 14, 455–467, doi:10.1038/nrc3760 (2014).

11. Dankner, M., Rose, A. A. N., Rajkumar, S., Siegel, P. M. & Watson, I. R. Classifying BRAF alterations in cancer: new rational therapeutic strategies for actionable mutations. Oncogene 37, 3183–3199, doi:10.1038/s41388-018-0171-x (2018).

12. Forbes, S. A. et al. COSMIC: mining complete cancer genomes in the Catalogue of Somatic Mutations in Cancer. Nucleic Acids Research 39, D945–D950, doi:10.1093/nar/gkq929 (2010).

13. Pratilas, C. A. et al. ^V600E^BRAF is associated with disabled feedback inhibition of RAF–MEK signaling and elevated transcriptional output of the pathway. Proceedings of the National Academy of Sciences 106, 4519–4524, doi:10.1073/pnas.0900780106 (2009).

14. Yao, Z. et al. BRAF Mutants Evade ERK-Dependent Feedback by Different Mechanisms that Determine Their Sensitivity to Pharmacologic Inhibition. Cancer cell 28, 370–383, doi:10.1016/j.ccell.2015.08.001 (2015).

15. Yao, Z. et al. Tumours with class 3 BRAF mutants are sensitive to the inhibition of activated RAS. Nature 548, 234, doi:10.1038/nature23291, https://www.nature.com/articles/nature23291#supplementary-information (2017).

16. Heidorn, S. J. et al. Kinase-dead BRAF and oncogenic RAS cooperate to drive tumor progression through CRAF. Cell 140, 209–221, doi:10.1016/j.cell.2009.12.040 (2010).

17. Wan, P. T. C. et al. Mechanism of Activation of the RAF-ERK Signaling Pathway by Oncogenic Mutations of B-RAF. Cell 116, 855–867, doi:https://doi.org/10.1016/S0092-8674(04)00215-6 (2004).

18. Shelledy, P. L. & Roman, P. B. D. Vemurafenib: First-in-Class BRAF-Mutated Inhibitor for the Treatment of Unresectable or Metastatic Melanoma. Journal of the Advanced Practitioner in Oncology 6, doi:10.6004/jadpro.2015.6.4.6 (2015).

19. Sosman, J. A. et al. Survival in BRAF V600–Mutant Advanced Melanoma Treated with Vemurafenib. New England Journal of Medicine 366, 707–714, doi:10.1056/NEJMoa1112302 (2012).

20. Salami, J. & Crews, C. M. Waste disposal-An attractive strategy for cancer therapy. Science 355, 1163–1167, doi:10.1126/science.aam7340 (2017).

21. Sakamoto, K. M. et al. Protacs: chimeric molecules that target proteins to the Skp1-Cullin-F box complex for ubiquitination and degradation. Proc Natl Acad Sci U S A 98, 8554–8559, doi:10.1073/pnas.141230798 (2001).

22. Bondeson, D. P. et al. Catalytic in vivo protein knockdown by small-molecule PROTACs. Nat Chem Biol 11, 611–617, doi:10.1038/nchembio.1858 (2015).

23. Bondeson, D. P. et al. Lessons in PROTAC Design from Selective Degradation with a Promiscuous Warhead. Cell Chem Biol 25, 78–87 e75, doi:10.1016/j.chembiol.2017.09.010 (2018).

24. Burslem, G. M. et al. Targeting BCR-ABL1 in Chronic Myeloid Leukemia by PROTAC-Mediated Targeted Protein Degradation. Cancer Res 79, 4744–4753, doi:10.1158/0008-5472.CAN-19-1236 (2019).

25. Burslem, G. M. et al. The Advantages of Targeted Protein Degradation Over Inhibition: An RTK Case Study. Cell Chem Biol 25, 67–77 e63, doi:10.1016/j.chembiol.2017.09.009 (2018).

26. Lai, A. C. et al. Modular PROTAC Design for the Degradation of Oncogenic BCR-ABL. Angew Chem Int Ed Engl 55, 807–810, doi:10.1002/anie.201507634 (2016).

27. Salami, J. et al. Androgen receptor degradation by the proteolysis-targeting chimera ARCC-4 outperforms enzalutamide in cellular models of prostate cancer drug resistance. Commun Biol 1, 100, doi:10.1038/s42003-018-0105-8 (2018).

28. Sun, B. et al. BET protein proteolysis targeting chimera (PROTAC) exerts potent lethal activity against mantle cell lymphoma cells. Leukemia 32, 343–352, doi:10.1038/leu.2017.207 (2018).

29. Hines, J., Lartigue, S., Dong, H., Qian, Y. & Crews, C. M. MDM2-Recruiting PROTAC Offers Superior, Synergistic Antiproliferative Activity via Simultaneous Degradation of BRD4 and Stabilization of p53. Cancer Res 79, 251–262, doi:10.1158/0008-5472.CAN-18-2918 (2019).

30. Lu, J. et al. Hijacking the E3 Ubiquitin Ligase Cereblon to Efficiently Target BRD4. Chem Biol 22, 755–763, doi:10.1016/j.chembiol.2015.05.009 (2015).

31. Smith, B. E. et al. Differential PROTAC substrate specificity dictated by orientation of recruited E3 ligase. Nat Commun 10, 131, doi:10.1038/s41467-018-08027-7 (2019).

32. Buhimschi, A. D. et al. Targeting the C481S Ibrutinib-Resistance Mutation in Bruton’s Tyrosine Kinase Using PROTAC-Mediated Degradation. Biochemistry 57, 3564–3575, doi:10.1021/acs.biochem.8b00391 (2018).

33. Karoulia, Z., Gavathiotis, E. & Poulikakos, P. I. New perspectives for targeting RAF kinase in human cancer. Nat Rev Cancer 17, 676–691, doi:10.1038/nrc.2017.79 (2017).

34. Bollag, G. et al. Clinical efficacy of a RAF inhibitor needs broad target blockade in BRAF-mutant melanoma. Nature 467, 596–599, doi:10.1038/nature09454 (2010).

35. Van Molle, I. et al. Dissecting fragment-based lead discovery at the von Hippel-Lindau protein:hypoxia inducible factor 1alpha protein-protein interface. Chem Biol 19, 1300–1312, doi:10.1016/j.chembiol.2012.08.015 (2012).

36. Meng, L. et al. Epoxomicin, a potent and selective proteasome inhibitor, exhibits in vivo antiinflammatory activity. Proceedings of the National Academy of Sciences 96, 10403–10408, doi:10.1073/pnas.96.18.10403 (1999).

37. Soucy, T. A. et al. An inhibitor of NEDD8-activating enzyme as a new approach to treat cancer. Nature 458, 732–736, doi:10.1038/nature07884 (2009).

38. Poulikakos, P. I. et al. RAF inhibitor resistance is mediated by dimerization of aberrantly spliced BRAF(V600E). Nature 480, 387–390, doi:10.1038/nature10662 (2011).

39. Lin, L. et al. Mapping the molecular determinants of BRAF oncogene dependence in human lung cancer. Proceedings of the National Academy of Sciences of the United States of America 111, E748–E757, doi:10.1073/pnas.1320956111 (2014).

40. Yao, Z. et al. RAF inhibitor PLX8394 selectively disrupts BRAF dimers and RAS-independent BRAF-mutant-driven signaling. Nat Med 25, 284–291, doi:10.1038/s41591-018-0274-5 (2019).

41. Sen, B. et al. Kinase-impaired BRAF mutations in lung cancer confer sensitivity to dasatinib. Sci Transl Med 4, 136ra170, doi:10.1126/scitranslmed.3003513 (2012).

42. Lavoie, H. & Therrien, M. Regulation of RAF protein kinases in ERK signalling. Nat Rev Mol Cell Biol 16, 281–298, doi:10.1038/nrm3979 (2015).

43. Cutler, R. E., Jr., Stephens, R. M., Saracino, M. R. & Morrison, D. K. Autoregulation of the Raf-1 serine/threonine kinase. Proc Natl Acad Sci U S A 95, 9214–9219, doi:10.1073/pnas.95.16.9214 (1998).

44. Tran, N. H., Wu, X. & Frost, J. A. B-Raf and Raf-1 are regulated by distinct autoregulatory mechanisms. J Biol Chem 280, 16244–16253, doi:10.1074/jbc.M501185200 (2005).

45. Han, X.-R. et al. Discovery of Selective Small Molecule Degraders of BRAF-V600E. Journal of Medicinal Chemistry, doi:10.1021/acs.jmedchem.9b02083 (2020).

46. Diedrich, B. et al. Discrete cytosolic macromolecular BRAF complexes exhibit distinct activities and composition. EMBO J 36, 646–663, doi:10.15252/embj.201694732 (2017).

47. Ritt, D. A., Monson, D. M., Specht, S. I. & Morrison, D. K. Impact of feedback phosphorylation and Raf heterodimerization on normal and mutant B-Raf signaling. Molecular and cellular biology 30, 806–819, doi:10.1128/MCB.00569-09 (2010).

48. Lito, P. et al. Relief of profound feedback inhibition of mitogenic signaling by RAF inhibitors attenuates their activity in BRAFV600E melanomas. Cancer cell 22, 668–682, doi:10.1016/j.ccr.2012.10.009 (2012).

49. Hatzivassiliou, G. et al. Mechanism of MEK inhibition determines efficacy in mutant KRAS-versus BRAF-driven cancers. Nature 501, 232–236, doi:10.1038/nature12441 (2013).

50. Lito, P. et al. Disruption of CRAF-mediated MEK activation is required for effective MEK inhibition in KRAS mutant tumors. Cancer Cell 25, 697–710, doi:10.1016/j.ccr.2014.03.011 (2014).

51. Morris, E. J. et al. Discovery of a novel ERK inhibitor with activity in models of acquired resistance to BRAF and MEK inhibitors. Cancer Discov 3, 742–750, doi:10.1158/2159-8290.Cd-13-0070 (2013).

52. Miura, G. MEK it work. Nature Chemical Biology 9, 601–601, doi:10.1038/nchembio.1350 (2013).

53. Röck, R. et al. BRAF inhibitors promote intermediate BRAF(V600E) conformations and binary interactions with activated RAS. Science Advances 5, eaav8463, doi:10.1126/sciadv.aav8463 (2019).

