## Supplement for "Mutant-selective Degradation by BRAF-targeting PROTACs": Supplementary files.pdf

#### Figure 1

**Vemurafenib-based PROTAC SJF-0628 potently, selectively, and efficiently induces degradation of mutant BRAF**

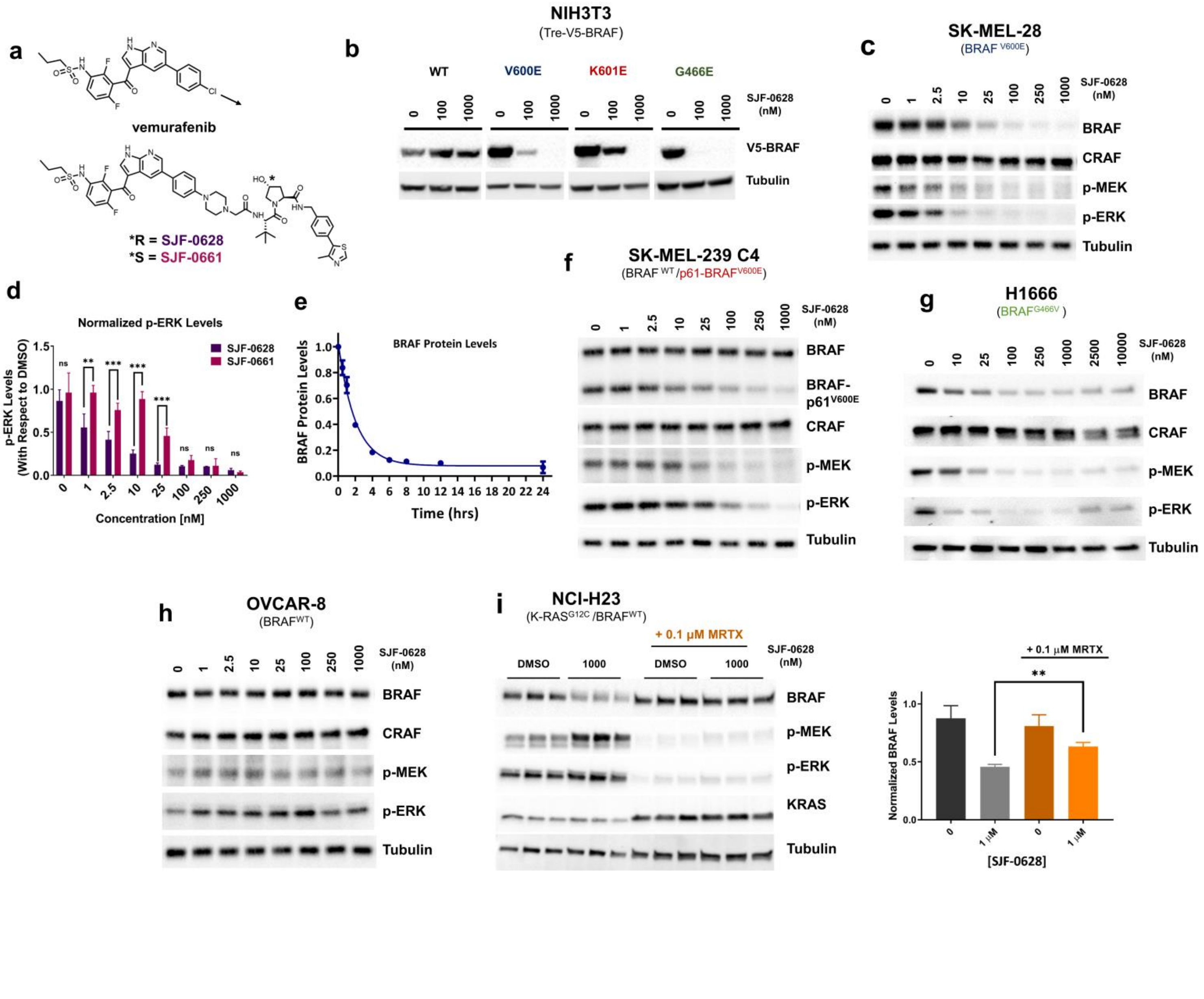

**a**, Chemical structure of **vemurafenib** and BRAF targeting PROTAC, **SJF-0628**, and its epimer, **SJF-0661.** SJF-0628 is composed of vemurafenib, a short piperazine-based linker, and a VHL recruiting ligand. **SJF-0661** has an identical warhead and linker as SJF-0628 but contains an inverted hydroxyl group in the VHL ligand and is therefore unable to engage VHL to induce ubiquitination. **b**, Inducible NIH3T3 cells expressing indicated V5-BRAF constructs (doxycycline 100-200 ng/mL, 24 hours) treated with increasing amounts of SJF-0628. **c**, SK-MEL-28 cells (homozygous $BRAF^{V600E}$) treated with indicated amounts of SJF-0628 induced BRAF degradation and suppression of MEK and ERK phosphorylation. **d**, Quantitation of ERK inhibition in SK-MEL-28 cells treated with SJF-0628 or SJF-0661 (mean ± s.d., n=3) *** $P$ value < 0.001. **e,** Quantitation of SJF-0628 treatment time course (100 nM) at indicated times in SK-MEL-28 cells shows maximal degradation within 4 hours (mean ± s.d., n=2). **f**, SJF-0628 induces selective degradation of p61-$BRAF^{V600E}$ mutant and inhibits MEK and ERK phosphorylation but spares $BRAF^{WT}$ and CRAF in SK-MEL-239-C4 cells. **g,** H1666 (heterozygous $BRAF^{G466V}$) treated with SJF-0628 shows BRAF degradation, but incomplete suppression of ERK signaling. **h**, $BRAF^{WT}$ is spared by SJF-0628 in OVCAR-8 cells but induces slight activation of ERK phosphorylation. **i**, Covalent inhibition of $KRAS^{G12C}$ by MRTX849 in H23 cells hinders PROTAC induced $BRAF^{WT}$ degradation ((mean ± s.d.,  n=3, ** P value < 0.01). $P$ value calculated by unpaired t-test.

**Figure 2**

**BRAF$^{WT}$ is unable to form a PROTAC-induced ternary complex in cells and thus not degraded**

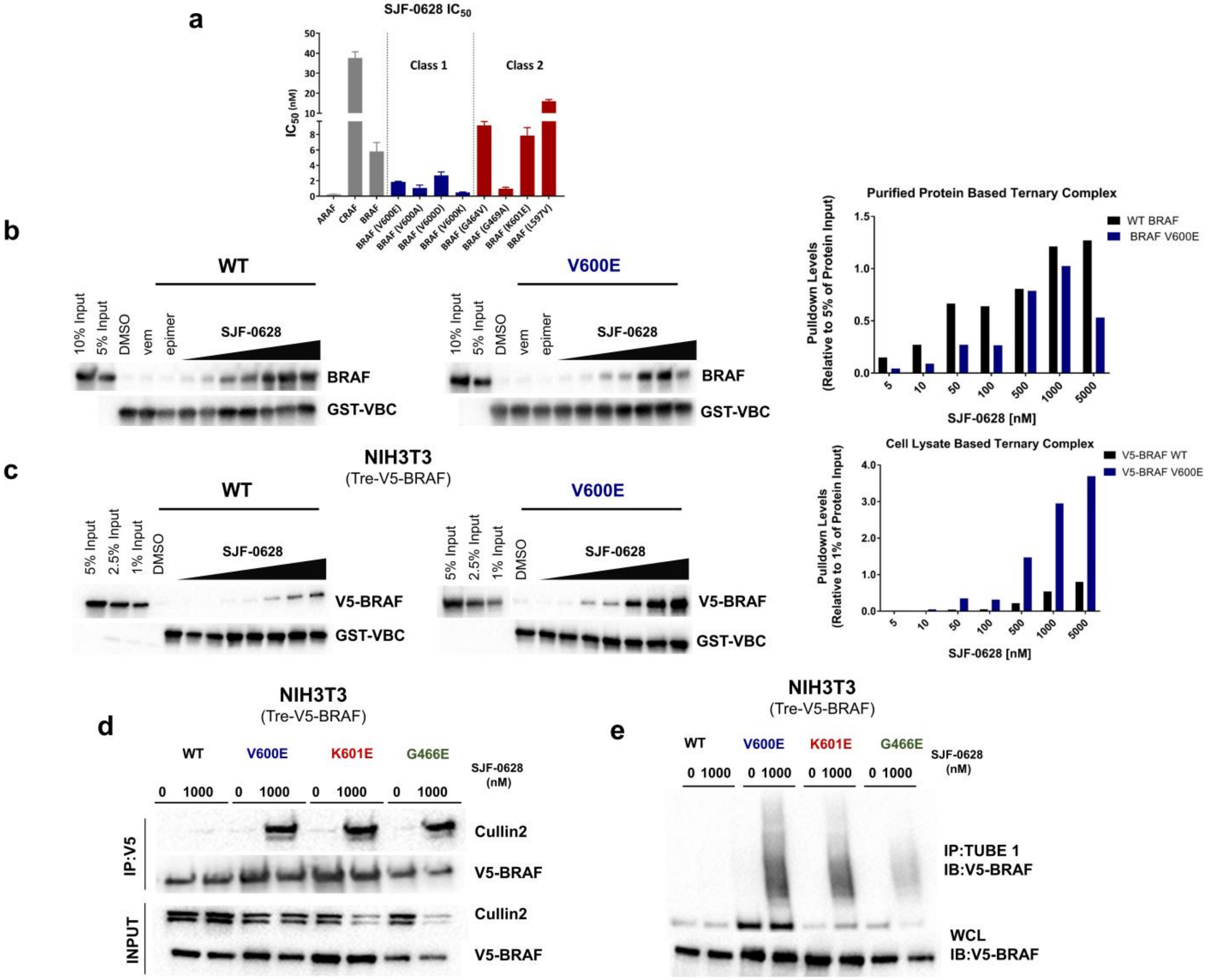


**a**, IC$_{50}$ values of radiolabeled kinase assay for WT RAF and Class 1 and 2 BRAF mutants (mean ± s.d., n=2). Plotted values shown in Table 1. **b**, Purified protein ternary complex assay. GST-VBC (VHL, Elongin B, Elongin C) is immobilized on glutathione beads and incubated with

DMSO, vemurafenib (500 nM), SJF-0661 (500 nM) or increasing concentrations of SJF-0628 and purified full length-BRAF to observe VBC:PROTAC:BRAF ternary complex. Quantified with respect to 5% input. **c**, Cell lysate based ternary complex assay (as described in **b**) but using NIH3T3 cell lysates (doxycycline 800 ng/mL) containing V5-BRAF$^{WT}$ or V5-BRAF$^{V600E}$ as input. Quantified with respect to 1% input. **d**, NIH3T3 cells expressing indicated V5-BRAF treated with DMSO or 1µM SJF-0628 for 1-hour followed by immunoprecipitation of V5-BRAF. **e,** Tandem Ubiquitin Binding Entities 1 (TUBE1) pull down of tetra-ubiquitinated proteins in NIH3T3 cells expressing indicated V5-BRAF after 1-hour treatment with vehicle and SJF-0628. Immunoblotted for V5-BRAF.

### Figure 3

**MEK inhibitors that activate BRAF also sensitize BRAF to PROTAC-induced ubiquitination and degradation.**

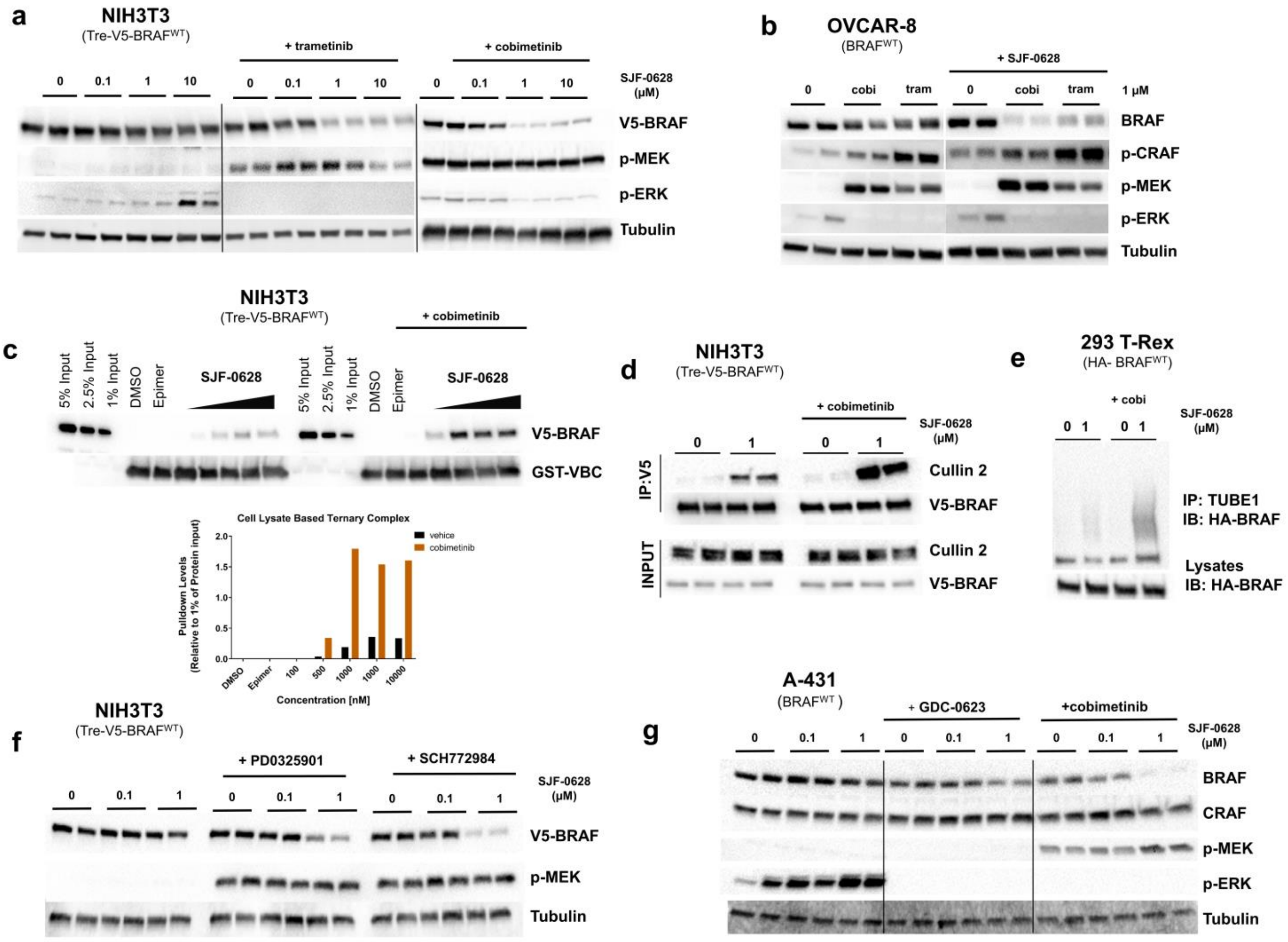


**a**, NIH3T3 cells with trametinib (1µM, 5 hours) or cobimetinib (500 nM, 3 hours) pre-treatment subsequently treated with increasing amounts of SJF-0628 (20 hours) promote degradation of BRAF$^{WT}$ and show a marked increase in p-MEK. **b**, OVCAR8 cells pre-treated with cobimetinib and trametinib (1µM, 2 hours) promote MEK and CRAF phosphorylation as well as BRAF degradation in the presence of SJF-0628. **c**, Cell lysate-based ternary complex assay shown in 2**c**

but using NIH3T3 lysates expressing $BRAF^{WT}$ and pre-treated with DMSO or 1µM cobimetinib for 3 hours. Cobimetinib pre-treatment promotes ternary complex formation. **d**, V5-BRAF immunoprecipitation in NIH3T3 cells pre-treated with 1µM of cobimetinib (2 hours) followed by treatment of SJF-0628 for 2.5 hours. **e**, TUBE1 pulldown in 293 T-Rex cells stably expressing HA- $BRAF^{WT}$ treated with cobimetinib (cobi) (2 hours, 1µM) and subsequently treated with SJF-0628 (2 hours). **f,** NIH3T3 cells pre-treated with 1µM PD0325901 (MEK inhibitor) or SCH772984 (ERK inhibitor) for 3 hours followed by treatment with indicated amount of SJF-0628 for 20 hours. **g**, A431 cells pre-treated with GDC-0623 and cobimetinib (500nM for 3 hours) then treated with SJF-0628 for 20 hours.

### Figure 4

**SJF-0628 outperforms vemurafenib in inhibiting growth of cell lines expressing mutant BRAF**

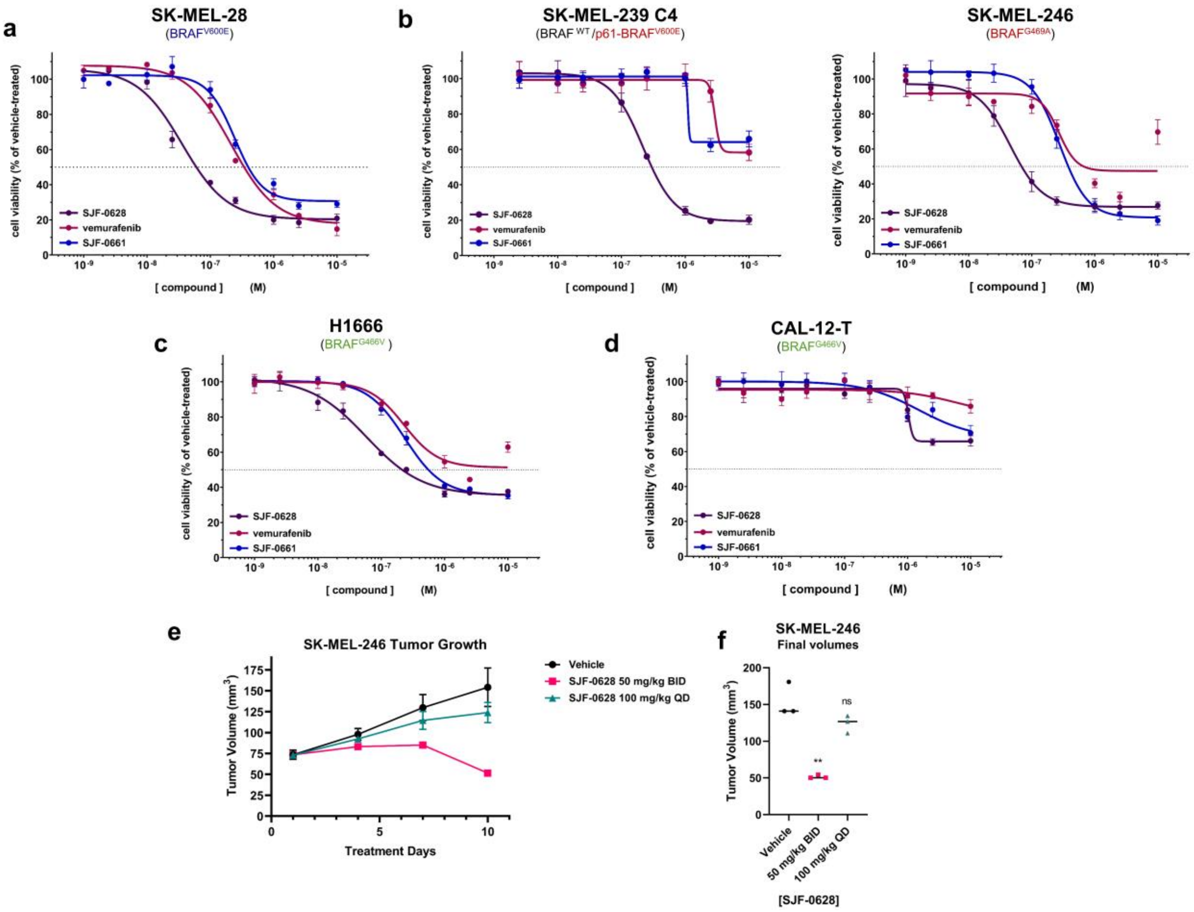


**a,** Cell proliferation assay in SK-MEL-28 cells treated with increasing amounts of vemurafenib, SJF-0628, or SJF-0661 for 3 days (mean ± s.d., n=3). $EC_{50}$ = 215 ± 1.09 nM, 37 ± 1.2 nM, and 243 ± 1.09 nM respectively. **b,** Cell proliferation assay in vemurafenib resistant SK-MEL-239-C4 cells treated with increasing amounts vemurafenib, SJF-0628, or SJF-0661 for 5 days (mean ± s.d., n=3). **c,** Cell proliferation assay in SK-MEL-246 (Class 2) cells treated with increasing amounts vemurafenib, SJF-0628, or SJF-0661 for 5 days (mean ± s.d., n=3) **d,** SJF-0628 EC50=218 nM±1.06 c,H1666 cells treated with SJF-0628, vemurafenib, or SJF-0661 for 5 days (mean ± s.d., n=3). **d,** Treatment of CAL-12-T cells with vemurafenib, SJF-0628, or SJF-0661 for 5 days shows minimal effect on cell viability (mean ± s.d., n=3). **e,** Results of an efficacy study in SK-MEL-246 tumor xenografts implanted in female athymic mice showing tumor regression with 50 mg/kg IP twice daily. **f,** Scatter plot result of final volumes (** *P* value < 0.01). *P* value calculated by unpaired t-test.

<u>**Extended Data Figure 1**</u>

**Vemurafenib based PROTAC, SJF-0628, induces mutant selective degradation of BRAF**

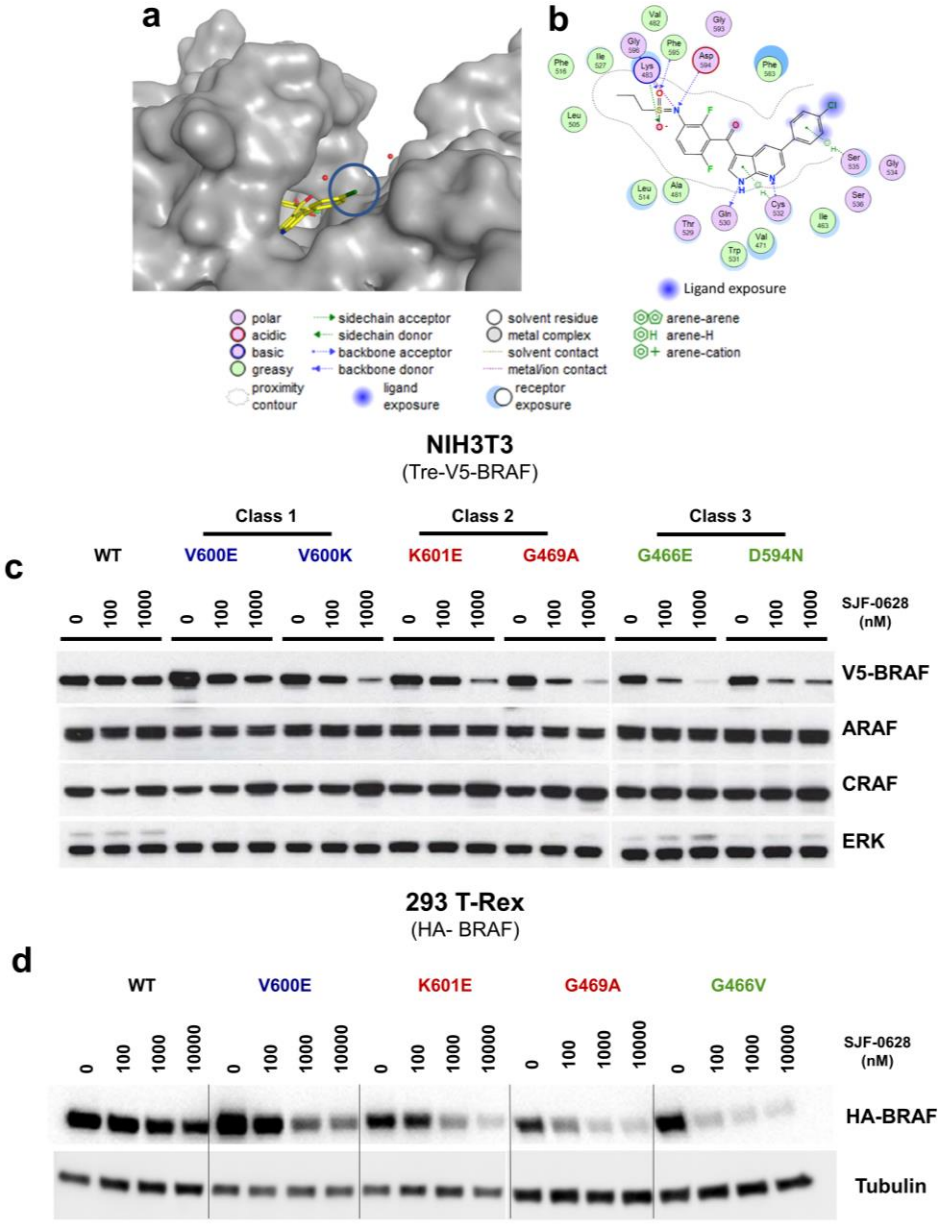


**a,** Crystal structure of BRAF$^{V600E}$ in complex with vemurafenib (PDB: 3OG7) **b**, Ligand interactions diagram showing important BRAF: vemurafenib interactions and solvent exposure. **c**, Inducible NIH3T3 cells expressing indicated V5-BRAF (doxycycline 500 ng/mL, 24 hours) treated with increasing amounts of SJF-0628 for 24 hours. **d**, 293 T-Rex cells (doxycycline 20 ng/mL, 24 hours) expressing indicated BRAF isoforms treated with increasing concentrations of SJF-0628 shows mutant selective degradation.

### Extended Data Figure 2

**SJF-0628 induces a sustained and efficient degradation of BRAF$^{V600E}$ via the proteasome**

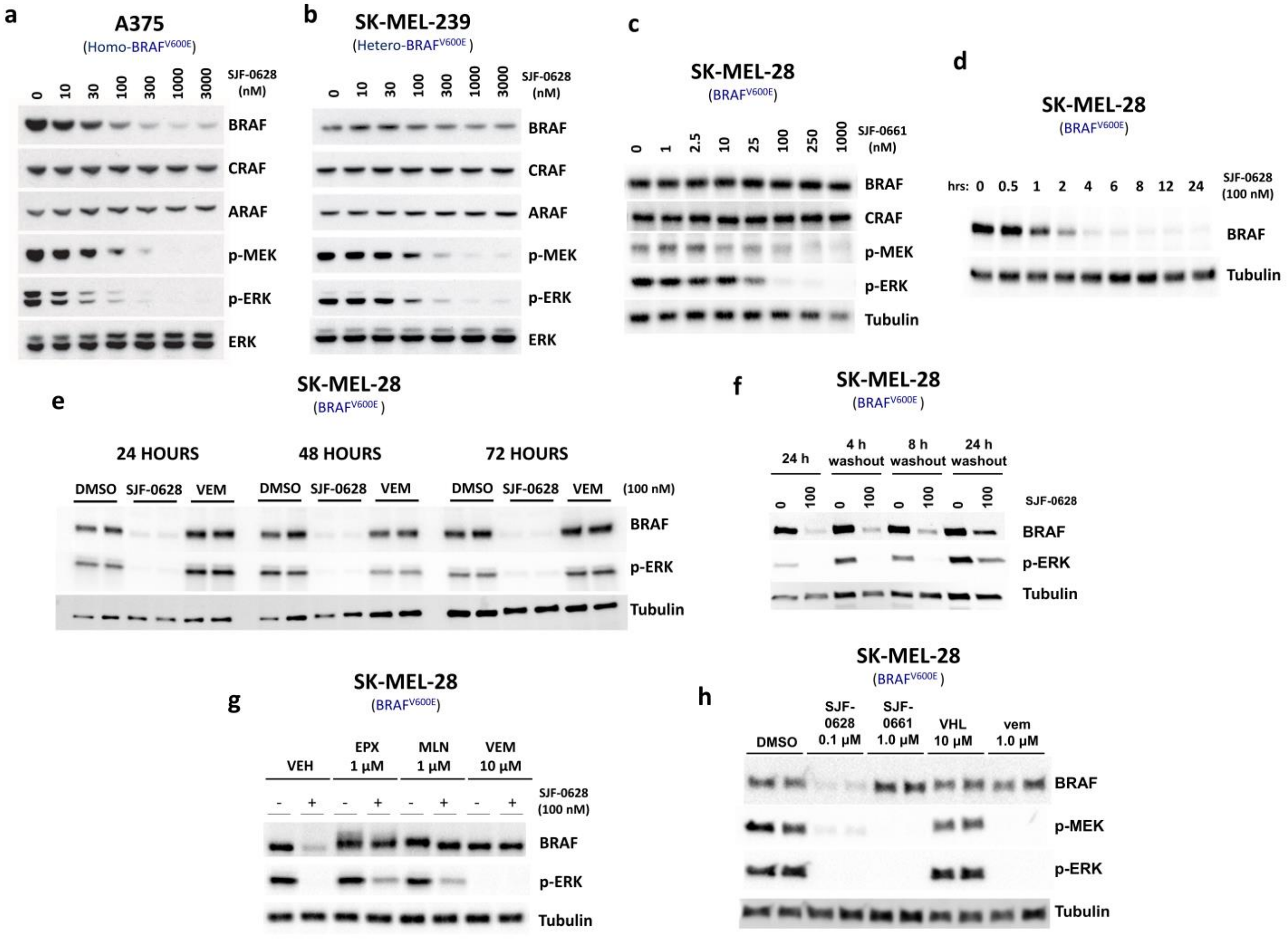

**a**, Treatment of A375 (homozygous BRAF$^{V600E}$) cells with SJF-0628 shows BRAF$^{V600E}$ degradation and inhibition of MEK and ERK phosphorylation. **b**, Treatment of SK-MEL-239 (heterozygous BRAF$^{V600E}$) cells with SJF-0628 shows minimal BRAF degradation but marked inhibition of MEK and ERK phosphorylation. **c**,SK-MEL-28 cells treated with negative control epimer, SJF-0661, does not affect BRAF levels but inhibits ERK signaling. **d**, Representative immunoblot of SJF-0628 time course (100 nM) at indicated times in SK-MEL-28 cells (plotted in Fig. 1e**)** shows maximal degradation within 4 hours. **e**, Treatment of SK-MEL-28 cells with 100 nM of SJF-0628, and vemurafenib for indicated times shows sustained degradation and inhibition of MAPK signaling. **f**, SK-MEL-28 cells were treated with 100 nM for 24 hours, washed 3 times with DPBS and replenished with fresh media. Cells were then lysed either 4, 8 or 24 hours after media removal to determine level of BRAF and MAPK recovery. **g**, SK-MEL-28 cells treated with a proteasome inhibitor (EPX = epoxomicin), a neddylation inhibitor (MLN = MLN4924), or excess vemurafenib (VEM) for 2 hours, then subsequently treated with DMSO or PROTAC for 8 hours. **h**, SK-MEL-28 cells treated with indicated compound for 6 hours. VHL ligand alone does not cause MAPK inhibition.

### Extended Data Figure 3

**SJF-0628 induces degradation of acquired and intrinsic vemurafenib-resistant BRAF mutants**

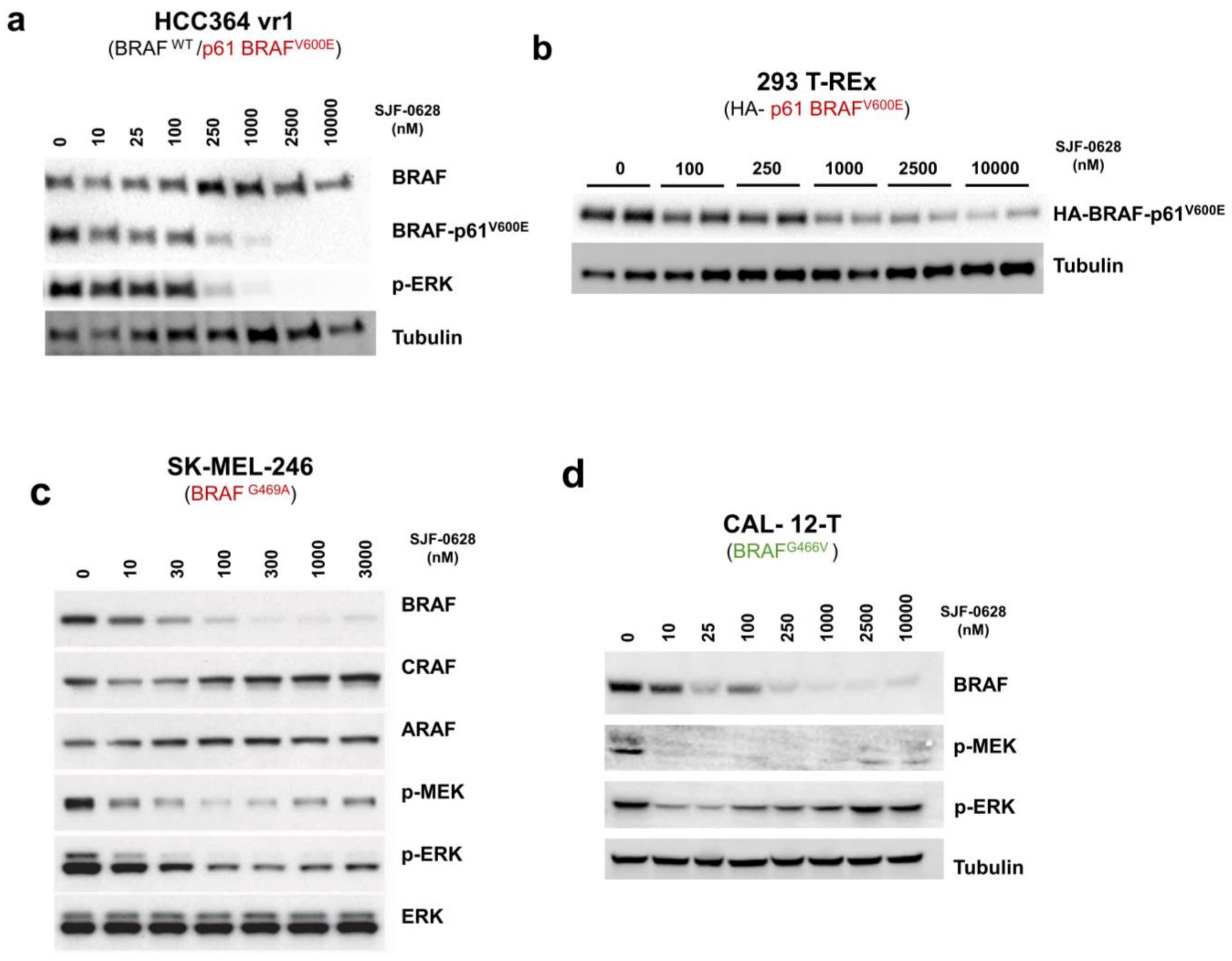

**a**, HCC364 vr1 ($BRAF^{WT}$, p61-$BRAF^{V600E}$) selectively induces degradation of p61-$BRAF^{V600E}$ and spares $BRAF^{WT}$. **b**, SJF-0628 treatment in 293 T-Rex cells expressing p61-$BRAF^{V600E}$ shows dose dependent decrease in HA-$p61^{V600E}$ protein levels. **c**, SK-MEL-246 (Class 2, BRAF $^{G469A}$) cells treated with increasing amount of SJF-0628 shows degradation of BRAF and inhibition of ERK signaling. **d**, CAL-12-T cells (homozygous $BRAF^{G466V}$) treated with SJF-0628 shows BRAF degradation, but incomplete suppression of ERK signaling.

### Extended Data Figure 4

**BRAF$^{WT}$ is sensitized to PROTAC induced degradation in the presence of upstream drivers**

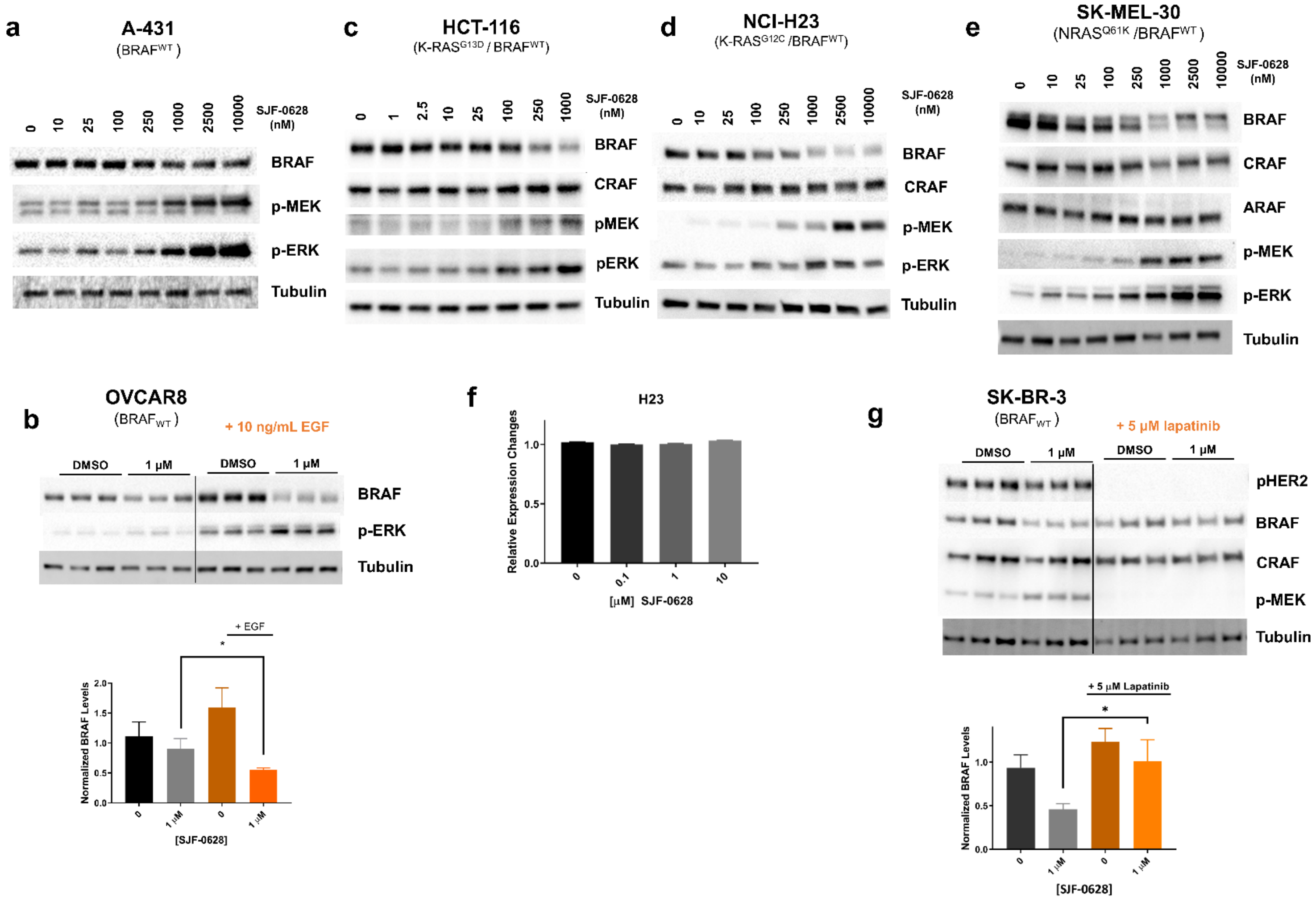


**a**, Treatment of A-431 cells (HER1 amplification) which expresses BRAF$^{WT}$ and RAS$^{WT}$ treated with increasing amounts of SJF-0628 show some degradation of BRAF$^{WT}$(~30%). **b**, Serum-starved OVCAR8 cells stimulated with 10 ng/mL of EGF promotes SJF-0628 induced degradation of $BRAF^{WT}$ **c**-**e**, SJF-0628 treatment in HCT116, H23, and SK-MEL-30 cells bearing a RAS mutation shows 40%-60% degradation of $BRAF^{WT}$ at high concentrations and paradoxical activation of MAPK signaling. **f,** Quantitative real time PCR of H23 cells treated with SJF-0628 for 20 hours (mean ± s.d.,n=3). **g**, Lapatinib treatment in SKBR3 cells hinders PROTAC induced degradation ( *$P$ value $<0.05$). $P$ value calculated by unpaired t-test.

### Extended Data Figure 5

**Vemurafenib based PROTACs spare BRAF$^{WT}$ despite linker length and composition**

a

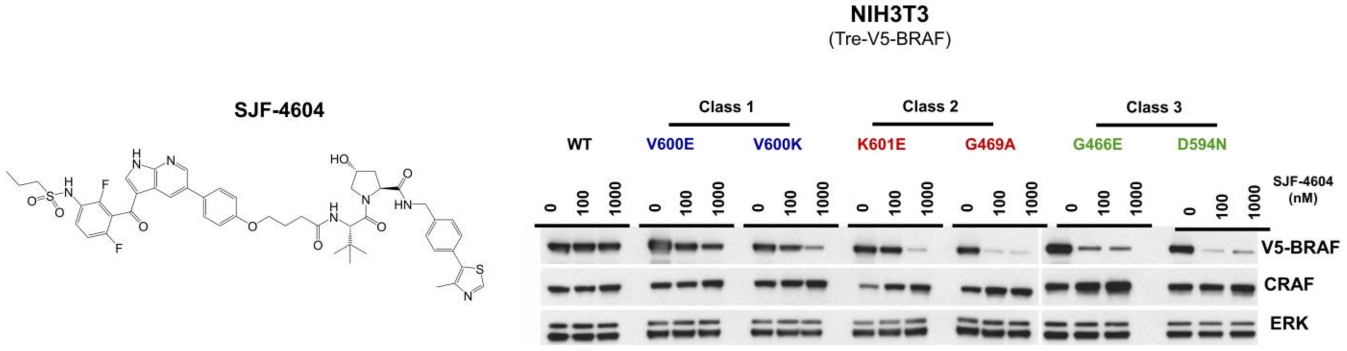


b

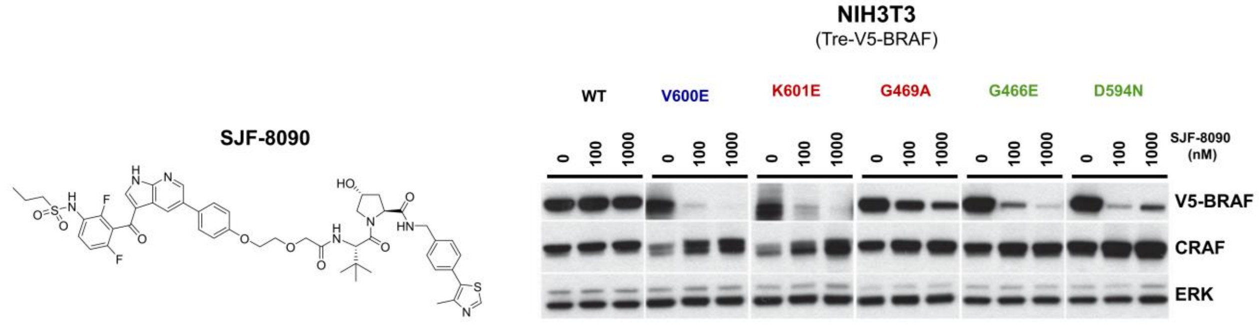


**a-b**, Structures and results of vemurafenib based PROTACs SJF-4604 and SJF-8090 treatment in inducible NIH3T3 cells expressing indicated BRAF protein for 24 hours. Both PROTACs show mutant selective degradation.

**Extended Data Figure 6**

**All three mutant BRAF classes bind SJF-0628 and form a stable ternary complex which promotes its ubiquitination**

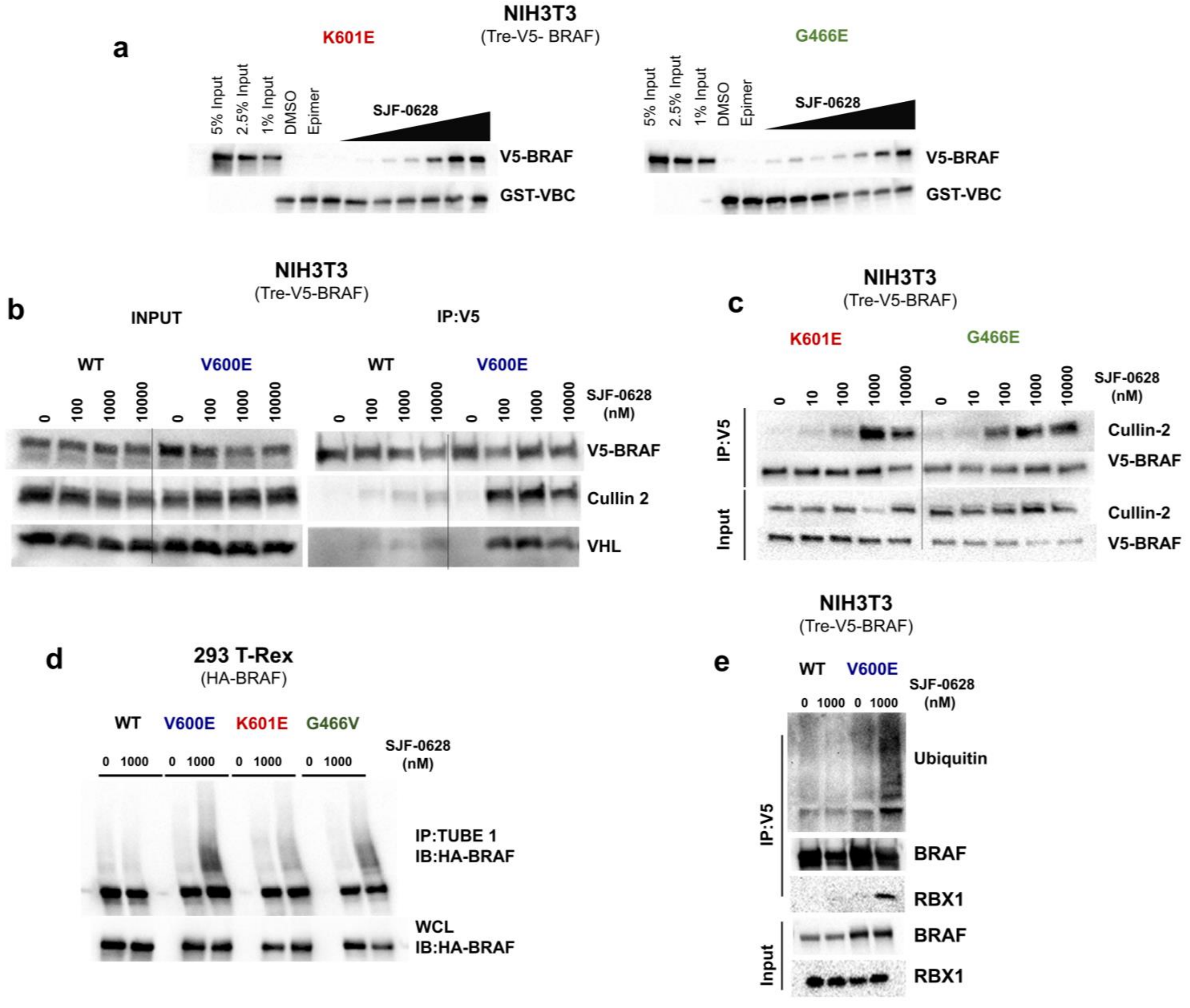


**a**, Cell lysate-based trimer assay of VBC immobilized on glutathione beads incubated with NIH3T3 cell lysates expressing $BRAF^{K601E}$ and $BRAF^{G466E}$ with vehicle, 500 nM SJF-0661, or increasing amounts of SJF-0628. **b**, V5-BRAF immunoprecipitation in NIH3T3 cells (WT and V600E) stably expressing human VHL treated with increasing concentrations SJF-0628 for 45 mins. **c,** V5-BRAF immunoprecipitation in NIH3T3 cells expressing $BRAF^{K601E}$ or $BRAF^{G466E}$ V600E treated with SJF-0628 for 1 hour. **d**, Tandem Ubiquitin Binding Entities 1 (TUBE1) Assay. Pulldown of tetra-ubiquitinated protein in 293 T-REx cells expressing indicated BRAF treated with SJF-0628 for 1 hour. Immunoblot for HA-BRAF. **e**, V5-BRAF immunoprecipitation in NIH3T3 cells expressing $BRAF^{WT}$ and $BRAF^{V600E}$ treated with SJF-0628 for 1 hour.

### Extended Data Figure 7

**Trametinib or cobimetinib pre-treatment promotes dose and time dependent degradation of BRAF$^{WT}$**

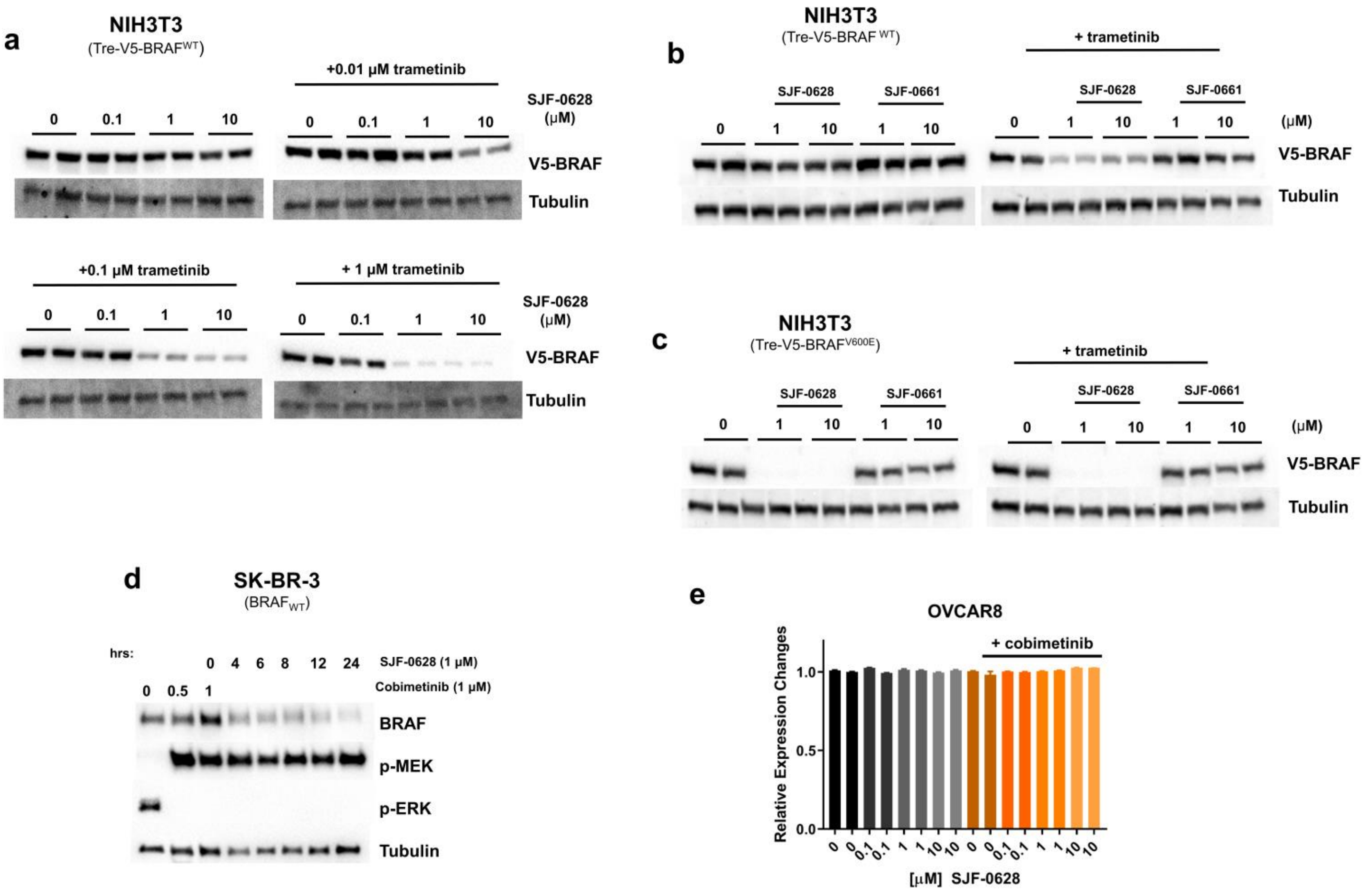


**a**, NIH3T3 cells expressing inducible V5- BRAF$^{WT}$ pre-treated with increasing concentrations of trametinib and subsequently treated with increasing concentrations of SJF-0628. **b-c**, NIH3T3 cells expressing BRAF$^{WT}$ and BRAF$^{V600E}$ pre-treated with trametinib followed by SJF-0628 or SJF-0661 treatment. **d,** Time course of SK-BR-3 cells pre-treated with 1 μM of cobimetinib for 1 hour then treated with 1 μM SJF-0628. **e**, mRNA expression changes of BRAF pre-treated with 1 μM of cobimetinib for 1 hour, then treated with 1 μM SJF-0628 for 20 hours(mean ± s.d.,n=3).

<u>**Extended Data Figure 8**</u>

**MAPK inhibitors that increase BRAF kinase activity promote SJF-0628 induced BRAF$^{WT}$ degradation**

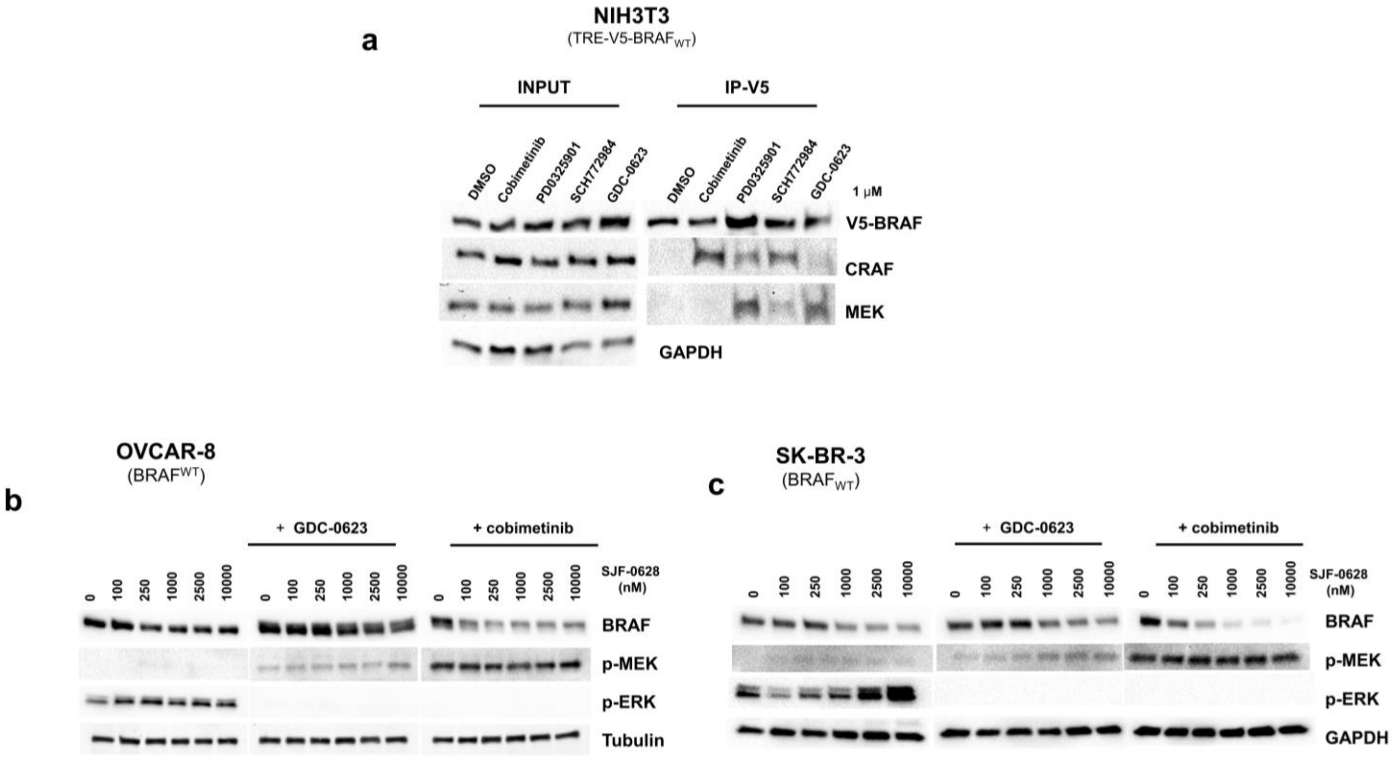


**a**, Immunoprecipitation of V5-BRAF$^{WT}$ from NIH3T3 cells treated with 1uM of the indicated MAPK pathway inhibitor. Cobimetinib treatment showed minimal BRAF association with MEK, but increased RAF dimerization while GDC-0623 showed minimal RAF dimerization, and increased BRAF: MEK association. **b-c**, OVCAR-8 cells and SK-BR-3 cells pre-treated with GDC-0623 and cobimetinib (500nM, 3 hours) then treated with SJF-0628 for 20 hours.

### Extended Data Figure 9

**SJF-0628 inhibits BRAF$^{WT}$ and BRAF$^{V600E}$ with a similar affinity and induces degradation of BRAF$^{V600E}$ *in vivo***

a

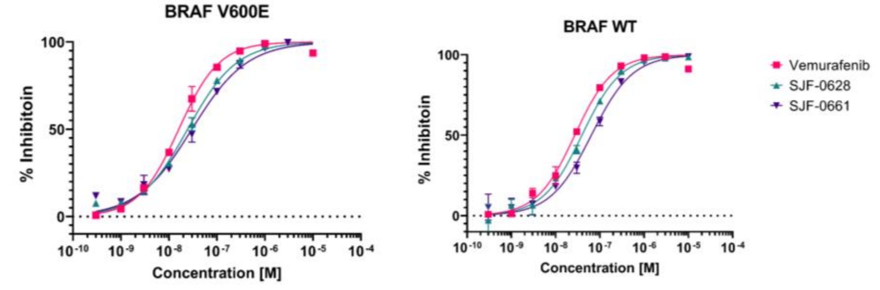


| Kinase | IC$_{50}$ (M) vemurafenib | SJF-0628 | SJF-0661 |
|---|---|---|---|
| BRAF V600E | 1.60E-08 | 2.44E-08 | 3.00E-08 |
| BRAF WT | 2.70E-08 | 3.92E-08 | 6.40E-08 |

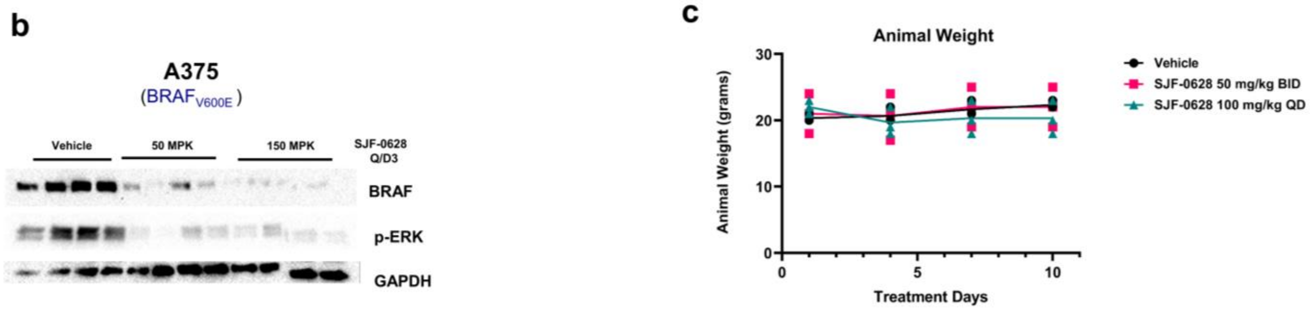


**a**, BRAF$^{WT}$ and BRAF$^{V600E}$ binding affinity and curves for ELISA kinase inhibition assay with SJF-0628, SJF-0661 and vemurafenib (n=2). **b,** BRAF$^{V600E}$ degradation in A375 xenograft in female Balb/c nude mice treated with indicated concentrations of SJF-0628, QDx3. Tumors were harvested 8 hours after last treatment. **c,** Average mice body weight in SK-MEL-246 efficacy study.

**Table 1**

| Kinase: | SJF-0628 IC$_{50}$ (nM) |
|---|---|
| ARAF | 0.27 |
| CRAF | 37.6 |
| BRAF | 5.80 |
| BRAF (V600E) | 1.87 |
| BRAF (V600A) | 1.06 |
| BRAF (V600D) | 2.68 |
| BRAF (V600K) | 0.49 |
| BRAF (G464V) | 9.18 |
| BRAF (G469A) | 0.98 |
| BRAF (K601E) | 7.84 |
| BRAF (L597V) | 16.0 |

**Table of IC$_{50}$ values from radio labeled kinase assay (n=2)**
